# Active hydraulics and odd elasticity of muscle fibers

**DOI:** 10.1101/2022.02.20.481216

**Authors:** Suraj Shankar, L. Mahadevan

## Abstract

Muscle is a complex hierarchically organized soft contractile engine. To understand the limits on the rate of contraction and muscle energetics, we construct a coarse-grained multiscale model that integrates over molecular details and describes muscle as an active sponge. Our analysis of existing experiments highlights the importance of spatially heterogeneous strains and local volumetric deformations in muscular contractions across species and muscle type. The minimal theoretical model shows how contractions generically induce intracellular fluid flow and power active hydraulic oscillations, which determine the limits of ultrafast muscular contractions. We further demonstrate that the viscoelastic response of muscle is naturally nonreciprocal – or odd – owing to its active and anisotropic nature. This points to an alternate mode of muscular power generation from periodic cycles in spatial strain alone, contrasting with previous descriptions based on temporal cycles. Our work suggests the need for a revised view of muscle dynamics that emphasizes the multiscale spatio-temporal origins of soft hydraulic power, with potential implications for physiology, biomechanics and locomotion.

## 1. Introduction

Muscle is the primary driver of nearly all motion across the animal kingdom. Since the pioneering work by H. E. Huxley, A. F. Huxley and others reviewed in [1, 2], much work has focused on elucidating the biochemical details of the molecular rate-limiting steps involved in the contractile machinery, i.e., the binding kinetics of actomyosin crossbridges [3, 4, 5] and calcium signaling that controls motor kinetics [6, 7]. But muscle fibers are more than molecular motors, being spatially and heirarchically organized across multiple scales [8] (Fig. 1A) with complex structural and mechanical properties. They are just as responsible for the slow movements of the sloth bear as the rapid contractions associated with high frequency sound production by rattlesnakes, fish swim-bladders and songbirds, and the flapping wings of insects, where operating frequencies can range between 10^2^ *−*10^3^ Hz [9, 10]. Such extremely fast motions naturally raise the question of the maximal rate at which muscle contracts, and the limits on its energetics. To understand these questions requires us to integrate processes across scales. On mesoscopic scales, striated muscle fibers are soft, wet and active materials composed of a dense, anisotropic and actively contracting polymeric lattice (sarcomeres forming the myofibril), bathed in cytosol. The dominant component of muscle fibers, by far, is water (0.7 *−* 0.9 volume fraction [11, 12]) and assumed to simply play a permissive role, subservient to cellular signalling and biochemical processes. But in recent years, intracellular fluid flows have increasingly been recognized for their central role in dictating cellular morphology, motility and physiology [13, 14, 15], e.g., in rapid nonmuscular movements in plants [16, 17].

**Figure 1.**
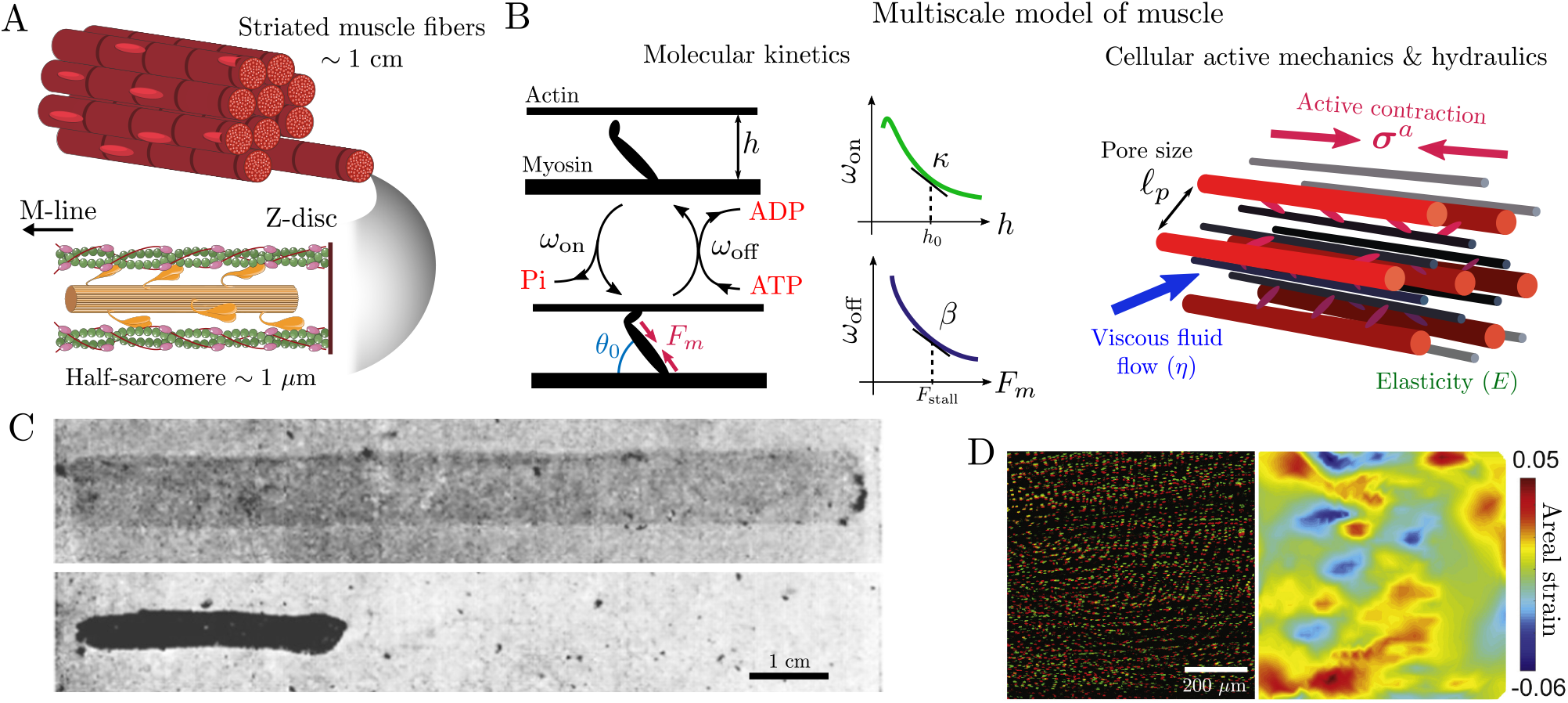
Muscle fibers are soft, wet, active and multiscale. (A) Striated muscle consists of long, multinucleated fibers that span several spatial scales, ranging from the contractile machinery arranged in periodically repeating sarcomeres (*∼* 1 *μ*m) to individual macroscopic fibers that can span 1 *−* 10 cm (figure prepared using Servier Medical Art images from Bioicons licensed under CC-BY 3.0 Unported). (B) A multiscale mathematical model for muscle fibers (Eqs. 1-5). On the molecular scale, motor heads along the myosin thick filament stochastically bind to the actin thin filament and generate a contractile force (*F*_*m*_) upon hydrolyzing ATP, with both radial and axial components for a finite crossbridge binding angle (*θ*). The kinetic rates (*ω*_on*/*off_ ) decrease with increasing inter-filament spacing (*h*) and motor force (*F*_*m*_), with *κ, β >* 0 measuring the linearized feedback response at resting, stall conditions (see SI Sec. I for details). On the mesoscale, the myofilament lattice is an anisotropic elastic network with a characteristic shear modulus *E* and pore size *𝓁*_*p*_, that is permeated by a fluid of viscosity *η*, and contracts under the action of a molecularly generated active stress (***σ***^*a*^). (C) Actomyosin threads contract by expelling water, changing both shape and size when placed in boiled muscle juice (a source of ATP) (adapted with permission from Ref. [18]).(D) In-vivo contraction of an electrically stimulated skeletal muscle (murine gastrocnemius) displaying spatial strain heterogeneities (right: heatmap of the 2D areal strain) measured using the stained nuclei as fiducial markers (green: undeformed, red: deformed; adapted with permission from Ref. [19]).

In animals, this begs a natural question: how important are spatial hydraulic effects in the dynamics of contracting muscle fibers? A coarse-grained view of muscle (as shown in Fig. 1B) suggests that muscle fibers behave as an active fluid-filled sponge. When a muscle fiber contracts, there must be relative movement of the actomyosin filament lattice relative to the ambient fluid. More specifically, intact muscle fibers cannot contract everywhere homogeneously (due to global incompressibility in the presence of an intact sarcolemma), but they can do so locally by slowly squeezing fluid through the pores of the myofilament lattice, a process that is potentially rate-limiting. The dynamical consequences of this process are typically neglected as most *in-vitro* studies focus on glycerinated (permeabilized) muscle fibers that allow free drainage of fluid and assume locally incompressible deformations [11]. That water movement may occur during muscle contraction (for various reasons including osmolyte imbalances) has been noted in old studies [20, 21], and in early experiments by Szent-Györgi [22] on extracted actomyosin threads that undergo syneresis by expelling water as they “violently contract” (Fig. 1C). More pertinently, active crossbridges are also known to produce transverse (radial) forces in addition to longitudinal ones, that can change the local volume of the sarcomere [23] and necessitate fluid redistribution. And finally, spatially nonuniform strains (Fig. 1D) have been reported during tetanic contractions of intact muscle fibres, both *ex-vivo* [24, 25] and *in-vivo* [26, 19], suggesting that local strain gradients and the accompanying pressure gradients and fluid hydraulics may all be relevant dynamically.

Recent experiments probing the rapid dynamics of the myofilament lattice using optical microscopy and small-angle X-ray scattering techniques [27, 28, 29, 30, 31, 32, 33] now allow for a quantification of some of these observations. We reanalyzed data on active oscillations of muscle fibers from various experiments performed across different muscle types and species to obtain time traces of local transverse (*∈*_*⊥*_) and longitudinal (*∈*_*zz*_) strains in the sarcomere (Fig. 2, see SI Sec. VA for details). Spontaneous contractile oscillations with frequency *ω ∼* 1 Hz can emerge in glycerinated skeletal muscle fibers (rabbit psoas, Fig. 2A-B) [27, 28]. The resulting periodic strains are not volume preserving, i.e., they are non-isochoric (Fig. 2, black line), despite the fibers being permeabilized. This is likely because the fibers are slender enough to lack radial gradients in deformation (see SI Sec. IIIB).

**Figure 2.**
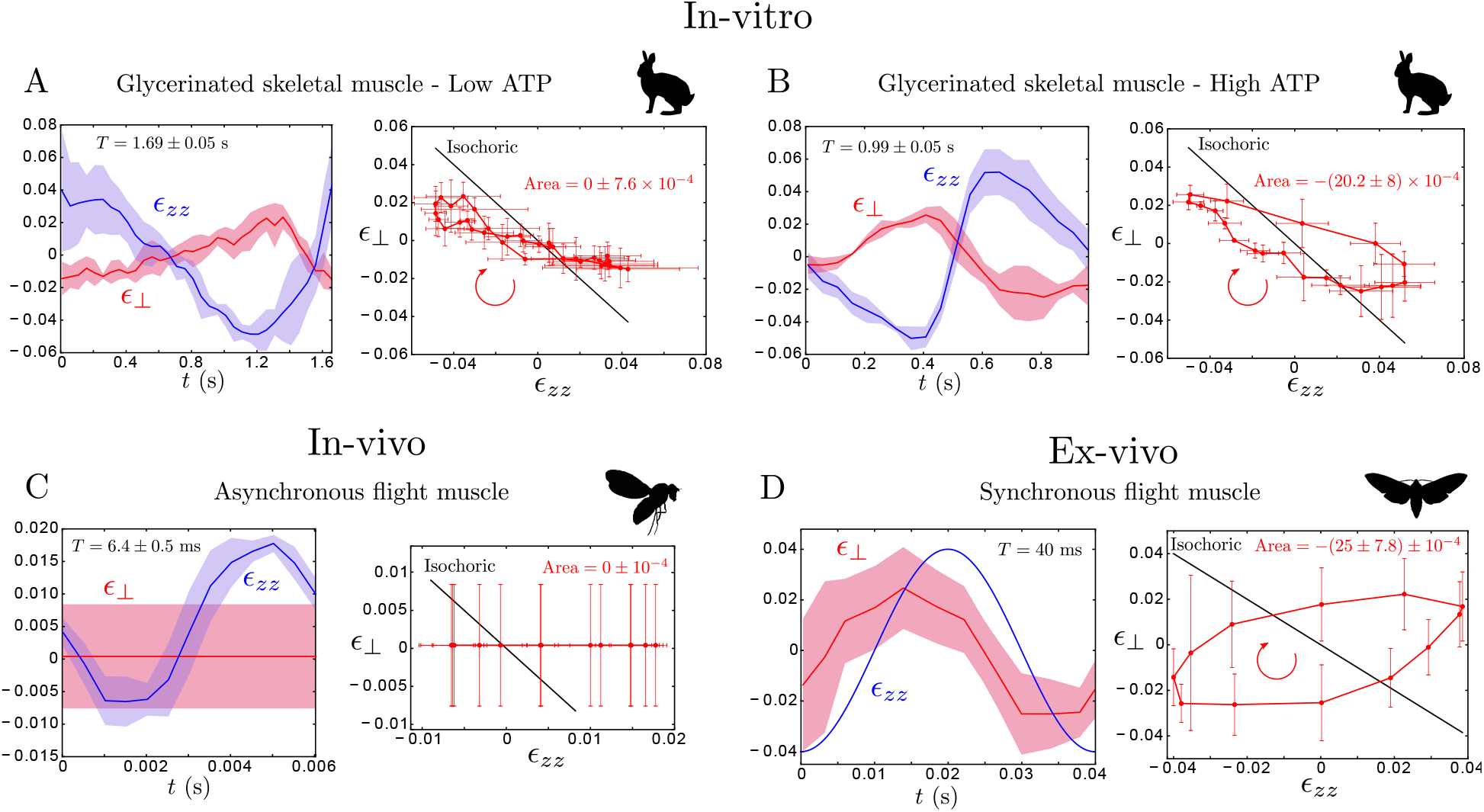
Volume changes and strain cycles are generic in self-oscillating muscle. (A-B) Skinned rabbit psoas muscle fibers exhibit spontaneous sarcomeric oscillations in-vitro under different ATP concentrations and sarcomere resting lengths [27, 28]. Deformations are generically not isochoric (*∈*_*zz*_ + *∈*_*⊥*_ = 0, black line), as a result, fluid flow is inevitable. At low ATP concentrations (A), the axial (*∈*_*zz*_) and transverse strains (*∈*_*⊥*_) oscillate nearly antiphase with frequency *ω ≈* 0.6 Hz [28], while at higher ATP concentrations (B), faster oscillations (*ω ≈* 1 Hz) develop a strain cycle [27] that encloses a significant (signed) area, see SI Sec. VA. The direction of circling the loop is shown by the red arrow. (C) In-vivo measurements of strain in asynchronous flight muscle (DLM: dorsal longitudinal) of a fruit fly in tethered flight (*ω ≈* 156 Hz) [30, 29, 31]. Axial strains plotted are obtained from high-speed optical measurements of muscle length [29] and transverse strains (computed from time-resolved X-ray diffractometry measurements with subnanometer precision, see SI Sec. VA for details) remain negligible at all times [30]. (D) Ex-vivo measurements of strain in intact synchronous flight muscle (DLM) from *Manduca sexta* (hawkmoth) [32], displaying nonisochoric strain cycles. The isolated, whole muscle is subject to oscillatory stretch (4% axial strain, *ω* = 25 Hz) and electrical stimulation (activation phase of 0.5), matching physiological in-vivo conditions. The transverse strain is estimated from time-resolved X-ray diffraction measurements of the lattice spacing. The data are presented as mean with shaded regions and error bars in A-C denoting one standard deviation from averaging over multiple periods, and a 95% confidence interval in D (sample size *N* = 1 in all). See SI Sec. VA for further details of the analysis. All data are digitally extracted and reanalyzed from Refs. [28, 27, 29, 32]. Animal silhouettes are adapted from the online open database PhyloPic and individual image credits are in the SI Sec. VD.

*In-vivo* measurements of the sarcomere geometry in intact asynchronous flight muscle of *Drosophila* [29, 30, 31] shows that, remarkably, the lattice contracts with a constant lattice spacing [30], hence *∈*_*⊥*_ *≈* 0 under natural flight conditions (wing-beat frequency *ω ≈* 156 Hz; Fig. 2C). Additionally, intact synchronous flight muscle of *Manduca sexta* displays periodic lattice dilations and contractions under physiological conditions (*ω ≈* 25 Hz; Fig. 2D), both *in-vivo* [33] and *ex-vivo* [32]. In all of the above examples (with and without a membrane), deformations of the myofilament lattice fail to preserve its local volume (*∈*_*zz*_ + *∈*_*⊥*_ *≠* 0), thereby necessitating fluid movement through the sarcomere and thus lead to spatiotemporal heterogeneities in strain.

Given the empirical evidence summarized above, how can we build a model that captures and explains the three-dimensional (3D) and volumetric deformations in muscle fibers? Here we do this and demonstrate two separate consequences of such spatial deformations in muscle fibers - (i) the dynamics of active contraction is constrained by flows within a fluid-filled fiber, and (ii) the 3D mechanical response of a muscle fiber is nonreciprocal [34, 35] allowing it to function as a soft engine using strain cycles.

Current approaches to explore multi-scale phenomena in muscle employ detailed, spatially explicit, computational models at both the cellular [36, 37, 38, 39, 40] and whole tissue level [41, 42, 43]. Other studies have recently begun addressing the role of fluid dynamics in this question by focusing on intramuscular pressure [44] and fluid flow within muscle microstructure [45, 46, 32], but largely ignore the multiscale, elastic, active and spatial aspects of the problem. Here we adopt a complementary perspective by developing a minimal continuum model that integrates these aspects, identifies the relevant coarse-grained variables and key dimensionless parameters, and highlights general biophysical principles about cellular constraints on muscular performance limits.

To do this, we turn to a framework to describe fluid-filled sponge-like materials, poroelasticity, originally developed to understand the mechanics of water-logged soils [47] and other geophysical problems [48], but has since been applied to describe cartilage [49], passive muscle [50], living cells [14, 15, 13], rapid nastic motions [16, 17], active gels [51, 52, 53, 54] etc. In Fig. 1B, we schematize a generalization of this framework to integrate the molecular actomyosin kinetics with the anisotropic elasticity, activity and flow in the cell to describe muscle fibers as an active self-squeezing sponge.

## 2 Biophysical model

At a minimal level, we model the muscle fiber as a cylinder (length *L*, radius *R*) of a biphasic mixture of an active porous solid (*ϕ*: solid fraction) immersed in fluid (1 *− ϕ*: fluid fraction). Assuming that 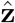 is along the long-axis of the fiber, the crystalline arrangement of interdigitated filaments and flexible proteins endows the sarcomere with a uniaxially anisotropic elastic stress (***σ***^*el*^) that is linearly related to the strain tensor ***∈*** = [***∇*u** + (***∇*u**)^*T*^ ]*/*2 (displacement 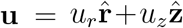, assuming axisymmetry), where the drained elastic moduli are computed as a function of *ϕ* using standard homogenization techniques (see SI Sec. IA for details). In addition, the interaction between the thick (myosin) and thin (actin) filaments also leads to an active stress (***σ***^*a*^). We assume that the passive elastic response of the porous solid is both anisotropic and compressible, approaching the incompressible limit only as *ϕ →* 1. The fluid stress, on the other hand, is dominated on large scales by an isotropic pressure *p*, whose gradients drive a flow velocity (**v**) with viscous dissipation being consequential only on the scale of the hydraulic pore size *𝓁*_*p*_ *∼* 20 *−* 55 nm [11, 8] (see SI Sec. IA for details). We emphasize that viscous forces are important on small scales, not because they balance individual motor forces (they don’t [1, 46]), but because they balance large-scale spatial *gradients* (*∼* 1*/L*) of the active stress. Mass and momentum conservation then collectively dictate overall force balance, global incompressibility and force balance in the fluid (Darcy’s law) as follows (see SI Sec. IA for details)

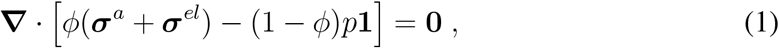

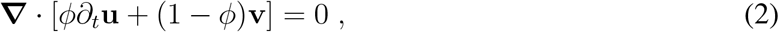

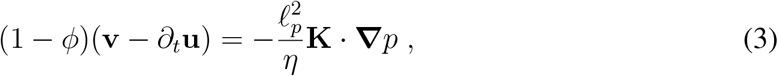

where *η* is the fluid viscosity and 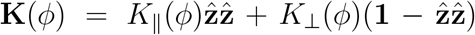 is a *ϕ*-dependent anisotropic permeability tensor (see SI Sec. IA for details). We note that the anisotropic structure of the sarcomere ensures that the lattice becomes radially impermeable to fluid flow (*K*_*⊥*_ *→* 0 as *ϕ → ϕ*_*∗*_ *≈* 0.91, see SI Sec. IA for details) before becoming incompressible (*ϕ →* 1). Hence volumetric deformations leading to intracellular flow are inevitable for all physiologically relevant *ϕ ∼* 0.1 *−* 0.22 [11, 8].

To determine the active stress ***σ***^*a*^ due to the molecular kinetics of the force-generating myosin motors, we use a simple two state model with *n*_*m*_(**x**, *t*) being the coarse-grained fraction of bound myosin motors and *⟨ y ⟩* (**x**, *t*) being the average extension of the motor head, at a given time *t* and position **x** in the cell. In the mean-field limit, the coupled dynamics of the actomyosin crossbridges is then given by (see SI Sec. IB for details)

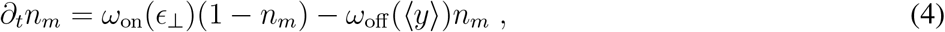

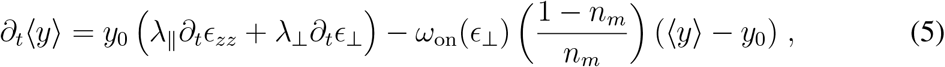

where *y*_0_ *∼* 8 *−* 10 nm [55] is the motor prestrain generated during the powerstroke and *λ* _*‖*_, *λ*_*⊥*_ are factors associated with the geometry of the binding crossbridge [56, 57] (see SI Sec. IB for details). The kinetic rates minimally incorporate biophysical feedback (Fig. 1B; see SI Sec. IB for details) through a load-dependent unbinding rate (*ω*_off_ (*⟨y ⟩*)) [58, 59, 60] which allows for stretch-activation [61], and a strain or lattice-spacing dependent binding rate (*ω*_on_(*∈*_*⊥*_)) due to mechanisms including lattice geometry [62, 36, 37], titin [63] etc., that allow for length-based regulation of force underlying the well-known Frank-Starling law [2]. As *ω*_on*/*off_ represent effective coarse-grained kinetic rates, we do not distinguish between different microscopic mechanisms of feedback. Assuming the myosin head behaves like a spring with stiffness *k*_*m*_, size *d*_*m*_ and a linear density *N* along the thick filament, the active contractile force density is *F*_*m*_ = *−k*_*m*_*n*_*m*_*⟨y ⟩* which gives rise to an anisotropic active stress 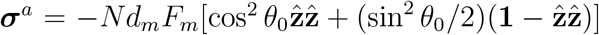 [64]. The active stress importantly includes both axial and radial components of the active force which are governed by the average crossbridge binding angle *θ*_0_ [56, 57, 36] (Fig. 1B; see SI Sec. IB for details). Eqs. 1-5 supplemented by appropriate boundary and initial conditions complete the specification of our multiscale continuum model.

## 3 Active hydraulic oscillations

To understand the dynamical consequences of the model, we consider two simple limits. In the passive, isotropic limit (***σ***^*a*^ = **0**, *K* _*‖*_= *K*_*⊥*_), we recover the classical result [47] that the hydrostatic pressure (*p*) equilibrates across a length *L* diffusively on a poroelastic timescale *τ*_*p*_ *∼* (*η/E*)(*L/𝓁*_*p*_)^2^ (see SI Sec. II for details), that combines the material (viscosity *η*, elastic modulus *E*) with the microstructural (pore size *𝓁*_*p*_). Anisotropy generalizes this result by distinguishing axial from radial flow. In contrast, in the active case, upon neglecting spatial heterogeneities and fluid flow (***∇****p ∼* **0**), our model matches previous kinetic theories of molecular motor assemblies [65, 66, 67, 68], where the kinetic timescale, *τ*_*k*_ = [*ω*_on_ + *ω*_off_ (*y*_0_)]^*−*1^ (see SI Sec. II for details) controls the residence time of bound motors and the rate of buildup of active stress. In this situation, if the load-dependent feedback is strong enough 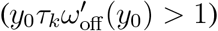, the molecular reaction develops an oscillatory instability with a characteristic frequency *ω ∼* 1*/τ*_*k*_ (see SI Sec. III for details). Combining the two limits and noting that in an anisotropic fiber, radial flow dominates axial flow, a key dimensionless parameter emerges - the radial *poroelastic Damköhler* number

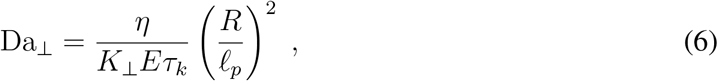

that captures the relative importance of radial fluid permeation (a mesoscopic time) to actomyosin kinetics (a molecular time). A similar measure for axial flow can also be constructed, see SI Sec. II. Assuming typical values for *E ∼* 0.1 *−* 10 MPa, *η ∼* 10^*−*3^ Pa.s, *𝓁*_*p*_ *∼* 20 *−* 60 nm, *τ*_*k*_ *∼* 1 *−* 10 ms and *R ∼* 5 *−* 100 *μ*m [11, 6, 69], we obtain a wide range of Da_*⊥*_ *∼* 10^*−*2^ *−* 10^3^ that is accessible to and seems to be exploited by the evolutionary range of muscle physiology (Fig. 3B).

**Figure 3.**
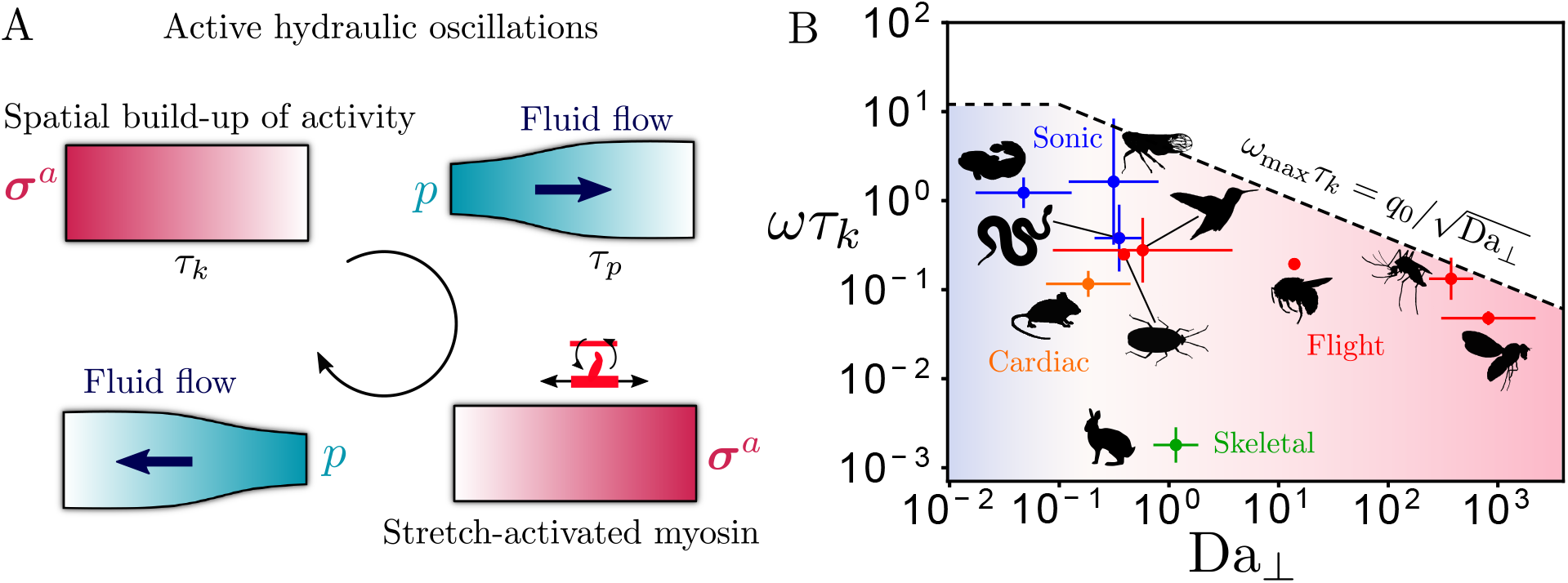
Active hydraulics limits muscular contraction rates. (A) Active hydraulic oscillations involving a periodic buildup of spatial gradients in active stresses along with a sloshing of fluid occur generically with a frequency 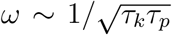 that combines the fast crossbridge cycling time (*τ*_*k*_) with the slow fluid permeation time (*τ*_*p*_). (B) Characteristic oscillation frequencies (*ω*) of contractions in fast sonic (blue), flight (red), cardiac (orange) and skeletal (green) muscles across species plotted against the estimated radial Damköhler number (Da_*⊥*_, Eq. 6). The data are compiled in SI Table 1 and presented as geometric means with the error bars representing the full range of estimated parameters. Active hydraulic oscillations dominate for Da_*⊥*_ *≥* 1 (shaded red region) with the theoretical scaling limit *ω*_max_ from Eq. 7 plotted (dashed line with constant *q*_0_ *∼* 3.83 computed for the longest wavelength radial mode, see SI Sec. VB and Fig. S2 for details). When Da_*⊥*_ *«* 1 (shaded blue region), flow is irrelevant and kinetics dominates, so the oscillation frequency saturates. Consistent with our prediction, the rates of spontaneous muscular contractions in the representative examples do not exceed the limit set by active hydraulics. Animal silhouettes are adapted from the online open database PhyloPic and individual image credits are in the SI Sec. VD.

When poroelastic, active and kinetic effects are all incorporated, we obtain a novel oscillatory instability that relies on spatial gradients in strain (Fig. 3A). A local buildup of active stress (on timescale *τ*_*k*_) squeezes the sarcomere, forcing fluid to flow and distend neighbouring regions of the lattice (on a timescale *τ*_*p*_), which in turn induces further buildup of myosin via stretch activation (in space). This mechanism of ‘active hydraulics’ intrinsically couples intracellular fluid flow, spatial deformation gradients and active stress generation through mechanical feedback. To see this, a minimal one-dimensional (1D) description with *∈*_*⊥*_ = 0 suffices (appropriate for *Drosophila* flight muscle [30]). Axial force balance along the muscle fiber implies 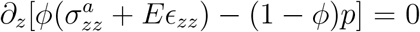, while poroelastic flow dictates 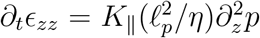 . Considering the slowest modes on scale of the system (*L*), these equations together (see SI Sec. IIIA for details) yield 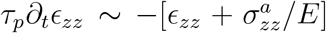 . In the limit when the active stress is slaved to the density of bound motors (see SI Sec. IIIA for details), we can write 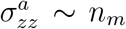 and linearize the kinetics about the steady state motor density 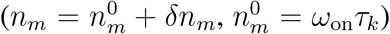 to obtain 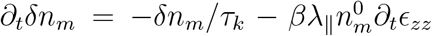 that includes the feedback mechanism through 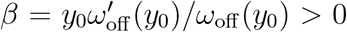 (see SI Sec. IIIA for details). For strong enough activity the coupled dynamics undergoes a Hopf bifurcation resulting in the spontaneous emergence of active hydraulic oscillations with a characteristic frequency 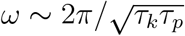 which predicts remarkably well the fruit fly wing-beat frequency *ω ≃* 150 *−* 160 Hz upon using estimates of *τ*_*k*_ *∼* 0.3 ms [4] and *τ*_*p*_ *∼* 5 *−* 6 s (see SI Sec. IIIA for details).

For general 3D deformations, fluid is more easily shunted radially rather than axially due to a smaller hydraulic resistance across a slender fiber (*R*^2^*/K*_*⊥*_ *« L*^2^*/K*_*‖*_) and the reduced permeability of Z-discs, a feature that survives the inclusion of heterogeneity in pore size. Hence, the radial, i.e., fastest poroelastic time (not the slowest) controls pressure relaxation and is hydraulically rate-limiting. Upon extending the previous 1D instability calculation to allow radial flow, spontaneous oscillations now emerge with a scaled characteristic frequency (for large Da_*⊥*_)

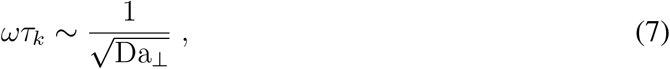

that involves both the kinetic and poroelastic time scales; a careful calculation shows that this phenomenon persists more generally (see SI Sec. III for details). Strikingly, spatiotemporal volumetric deformations trigger active hydraulic oscillations (inevitable for Da_*⊥*_ *≥* 1), even when the instability mechanism is kinetic (see SI Sec. III for details), hence offering a natural explanation for the experimental data in Fig. 2. Thus, we see that active hydraulics, rather than just kinetics, determines the fastest rate of spontaneous muscle contraction; in Fig. 3B, we plot the scaling relation in Eq. 7 and compare it to existing experimental data on muscular contractions across the animal kingdom. As our analysis focuses on rate limitations intrinsic to muscle fibers, we neglect constraints set by calcium and neuromuscular control, that can be surpassed in ultrafast contractions, for e.g., in asynchronous insect flight muscle [69]. Note that, Ca^+2^ cycling only introduces additional microscopic time scales that effectively replace actomyosin kinetic rates, if slower. Using representative estimates of the poroelastic, kinetic and contraction time scales in fast sonic (blue), flight (red), cardiac (orange) and skeletal (green) muscles (see SI Sec. VB and Table 1 for details), we find that while synchronous muscles are typically dominated by kinetics (Da_*⊥*_ *<* 1, shaded blue region), asynchronous muscles responsible for insect flight are often hydraulically dominated (Da_*⊥*_ *≥* 1, shaded red region), and the data is consistent with the maximal contraction rate being set by active hydraulics.

While features such as Ca^+2^ regulation and external inertial loads that are neglected here play a role in dictating muscle contraction rates, our analysis suggests that active hydraulics is also an important limiter of contraction rates in physiological settings. Direct measurements of spatial gradients of deformation and intracellular fluid flow in contracting muscle would provide a concrete test of these predictions.

## 4 Nonreciprocal mechanics of an odd elastic engine

We now show that spatial 3D deformations also have unusual mechanical and energetic implications. The mechanical response of muscle is quantified by the relation between stresses (forces) and strains (deformations) which can be time (or frequency) dependent in complex viscoelastic materials [70]. For small deformations, we can linearize the dynamical equations (Eqs. 1-5) about the resting state, Fourier transform in time (***∈*** (*ω*) =∫ d*t e*^*−iωt*^ ***∈*** (*t*)) and obtain an effective (visco)elastic constitutive relation *σ*_*ij*_(*ω*) = *𝒜*_*ijkl*_(*ω*)*E*_*kl*_(*ω*) that linearly relates the total stress (***σ*** = *ϕ*(***σ***^*el*^ + ***σ***^*a*^) *−* (1 *− ϕ*)*p***1**) to the strain tensor, shown pictorially for the shear and isotropic components in Fig. 4A (see SI Sec. IV for details). The frequency dependent complex modulus ***𝒜*** includes both an (in-phase) elastic response Re[***𝒜***] and an (out-of-phase) viscous response Im[***𝒜***]. In a passive system, time-reversal symmetry enforces *𝒜*_*ijkl*_(*ω*) = *𝒜*_*klij*_(*ω*) [71], but in an active system such as muscle, the lack of energy conservation allows new nonsymmetric terms (recently christened ‘odd (visco)elasticity’ [34, 72] in chiral and active media) that violate a fundamental property of mechanics - Maxwell-Betti reciprocity [71, 73]. Thus, in addition to the bulk (*B*_eff_ (*ω*)) and Young’s moduli (*Y*_eff_ (*ω*)) that govern the standard compressive and shear response and acquire frequency dependent corrections from motor kinetics, muscle’s uniaxial anisotropy couples isotropic dilations to shear via a passive anisotropic modulus *C*(*ω*) along with an exclusively active and nonreciprocal ‘odd-modulus’ *ζ*(*ω*) (Fig. 4A). While complete expressions for the frequency dependent moduli in terms of microscopic parameters are complicated and provided in SI Sec. IV, one simplifying limit assumes a uniform horizontal motor binding geometry (*θ*_0_ = 0) which yields the following expression for the odd-modulus

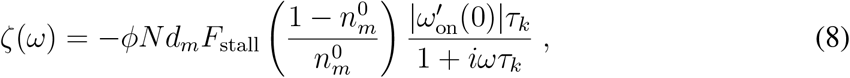

where the stall force 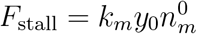 is the average active force generated by unloaded motors and 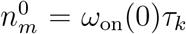 is the steady-state bound motor fraction (zero load duty-ratio). It is worth emphasizing that spatial anisotropy and activity are sufficient for odd (visco)elasticty to emerge in muscle, even without chiral effects that are often invoked [34]. Interestingly, the possible presence of such an odd-modulus in muscle was noted in passing in old work [74], though its implications were unrecognized.

**Figure 4.**
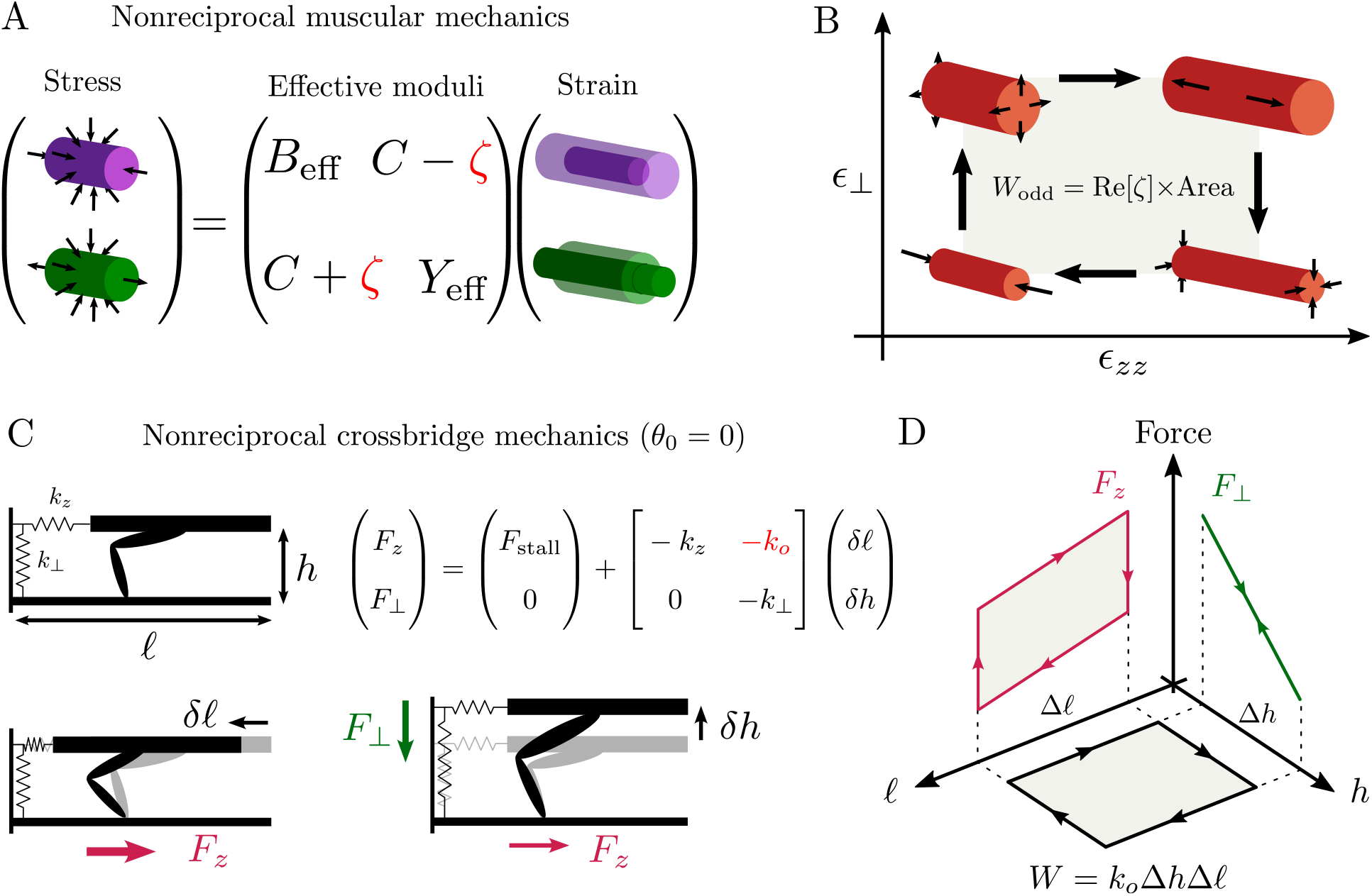
Odd elasticity and strain cycle engine in muscle. (A) A pictorial representation of a stress-strain relation involving shear and volume changes, for a 3D continuum material. As muscle fibers are active, the matrix of effective moduli can be nonsymmetric, with the asymmetry quantified by an odd or nonreciprocal modulus *ζ*. While the bulk (*B*_eff_ ) and Young’s (*Y*_eff_ ) moduli are present in isotropic solids, the passive coupling *C* is only present in anisotropic solids and *ζ* requires both anisotropy and activity to be present. (B) The presence of *ζ* ≠ 0 allows work to be actively produced by cycling strains in different directions, say axial and radial (right). The work done by the odd-modulus (*W*_odd_, Eq. 9) is proportional to odd-elasticity (Re[*ζ*]) and the signed area enclosed by the closed loop in strain space. (C) The average quasistatic elastic response of a crossbridge is nonreciprocal. For the simple binding geometry with *θ*_0_ = 0, axial forces (*F*_*z*_, magenta) include both active and passive (spring constant *k*_*z*_) contributions, while radial forces (*F*_*⊥*_, green) are purely passive (spring constant *k*_*⊥*_). As crossbridge binding kinetics depends on filament spacing (*h*), the average axial active force decreases with increasing radial stretch (*δh*), but the radial force is unchanged by an axial stretch (*δ𝓁*), hence the response is nonreciprocal. Linearized force response about rest state (with constant stall force *F*_stall_) shown for small radial and axial stretch, with the nonreciprocal odd spring constant *k*_*o*_ shown. (D) A strain cycle in axial and radial deformations generates work proportional to the area of the cycle and the odd spring constant *k*_*o*_, i.e., *W* = *k*_*o*_Δ*h*Δ*𝓁*. The corresponding axial (*F*_*z*_, magenta) and radial (*F*_*⊥*_, green) forces are plotted, showing conventional work loops.

A unique consequence of nonreciprocal mechanics is the ability to generate work from cycles of strain (Fig. 4B). Spontaneous strain cycles in muscle fibers (see Fig. 2B,D) have been previously interpreted as a time-varying Poisson ratio [32], but in an active medium, these cycles perform work. Upon writing 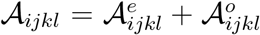 in terms of ‘even’ and ‘odd’ tensors (under the exchange of *ij* and *kl* indices), we can compute the cumulative mechanical work in a cycle as *W* = *−∮σ*_*ij*_d*E*_*ij*_ (*W >* 0 when work is produced, *W <* 0 when dissipated), which includes two qualitatively different terms *W* = *W*_even_ + *W*_odd_ (see SI Sec. IV for details). The even viscous term 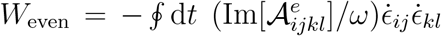 depends on the traversal frequency *ω* and *strain rates* (***⋵***) whereas the odd elastic term 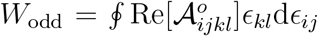 depends on *strain* (see SI Sec. IV for details). While *W*_even_ is simply an anisotropic generalization of standard viscous dissipation (Im[***𝒜***]*/ω* is like a viscosity), an intuitive explanation for *W*_odd_ is that, in the absence of energy conservation, cyclic deformations in different directions do not bring the system back to its initial energy, hence work can be either produced or absorbed. In the axisymmetric limit, relevant for muscle, we can simplify *W*_odd_ as (see SI Sec. IV for details)

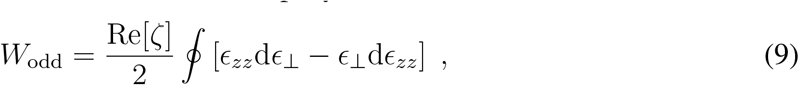

i.e., work from odd elasticity depends on the odd-modulus *ζ* and the area enclosed by a loop in the space of axial and transverse strains (Fig. 4B). Crucially this mechanism of power generation relies on 3D *spatial* deformations (axial and radial), rather than *temporal* variations.

What is the microscopic origin of muscular odd elasticity? We show that a crossbridge, upon averaging over stochastic binding events, behaves as a nonreciprocal mechanical element. To illustrate this point, we consider the simplifying limit of rate-independent (quasistatic) deformations and a horizontal binding geometry (*θ*_0_ = 0), so all active forces are purely axial and molecular kinetics are equilibriated (Fig. 4C). As the binding rate depends on the filament spacing (*h*), a radial stretch (*δh*) modifies the active axial force (*F*_*z*_) exerted, but an axial stretch (*δ𝓁*) generates no radial force (*F*_*⊥*_), which is purely passive when *θ*_0_ = 0. This asymmetry in response generalizes to arbitrary binding geometries and highlights the mechanical nonreciprocity of a crossbridge. Notably, this previously seemingly unrecognized general property is present in most microscopic crossbridge models that account for lattice deformations [75, 36, 37, 27].

To see this more explicitly, we linearize the averaged forces about stall conditions for small deformations, keeping *θ*_0_ = 0, to obtain *F*_*z*_ = *F*_stall_ *− k*_*z*_*δ𝓁 − k*_*o*_*δh* and *F*_*⊥*_ = *−k*_*⊥*_*δh*, where *k*_*z*_ (*k*_*⊥*_) are axial (radial) spring constants that account for passive elasticity (Fig. 4C). Non-reciprocity is then quantified by an “odd” spring constant 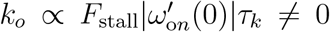 which originates from the strain dependent binding kinetics (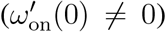 ) and is necessarily active (*F*_stall_ ≠ 0). Comparing with the macroscopic odd modulus in Eq. 8, we directly see *ζ ∝ −k*_*o*_.

Although *k*_*o*_ is a rate-independent, elastic constant (i.e., coefficient relating force to deformation), it does *not* conserve energy. As a result mechanical work done by the crossbridge is history dependent. Performing a quasistatic deformation cycle generates work given by *W* = *− ∮* (*F*_*z*_d*𝓁* + *F*_*⊥*_d*h*) which is equal to *k*_*o*_ times the area enclosed by the strain cycle (Fig. 4D). A more conventional work loop analysis (i.e., area enclosed by the *force*-displacement curve) [76] recapitulates the same result, as long as both axial and radial deformations and forces are correctly accounted for (Fig. 4D).

Where might we see these effects in muscle? While a fully 3D characterization of muscle’s viscoelastic response is currently unavailable, we use our model to analyze existing muscle rheology experiments [77, 78, 79] in a simpler 1D geometry that measures the uniaxial response (*σ*_*zz*_) in skinned muscle fibers subject to small amplitude oscillatory axial strains (*∈*_*zz*_). Hydraulic effects are assumed to be irrelevant in these experiments as the fibers are permeabilized. By compiling data across muscle types and species (*Drosophila* insect flight [77], mouse cardiac [78] and rabbit skeletal [79]), we fit our biophysical model to the measured effective (complex) Young’s modulus *Y*_eff_ (*ω*) = *σ*_*zz*_(*ω*)*/ ∈*_*zz*_(*ω*) (Fig. 5, see SI Sec. VC for analysis details).

**Figure 5.**
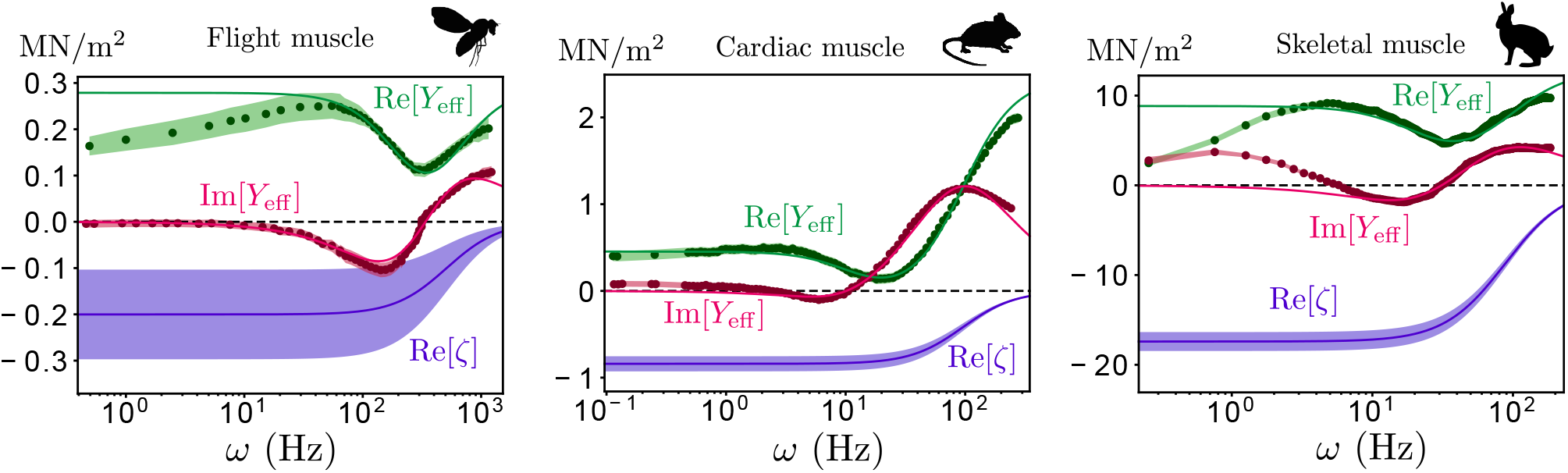
Active and nonreciprocal viscoelasticity across muscle types and species. Model fits of linear response (lines) to experimental data (circles are the mean, shaded region is one standard deviation) of sinusoidal uniaxial rheology digitally extracted and compiled from Ref. [77] (*Drosophila* flight muscle, left), Ref. [78] (mouse cardiac muscle, center) and Ref. [79] (rabbit skeletal muscle, right). As the muscle fibers are small and permeabilized, we assume affine deformations and neglect pressure gradients (see SI Sec. VC for fit details). The real and imaginary parts of the response correspond to the elastic (Re[*Y*_eff_ ], green) and the viscous (Im[*Y*_eff_ ], magenta) moduli. Dashed black line marks zero. The low frequency response in the experiments sometimes includes a weak power law or logarithmic dependence on frequency, possibly arising from internal passive processes that we neglect. Model estimates using Eq. 8 of the nonreciprocal (odd) elastic modulus Re[*ζ*] are shown in blue, with shaded region denoting one standard deviation (see SI Sec. VC for details). Animal silhouettes are adapted from the online open database PhyloPic and individual image credits are in SI Sec. VD.

In all three cases, a common characteristic response is seen - the elastic modulus (Re[*Y*_eff_ ], Fig. 5 green) transitions from a low frequency (largely passive) stiffness to a high-frequency rigor-like response from transiently crosslinked actomyosin, passing through an intermediary softening on time scales associated with crossbridge cycling, i.e., when the kinetic cycling of motors enhances compliance via filament sliding. The viscous modulus (Im[*Y*_eff_ ], Fig. 5 magenta) on the other hand is *negative* at low frequency (hence active and work producing), switching to positive (dissipative) values at higher frequencies (skeletal muscle displays additional features at low frequency arising from passive dissipation in sarcomeric polymers that we neglect, Fig. 5 right). In the 1D setting, the negative viscous modulus offers the only route to produce positive work through temporal changes in strain and odd-elastic effects are absent. Surprisingly, Im[*Y*_eff_ ] vanishes precisely at the frequency of pure kinetic oscillations (see SI Sec. VC for details), indicating that either hydraulics (which modifies the oscillation frequency) or 3D deformations (allowing odd elasticity) are essential for sustained power generation from cyclic contractions.

Using our model fit and known structural parameters, we estimate the frequency dependent odd-elastic modulus (Re[*ζ*]) for various muscle types (Fig. 5, blue lines; see SI Sec. VC for details). Notably, we see that the odd-modulus is predominantly negative and it remains nonvanishing at low frequencies, where its magnitude is controlled by the active stress and the strain based feedback on myosin kinetics (see Eq. 8). As Fig. 2B,D show, strain cycles (enclosing negative area) can be seen during spontaneous muscle contractions, which when combined with our model prediction of Re[*ζ*] *<* 0 and Eq. 9 are consistent with strain cycles actively producing work through odd elasticity. Using Re[*ζ*] *∼ −*(0.1 *−* 10) MPa (*ω <* 10 Hz,

Fig. 5) and typical strain cycles in Fig. 2, we estimate the odd elastic work that can be produced to be *W*_odd_ *∼* 0.2 *−* 20 kPa. Assuming an oscillation frequency *ω* = 10 Hz and muscle mass density *ρ*_*m*_ = 10^3^ kg/m^3^, odd elasticity can generate a mass specific power *P*_odd_ *∼ ωW*_odd_*/ρ*_*m*_ *∼* 2 *−* 200 W/kg which can be potentially significant in physiological conditions [76]. A direct test of our predictions of odd elasticity would be afforded by considering the 3D structural and force dynamics of muscle fibers using a combination of X-ray based methods [29, 23, 33] and force spectroscopy techniques [80, 81].

## 5 Discussion

Recognizing the importance of spatial strain gradients and the dynamics of fluid movement leads naturally to an emergent maximal rate of muscle contraction *ω*_max_ = *ω*_max_(*τ*_*k*_, *𝓁*_*p*_, *η, E, R, L, · · ·*) based on active hydraulic oscillations that combines a molecular kinetic time scale (*τ*_*k*_) with a mesoscopic poroelastic time scale (*τ*_*p*_), incorporating micro-structural (*𝓁*_*p*_) and macro-geometric (*R, L*) scales as well as material properties (*η, E*). Our analysis of published 3D spatial deformation data complements previous temporal studies of muscle rheology and highlights how muscle functions as an active elastic engine, whereby work can be produced from strain cycles via an emergent nonreciprocal response naturally present in anisotropic active solids. Additionally, we may estimate the maximal power density *P*_max_ based on a (size independent) maximal strain (*∈*_max_) and stress (*σ*_max_) of muscle [82], *P*_max_ *∼ σ*_max_*∈*_max_*ω*_max_. When molecular kinetics dominates (Da_*⊥*_ *<* 1, *ω*_max_ *∼* 1*/τ*_*k*_), muscular power is not constrained by size (*P*_max_ *∝ L*^0^). But in the active hydraulic regime (Da_*⊥*_ *>* 1,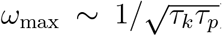 ), the maximal power density decreases with size (*P*_max_ *∝* 1*/L*), so larger organisms may need additional spring based mechanisms to amplify power output [83].

Beyond muscle, the principles of active hydraulics may also apply to other soft contractile systems. For example, most current soft muscle mimetic actuators [84, 85, 86, 87] have contraction rates that are largely limited by the slow propagation of the actuation signal which is typically diffusive, with *ω*_max_ *∼* 1*/L*^2^ [88]. Instead, our work suggests that if soft actuators were triggered locally rather than externally, then active hydraulics enables an alternate mechanism for faster contractions with *ω*_max_ *∼* 1*/L*.

Our results highlight the importance of a spatiotemporally integrated and multiscale view of muscle physiology for understanding this ubiquitous effector of soft power underlying animal movement. While we have demonstrated general biophysical consequences of an active sponge-like description of muscle, the specific physiological regimes and organisms where active hydraulics and odd elasticity are functional remains to be explored. A natural system to study this might be insect flight muscle, given its evolutionary age, phylogenetic breadth, and relatively simple physiology [89]. Further work incorporating tissue-scale response, neural control, Ca^+2^ signaling, inertial loading response etc., is required to understand the generality of the phenomena studied here, and connect with more macroscopic approaches to comparative biomechanics [90]. Ultimately, we will need an integrative view of muscle that spans the range from molecular motors to whole tissues to determine its ultimate performance limits and its failure modes, and thence eventually uncover its evolutionary trajectories and physiological functions.

## Acknowledgments

SS acknowledges support from the Harvard Society of Fellows, and LM acknowledges partial support from the NSF-Simons Center for Mathematical and Statistical Analysis of Biology 1764269, the Simons Foundation and the Henri Seydoux Fund. We thank Shriya Srinivasan for useful discussions.

## Author Contributions Statement

LM conceived the research topic/approach; SS and LM formulated the theoretical model; SS performed the analytical calculations; SS compiled and analyzed the data; SS and LM wrote the paper.

## Methods

Supplementary Information provides detailed descriptions of the theoretical model, analytical calculations, data acquisition and reanalysis.

## Data Availability

All original data supporting the findings of this work were obtained from published literature as indicated in the Supplementary Information (Sec. V). All reanalyzed versions of the data used in this work are available within the paper and Supplementary Information (Sec. V).

## References

[1] A. Huxley. Reflections on Muscle. Sherrington lectures. Liverpool University Press, 1980.

[2] Dorothy M Needham. Machina carnis: the biochemistry of muscular contraction in its historical development. Cambridge University Press, 1971.

[3] Miklós Nyitrai, Rosetta Rossi, Nancy Adamek, Maria Antonietta Pellegrino, Roberto Bottinelli, and Michael A Geeves. What limits the velocity of fast-skeletal muscle contraction in mammals? Journal of molecular biology, 355(3):432–442, 2006.

[4] Douglas M Swank, Vivek K Vishnudas, and David W Maughan. An exceptionally fast actomyosin reaction powers insect flight muscle. Proceedings of the National Academy of Sciences, 103(46):17543–17547, 2006.

[5] Andrew F Mead, Nerea Osinalde, Niels Ørtenblad, Joachim Nielsen, Jonathan Brewer, Michiel Vellema, Iris Adam, Constance Scharff, Yafeng Song, Ulrik Frandsen, et al. Fundamental constraints in synchronous muscle limit superfast motor control in vertebrates. Elife, 6:e29425, 2017.

[6] Albert M Gordon, Earl Homsher, and Mike Regnier. Regulation of contraction in striated muscle. Physiological reviews, 80(2):853–924, 2000.

[7] Ivana Y Kuo and Barbara E Ehrlich. Signaling in muscle contraction. Cold Spring Harbor perspectives in biology, 7(2):a006023, 2015.

[8] Joseph D Powers, Sage A Malingen, Michael Regnier, and Thomas L Daniel. The sliding filament theory since andrew huxley: Multiscale and multidisciplinary muscle research. Annual Review of Biophysics, 50:373–400, 2021.

[9] Lawrence C Rome and Stan L Lindstedt. The quest for speed: muscles built for high-frequency contractions. Physiology, 13(6):261–268, 1998.

[10] Douglas A Syme and Robert K Josephson. How to build fast muscles: synchronous and asynchronous designs. Integrative and comparative biology, 42(4):762–770, 2002.

[11] Barry M Millman. The filament lattice of striated muscle. Physiological reviews, 78(2):359–391, 1998.

[12] Joseph D Powers, C David Williams, Michael Regnier, and Thomas L Daniel. A spatially explicit model shows how titin stiffness modulates muscle mechanics and energetics. Integrative and comparative biology, 58(2):186–193, 2018.

[13] Alex Mogilner and Angelika Manhart. Intracellular fluid mechanics: Coupling cytoplasmic flow with active cytoskeletal gel. Annual Review of Fluid Mechanics, 50:347–370, 2018.

[14] Guillaume T Charras, Timothy J Mitchison, and L Mahadevan. Animal cell hydraulics. Journal of cell science, 122(18):3233–3241, 2009.

[15] Emad Moeendarbary, Leó Valon, Marco Fritzsche, Andrew R Harris, Dale A Moulding, Adrian J Thrasher, Eleanor Stride, L Mahadevan, and Guillaume T Charras. The cytoplasm of living cells behaves as a poroelastic material. Nature materials, 12(3):253–261, 2013.

[16] Yoël Forterre, Jan M Skotheim, Jacques Dumais, and Lakshminarayanan Mahadevan. How the venus flytrap snaps. Nature, 433(7024):421–425, 2005.

[17] Jan M Skotheim and Lakshminarayanan Mahadevan. Physical limits and design principles for plant and fungal movements. Science, 308(5726):1308–1310, 2005.

[18] Beáta Bugyi and Miklós Kellermayer. The discovery of actin:”to see what everyone else has seen, and to think what nobody has thought”. Journal of Muscle Research and Cell Motility, 41(1):3–9, 2020.

[19] Soham Ghosh, Benjamin Seelbinder, Jonathan T Henderson, Ryan D Watts, Adrienne K Scott, Alexander I Veress, and Corey P Neu. Deformation microscopy for dynamic intracellular and intranuclear mapping of mechanics with high spatiotemporal resolution. Cell reports, 27(5):1607–1620, 2019.

[20] Benjamin Kaminer. Water loss during contracture of muscle. The Journal of general physiology, 46(1):131–142, 1962.

[21] Károly Trombitás, Peter Baatsen, John Schreuder, and Gerald H Pollack. Contraction-induced movements of water in single fibres of frog skeletal muscle. Journal of Muscle Research & Cell Motility, 14(6):573–584, 1993.

[22] Albert Szent-Györgyi. The contraction of myosin threads. Studies from the Institute of Medical Chemistry University Szeged: Myosin and muscular contraction, 1:17–26, 1942.

[23] G Cecchi, MA Bagni, PJ Griffiths, CC Ashley, and Y Maeda. Detection of radial crossbridge force by lattice spacing changes in intact single muscle fibers. Science, 250(4986):1409–1411, 1990.

[24] JG Pinto and ROLAND Win. Non-uniform strain distribution in papillary muscles. American Journal of Physiology-Heart and Circulatory Physiology, 233(3):H410–H416, 1977.

[25] IR Neering, LA Quesenberry, VA Morris, and SR Taylor. Nonuniform volume changes during muscle contraction. Biophysical journal, 59(4):926–933, 1991.

[26] Soham Ghosh, James G Cimino, Adrienne K Scott, Frederick W Damen, Evan H Phillips, Alexander I Veress, Corey P Neu, and Craig J Goergen. In vivo multiscale and spatially-dependent biomechanics reveals differential strain transfer hierarchy in skeletal muscle. ACS biomaterials science & engineering, 3(11):2798–2805, 2017.

[27] Takumi Washio, Seine A Shintani, Hideo Higuchi, Seiryo Sugiura, and Toshiaki Hisada. Effect of myofibril passive elastic properties on the mechanical communication between motor proteins on adjacent sarcomeres. Scientific reports, 9(1):1–17, 2019.

[28] Fumiaki Kono, Seitaro Kawai, Yuta Shimamoto, and Shin’ichi Ishiwata. Nanoscopic changes in the lattice structure of striated muscle sarcomeres involved in the mechanism of spontaneous oscillatory contraction (spoc). Scientific reports, 10(1):1–14, 2020.

[29] Wai Pang Chan and Michael H Dickinson. In vivo length oscillations of indirect flight muscles in the fruit fly drosophila virilis. The Journal of experimental biology, 199(12):2767–2774, 1996.

[30] TC Irving and DW Maughan. In vivo x-ray diffraction of indirect flight muscle from drosophila melanogaster. Biophysical journal, 78(5):2511–2515, 2000.

[31] Michael Dickinson, Gerrie Farman, Mark Frye, Tanya Bekyarova, David Gore, David Maughan, and Thomas Irving. Molecular dynamics of cyclically contracting insect flight muscle in vivo. Nature, 433(7023):330–334, 2005.

[32] Julie A Cass, C Dave Williams, Tom C Irving, Eric Lauga, Sage Malingen, Thomas L Daniel, and Simon N Sponberg. A mechanism for sarcomere breathing: volume change and advective flow within the myofilament lattice. Biophysical Journal, 2021.

[33] Sage A Malingen, Anthony M Asencio, Julie A Cass, Weikang Ma, Thomas C Irving, and Thomas L Daniel. In vivo x-ray diffraction and simultaneous emg reveal the time course of myofilament lattice dilation and filament stretch. Journal of Experimental Biology, 223(17):jeb224188, 2020.

[34] Colin Scheibner, Anton Souslov, Debarghya Banerjee, Piotr Surówka, William TM Irvine, and Vincenzo Vitelli. Odd elasticity. Nature Physics, 16(4):475–480, 2020.

[35] Michel Fruchart, Colin Scheibner, and Vincenzo Vitelli. Odd viscosity and odd elasticity. Annual Review of Condensed Matter Physics, 14:471–510, 2023.

[36] C David Williams, Michael Regnier, and Thomas L Daniel. Axial and radial forces of cross-bridges depend on lattice spacing. PLoS computational biology, 6(12):e1001018, 2010.

[37] C David Williams, Mary K Salcedo, Thomas C Irving, Michael Regnier, and Thomas L Daniel. The length–tension curve in muscle depends on lattice spacing. Proceedings of the Royal Society B: Biological Sciences, 280(1766):20130697, 2013.

[38] P Bryant Chase, J Michael Macpherson, and Thomas L Daniel. A spatially explicit nanomechanical model of the half-sarcomere: myofilament compliance affects ca 2+-activation. Annals of biomedical engineering, 32:1559–1568, 2004.

[39] Srboljub M Mijailovich, Oliver Kayser-Herold, Boban Stojanovic, Djordje Nedic, Thomas C Irving, and Michael A Geeves. Three-dimensional stochastic model of actin– myosin binding in the sarcomere lattice. Journal of General Physiology, 148(6):459–488, 2016.

[40] Sarah Kosta, Dylan Colli, Qiang Ye, and Kenneth S Campbell. Fibersim: a flexible open-source model of myofilament-level contraction. Biophysical Journal, 121(2):175–182, 2022.

[41] Benjamin B Wheatley, Gregory M Odegard, Kenton R Kaufman, and Tammy L Haut Donahue. A case for poroelasticity in skeletal muscle finite element analysis: experiment and modeling. Computer methods in biomechanics and biomedical engineering, 20(6):598–601, 2017.

[42] LA Spyrou, S Brisard, and K Danas. Multiscale modeling of skeletal muscle tissues based on analytical and numerical homogenization. Journal of the mechanical behavior of biomedical materials, 92:97–117, 2019.

[43] Thomas J Roberts, Carolyn M Eng, David A Sleboda, Natalie C Holt, Elizabeth L Brainerd, Kristin K Stover, Richard L Marsh, and Emanuel Azizi. The multi-scale, three-dimensional nature of skeletal muscle contraction. Physiology, 34(6):402–408, 2019.

[44] David A Sleboda and Thomas J Roberts. Internal fluid pressure influences muscle contractile force. Proceedings of the National Academy of Sciences, 117(3):1772–1778, 2020.

[45] Loribeth Q Evertz, Sarah M Greising, Duane A Morrow, Gary C Sieck, and Kenton R Kaufman. Analysis of fluid movement in skeletal muscle using fluorescent microspheres. Muscle & nerve, 54(3):444–450, 2016.

[46] Sage A Malingen, Kaitlyn Hood, Eric Lauga, Anette Hosoi, and Thomas L Daniel. Fluid flow in the sarcomere. Archives of Biochemistry and Biophysics, 706:108923, 2021.

[47] Maurice A Biot. General theory of three-dimensional consolidation. Journal of applied physics, 12(2):155–164, 1941.

[48] Herbert F Wang. Theory of linear poroelasticity with applications to geomechanics and hydrogeology. Princeton University Press, 2017.

[49] Van C Mow, Mark H Holmes, and W Michael Lai. Fluid transport and mechanical properties of articular cartilage: a review. Journal of biomechanics, 17(5):377–394, 1984.

[50] Ming Yang and Larry A Taber. The possible role of poroelasticity in the apparent viscoelastic behavior of passive cardiac muscle. Journal of biomechanics, 24(7):587–597, 1991.

[51] Andrew C Callan-Jones and Frank Jülicher. Hydrodynamics of active permeating gels. New Journal of Physics, 13(9):093027, 2011.

[52] Markus Radszuweit, Sergio Alonso, Harald Engel, and Markus Bär. Intracellular mechanochemical waves in an active poroelastic model. Physical review letters, 110(13):138102, 2013.

[53] Yaron Ideses, Vitaly Erukhimovitch, Ron Brand, D Jourdain, J Salmeron Hernandez, UR Gabinet, SA Safran, Karsten Kruse, and A Bernheim-Groswasser. Spontaneous buckling of contractile poroelastic actomyosin sheets. Nature communications, 9(1):1–13, 2018.

[54] Christoph A Weber, Chris H Rycroft, and L Mahadevan. Differential activity-driven instabilities in biphasic active matter. Physical review letters, 120(24):248003, 2018.

[55] Andrew F Huxley and Ro M Simmons. Proposed mechanism of force generation in striated muscle. Nature, 233(5321):533–538, 1971.

[56] M Schoenberg. Geometrical factors influencing muscle force development. i. the effect of filament spacing upon axial forces. Biophysical journal, 30(1):51–67, 1980.

[57] M Schoenberg. Geometrical factors influencing muscle force development. ii. radial forces. Biophysical journal, 30(1):69–77, 1980.

[58] Claudia Veigel, Justin E Molloy, Stephan Schmitz, and John Kendrick-Jones. Load-dependent kinetics of force production by smooth muscle myosin measured with optical tweezers. Nature cell biology, 5(11):980–986, 2003.

[59] Bin Guo and William H Guilford. Mechanics of actomyosin bonds in different nucleotide states are tuned to muscle contraction. Proceedings of the National Academy of Sciences, 103(26):9844–9849, 2006.

[60] Gabriella Piazzesi, Massimo Reconditi, Marco Linari, Leonardo Lucii, Pasquale Bianco, Elisabetta Brunello, Valérie Decostre, Alex Stewart, David B Gore, Thomas C Irving, et al. Skeletal muscle performance determined by modulation of number of myosin motors rather than motor force or stroke size. Cell, 131(4):784–795, 2007.

[61] John William Sutton Pringle. The croonian lecture, 1977-stretch activation of muscle: function and mechanism. Proceedings of the Royal Society of London. Series B. Biological Sciences, 201(1143):107–130, 1978.

[62] Thomas C Irving, John Konhilas, Darold Perry, Robert Fischetti, and Pieter P De Tombe. Myofilament lattice spacing as a function of sarcomere length in isolated rat myocardium. American Journal of Physiology-Heart and Circulatory Physiology, 279(5):H2568–H2573, 2000.

[63] Younss Ait-Mou, Karen Hsu, Gerrie P Farman, Mohit Kumar, Marion L Greaser, Thomas C Irving, and Pieter P de Tombe. Titin strain contributes to the frank–starling law of the heart by structural rearrangements of both thin-and thick-filament proteins. Proceedings of the National Academy of Sciences, 113(8):2306–2311, 2016.

[64] M Cristina Marchetti, Jean-François Joanny, Sriram Ramaswamy, Tanniemola B Liverpool, Jacques Prost, Madan Rao, and R Aditi Simha. Hydrodynamics of soft active matter. Reviews of Modern Physics, 85(3):1143, 2013.

[65] Frank Jülicher and Jacques Prost. Spontaneous oscillations of collective molecular motors. Physical review letters, 78(23):4510, 1997.

[66] Stefan Günther and Karsten Kruse. Spontaneous waves in muscle fibres. new Journal of physics, 9(11):417, 2007.

[67] Thomas Guérin, Jacques Prost, Pascal Martin, and Jean-François Joanny. Coordination and collective properties of molecular motors: theory. Current opinion in cell biology, 22(1):14–20, 2010.

[68] Thomas Guérin, J Prost, and J-F Joanny. Dynamical behavior of molecular motor assemblies in the rigid and crossbridge models. The European Physical Journal E, 34(6):1–21, 2011.

[69] Robert K Josephson, Jean G Malamud, and Darrell R Stokes. Asynchronous muscle: a primer. Journal of Experimental Biology, 203(18):2713–2722, 2000.

[70] Roderic S Lakes. Viscoelastic solids, volume 9. CRC press, 1998.

[71] Maurice A Biot. Xliii. non-linear theory of elasticity and the linearized case for a body under initial stress. The London, Edinburgh, and Dublin Philosophical Magazine and Journal of Science, 27(183):468–489, 1939.

[72] Debarghya Banerjee, Vincenzo Vitelli, Frank Jülicher, and Piotr Surówka. Active viscoelasticity of odd materials. Physical Review Letters, 126(13):138001, 2021.

[73] C Truesdell. The meaning of betti’s reciprocal theorem. J. Research of NBS B, 67:85–86, 1963.

[74] George I Zahalak. Non-axial muscle stress and stiffness. Journal of theoretical biology, 182(1):59–84, 1996.

[75] Bertrand C W Tanner, Thomas L Daniel, and Michael Regnier. Sarcomere lattice geometry influences cooperative myosin binding in muscle. PLoS computational biology, 3(7):e115, 2007.

[76] Robert K Josephson. Mechanical power output from striated muscle during cyclic contraction. Journal of Experimental Biology, 114(1):493–512, 1985.

[77] Bertrand CW Tanner, Gerrie P Farman, Thomas C Irving, David W Maughan, Bradley M Palmer, and Mark S Miller. Thick-to-thin filament surface distance modulates cross-bridge kinetics in drosophila flight muscle. Biophysical journal, 103(6):1275–1284, 2012.

[78] Bradley M Palmer, Takeki Suzuki, Yuan Wang, William D Barnes, Mark S Miller, and David W Maughan. Two-state model of acto-myosin attachment-detachment predicts c-process of sinusoidal analysis. Biophysical journal, 93(3):760–769, 2007.

[79] Masataka Kawai and Philip W Brandt. Sinusoidal analysis: a high resolution method for correlating biochemical reactions with physiological processes in activated skeletal muscles of rabbit, frog and crayfish. Journal of Muscle Research & Cell Motility, 1(3):279–303, 1980.

[80] Otger Campas. A toolbox to explore the mechanics of living embryonic tissues. In Seminars in cell & developmental biology, volume 55, pages 119–130. Elsevier, 2016.

[81] Pere Roca-Cusachs, Vito Conte, and Xavier Trepat. Quantifying forces in cell biology. Nature cell biology, 19(7):742–751, 2017.

[82] James H Marden and Lee R Allen. Molecules, muscles, and machines: universal performance characteristics of motors. Proceedings of the National Academy of Sciences, 99(7):4161–4166, 2002.

[83] Mark Ilton, M Saad Bhamla, Xiaotian Ma, Suzanne M Cox, Leah L Fitchett, Yongjin Kim, Je-sung Koh, Deepak Krishnamurthy, Chi-Yun Kuo, Fatma Zeynep Temel, et al. The principles of cascading power limits in small, fast biological and engineered systems. Science, 360(6387):eaao1082, 2018.

[84] Seyed M Mirvakili and Ian W Hunter. Artificial muscles: Mechanisms, applications, and challenges. Advanced Materials, 30(6):1704407, 2018.

[85] Victor V Yashin and Anna C Balazs. Pattern formation and shape changes in self-oscillating polymer gels. Science, 314(5800):798–801, 2006.

[86] Pierre-Gilles De Gennes, Matthieu Hébert, and Rama Kant. Artificial muscles based on nematic gels. In Macromolecular Symposia, volume 113, pages 39–49. Wiley Online Library, 1997.

[87] PG De Gennes, Ko Okumura, M Shahinpoor, and Kwang J Kim. Mechanoelectric effects in ionic gels. Europhysics Letters, 50(4):513, 2000.

[88] Pierre-Gilles de Gennes. A semi-fast artificial muscle. Comptes Rendus de l’Academie des Sciences Series IIB Mechanics Physics Chemistry Astronomy, 5(324):343–348, 1997.

[89] Jeff Gau, James Lynch, Brett Aiello, Ethan Wold, Nick Gravish, and Simon Sponberg. Bridging two insect flight modes in evolution, physiology and robophysics. Nature, pages 1–8, 2023.

[90] David Labonte. A theory of physiological similarity in muscle-driven motion. Proceedings of the National Academy of Sciences, 120(24):e2221217120, 2023.

