## Supplementary Information for "Active hydraulics and odd elasticity of muscle fibers"

#### CONTENTS


I. Active poroelastic model of muscle 1
A. Active and anisotropic poroelasticity 1
B. Molecular motor kinetics 3

II. Nondimensionalization and linearized dynamics 6

III. Oscillatory instability 8
A. 1D model with zero transverse strain ($|\epsilon_\perp| \ll |\epsilon_{zz}|$) 8
B. 1D model with nonzero transverse strain ($|\epsilon_\perp| \sim |\epsilon_{zz}|$) 9
C. 3D oscillations with stable kinetics ($|\epsilon_\perp| \gg |\epsilon_{zz}|$) 13

IV. Linear response and nonreciprocal elasticity 15

V. Data compilation and analysis 17
A. 3D strains in spontaneous sarcomeric oscillations 17
B. Estimating kinetic, poroelastic and contraction time scales in muscle 19
C. Oscillatory muscle rheology experiments 20
D. Silhouette image credits 22

References 23


### I. ACTIVE POROELASTIC MODEL OF MUSCLE

We build a multiscale model of muscle that incorporates active contractility induced by the binding of myosin motors, the elasticity of the sarcomeres and permeation of fluid through a dense but porous elastomer. Various continuum models have been proposed previously to describe different aspects of motor-filament assemblies, both phenomenologically [S1–S7] and via systematic coarse-graining techniques [S8–S13]. While many of these models focus on dilute suspensions, the opposite limit of strongly crosslinked active elastomers (most relevant to muscle) has only been explored recently [S7, S11, S13]. We employ an approach similar to these recent studies here, by accounting for the relevant symmetries and structural features unique to muscle.

#### A. Active and anisotropic poroelasticity

The muscle cell is modeled on the mesoscale as a cylindrical fiber of length $L$ and radius $R$ ($R \ll L$) composed of a biphasic mixture of an active porous solid (volume fraction $\phi$) immersed in a passive Newtonian fluid (volume fraction $1-\phi$). Spatial coordinates are denoted by $\mathbf{x} = (r, \varphi, z)$ The deformation of the solid is assumed to be axisymmetric ($\partial_\varphi = 0$) with a displacement $\mathbf{u} = u_r\hat{\mathbf{r}} + u_z\hat{\mathbf{z}}$. The nonvanishing components of the linearized strain tensor $\boldsymbol{\epsilon} = [\boldsymbol{\nabla}\mathbf{u} + (\boldsymbol{\nabla}\mathbf{u})^T]/2$ are

$$\epsilon_{zz} = \partial_z u_z \ , \quad \epsilon_{rr} = \partial_r u_r \ , \quad \epsilon_{\varphi\varphi} = \frac{u_r}{r} \ , \quad \epsilon_{rz} = \frac{1}{2}(\partial_z u_r + \partial_r u_z) \ . \tag{S1}$$

The transverse strain is denoted by $\epsilon_\perp = \epsilon_{rr} + \epsilon_{\varphi\varphi} = \partial_r u_r + u_r/r$.

The solid component corresponding to the anisotropic myofibrillar lattice is characterized by both an elastic response and an active response. Notably, unlike nematic elastomers with their characteristic soft mode elasticity and vanishing shear response [S14, S15], the uniaxial nematic order of strongly aligned actomyosin filaments in the myofibril is quenched. As a result, the network displays a finite shear modulus even when passive without requiring the effects discussed in Ref. [S7]. Furthermore, we note that while the bare microscopic filaments are essentially incompressible, the finite porosity of the solid structure makes the material effectively compressible. Assuming a linear constitutive relation for the elastic stress $\boldsymbol{\sigma}^{el}$ (valid for small deformations), a uniaxially anisotropic (along $z$) solid has five independent elastic moduli in general [S16], corresponding to the transverse plane-strain bulk modulus ($B_\perp$), the longitudinal and transverse shear moduli ($G_z$ and $G_\perp$), the longitudinal Young's modulus ($Y_z$) and the major Poisson's ratio ($\nu_z$). Then we can write the general form of the elastic constitutive equation as

$$\sigma^{el}_\perp = 2B_\perp \left(\epsilon_\perp + 2\nu_z\epsilon_{zz}\right) \ , \quad \sigma^{el}_{zz} = 2\nu_z B_\perp \left(\epsilon_\perp + 2\nu_z\epsilon_{zz}\right) + Y_z\epsilon_{zz} \ , \tag{S2a}$$

$$\sigma^{el}_{rr} - \sigma^{el}_{\varphi\varphi} = 2G_\perp(\epsilon_{rr} - \epsilon_{\varphi\varphi}) \ , \quad \sigma^{el}_{r\varphi} = 2G_\perp\epsilon_{r\varphi} \ , \quad \sigma^{el}_{rz} = 2G_z\epsilon_{rz} \ . \tag{S2b}$$

where $\sigma^{el}_{\perp} = \sigma^{el}_{rr} + \sigma^{el}_{\varphi\varphi}$ is the transverse stress. For a given solid fraction $\phi$, the effective moduli for the drained solid can be obtained by coarse-graining over the microstructure using standard effective medium and homogenization techniques developed in the context of fibrous composites, see for instance Refs. [S17–S19]. Assuming the bare material is both isotropic and incompressible, i.e., with a single shear modulus $G_0$ and a microscopic Poisson's ratio $\nu_0 = 1/2$, the homogenized elastic moduli for the drained system are given by [S18, S19]

$$Y_z = 3G_0\phi \ , \quad G_z = G_\perp = G_0\left(\frac{\phi}{2-\phi}\right) \ , \quad B_\perp = G_0\left(\frac{\phi}{1-\phi}\right) \ , \quad \nu_z = \frac{1}{2} \ , \tag{S3}$$

where we have used (microscopic) incompressiblity to write the bare Young's modulus as $Y_{\text{bare}} = 2G_0(1+\nu_0) = 3G_0$. Although a uniaxial solid has five independent elastic coefficients, we only in fact have one independent elastic modulus here, $G_0$, and all the rest are related through the porosity $1-\phi$. Also, for $\phi \to 1$, the voids vanish and the transverse bulk modulus $B_\perp \to \infty$, recovering the pure incompressible bulk solid, consistent with our expectations. While the details of the drained moduli in Eq. S3 are specific to the coarse-graining technique used, there are simple scaling features in the limits of $\phi \to 0$ and $\phi \to 1$ in general.

The active stress $\boldsymbol{\sigma}^a$ includes the forces generated internally by the action of molecular motors and is constructed explicitly in Sec. I B below.

Finally, the permeating fluid has velocity $\mathbf{v}$ which is driven by gradients in the local hydrostatic pressure $p$. As inertia is negligible, global force balance of the solid and fluid composite material dictates

$$\boldsymbol{\nabla}\cdot\left[\phi(\boldsymbol{\sigma}^a + \boldsymbol{\sigma}^{el}) - (1-\phi)p\mathbf{1}\right] = \mathbf{0} \ . \tag{S4}$$

This is appropriate for the case when the solid constituent forming the porous skeleton is itself incompressible, as is relevant for us. In the absence of mass exchange or interconversion, pressure serves as a Lagrange multiplier that enforces total incompressibility which reads as

$$\boldsymbol{\nabla}\cdot[\phi\partial_t\mathbf{u} + (1-\phi)\mathbf{v}] = 0 \ , \tag{S5}$$

where the fluid velocity $\mathbf{v}$ is determined by Darcy's law [S20]

$$(1-\phi)(\mathbf{v} - \partial_t\mathbf{u}) = -\frac{\ell_p^2}{\eta}\mathbf{K}\cdot\boldsymbol{\nabla}p \ , \tag{S6}$$

with $\eta$ as the fluid viscosity, $\ell_p$ the pore size, and the uniaxially anisotropic permeability tensor $\mathbf{K} = K_\parallel\hat{\mathbf{z}}\hat{\mathbf{z}} + K_\perp(\mathbf{1} - \hat{\mathbf{z}}\hat{\mathbf{z}})$. Just like the elastic stress, the hydraulic permeability tensor can be computed by assuming viscous flow through a periodic hexagonal lattice of cylinders [S21–S25] as

$$K_\parallel = C_1\left(\frac{r_0}{\ell_p}\right)^2\frac{(1-\phi)^3}{\phi^2} \ , \quad K_\perp = C_2\left(\frac{r_0}{\ell_p}\right)^2\left[\left(\frac{\phi_*}{\phi}\right)^{1/2} - 1\right]^{5/2} \ , \tag{S7}$$

where $r_0 \sim 8-10$ nm is the typical radius of thick filaments *in-vivo* [S26] and the numerical constants $C_1 \sim 0.15$, $C_2 \sim 0.2$ vary weakly with $\phi$ [S22, S25]. We note that while the axial permeability $K_\parallel$ is the usual Carman-Kozeny formula, the transverse permeability $K_\perp$ is different and vanishes when the solid fraction approaches hexagonal close packing at $\phi_* = \pi/(2\sqrt{3}) \approx 0.907$, beyond which the fluid cannot percolate radially. Nonetheless, the matrix remains permeable along the long axis of the fiber ($K_\parallel \neq 0$). Upon taking a divergence of Eq. S6 and using incompressibility from Eq. S5, we obtain

$$\frac{\ell_p^2}{\eta}\boldsymbol{\nabla}\cdot\mathbf{K}\cdot\boldsymbol{\nabla}p = \partial_t(\boldsymbol{\nabla}\cdot\mathbf{u}) \ . \tag{S8}$$

In the absence of an active stress ($\boldsymbol{\sigma}^a = \mathbf{0}$), Eqs. S4, S5, S6 recover classical passive poroelasticity [S20] (in an anisotropic medium), with the three equations determining the three unknown variables - the solid displacement ($\mathbf{u}$), the fluid velocity ($\mathbf{v}$) and the fluid pressure ($p$). The elastic stress $\boldsymbol{\sigma}^{el} = \boldsymbol{\sigma}^{el}(\mathbf{u})$ is a function of the displacement (or strain) through the constitutive relation (Eq. S2), so it is not an independent variable. Later in Sec. I B we will see how adding the active stress ($\boldsymbol{\sigma}^a$) and the motor density ($n_m$) includes two more dynamical variables into the description of an *active* poroelastic material.

To supplement the bulk equations, we need some boundary conditions. The surface of the muscle fiber is a free boundary, so the normal component of the total stress $\boldsymbol{\sigma} = \phi(\boldsymbol{\sigma}^a + \boldsymbol{\sigma}^{el}) - (1-\phi)p\mathbf{1}$ vanishes at the surface, which is given by

$$\sigma_{rr}(R) = 0 \ , \quad \sigma_{zr}(R) = 0 \ . \tag{S9}$$

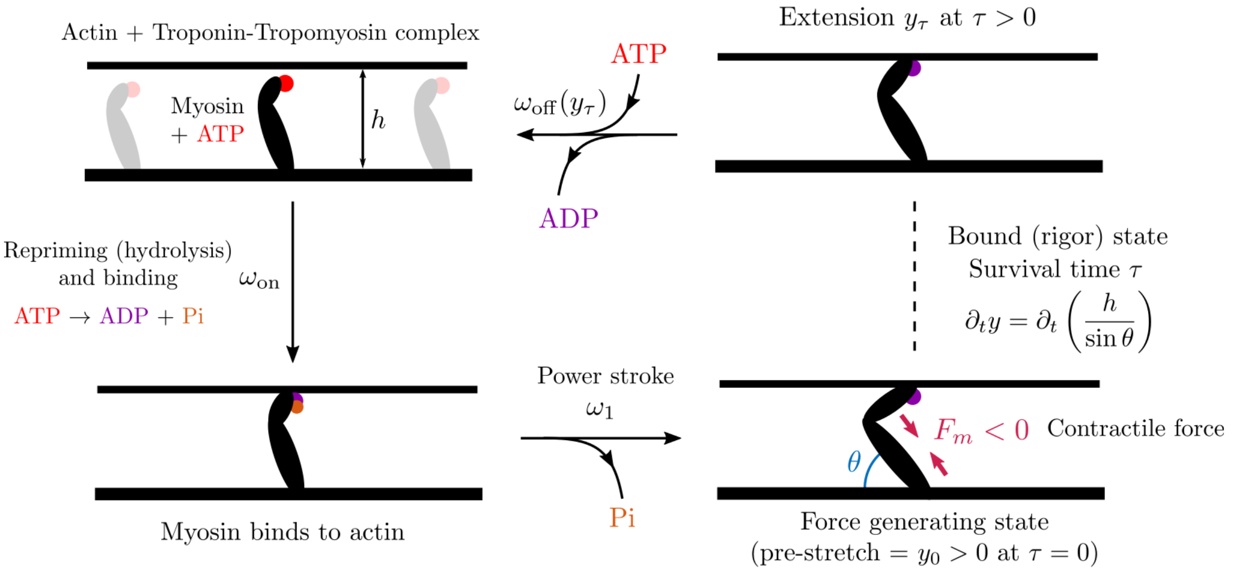


FIG. S1. A simplified sketch of the Lymn-Taylor cycle showing the different states of actin and myosin. Open binding sites on actin thin filments are always assumed to be available freely. Myosin motor heads with bound hydrolyzed ATP (binding and repriming steps are combined as one here for simplicity) attach to actin at an open binding site with a rate $\omega_{\rm on}$ and undergo a rapid powerstroke with rate $\omega_1 \gg \omega_{\rm on}$. The phosphate release step triggers a conformation change in the motor head that is modeled simply as a prestrain $y_0 > 0$, which corresponds to an initial contractile force density $F_m = -k_m n_m y_0 < 0$. The motor remains bound for a survival time $\tau$ during which the motor strain tracks the local deformation of the network simply through geometry, $\partial_t y = \partial_t(h/\sin\theta)$. Finally, the motor unbinds stochastically with a rate $\omega_{\rm off}$ when ATP displaces the residual ADP (both processes are combined into a single step here).

Finally, in the radial direction, we have the following boundary condition for the fluid pressure,

$$-\frac{\ell_p^2 K_\perp}{\eta}\partial_r p(R) = K_s\left[p(R) - p_\infty\right] , \tag{S10}$$

where $K_s$ is the surface permeability characterizing the net fluid flux for a given pressure drop across the surface and $p_\infty$ is the ambient pressure that we shall set to zero ($p_\infty = 0$) without loss of generality. For finite $K_s$, we have a nonzero pressure drop across the outer surface (the sarcolemma) that drives fluid flow relative to the solid motion. The surface is freely draining in the limit $K_s \to \infty$, corresponding to the typical experimental situation of studying permeabilized or glycerinated muscle fibres that can freely exchange fluid with an external bath. The opposite limit of jacketed boundary conditions, i.e., $K_s = 0$, corresponds to an impermeable membrane with no efflux, as is relevant for the physiological case of cells with an intact sarcolemma *in-vivo*.

##### B. Molecular motor kinetics

The active stress $\boldsymbol{\sigma}^a$ captures the effect of collective force generation by the molecular motors. On general symmetry grounds, we expect $\boldsymbol{\sigma}^a$ to be uniaxially anisotropic (due to the anisotropy of muscle fibers) and proportional to both the density of motors and the force generated per motor.

To compute the active stress $\boldsymbol{\sigma}^a$, we need a model for the mechanochemical kinetics of actomyosin that provide the molecular basis of contractility in muscle. A simple minimal description of the biochemical reaction is the Lymn-Taylor cycle [S27], which consists of four steps -

1. A detached myosin motor head, complexed with ADP and Pi binds to an available binding site on the actin thin filament. In striated muscle, this binding step is regulated by calcium ions ($Ca^{+2}$) which are required to first bind with a protein complex (troponin-tropomyosin) wound around the actin filament, in order to free up a binding site for myosin [S28]. We shall assume that $Ca^{+2}$ is present in excess and is not rate limiting (hence neurogenic control is neglected), which is the case for asynchronous flight muscle [S29].

2. The bound myosin releases the inorganic phosphate group (Pi) in the fastest step of the cycle and undergoes the power stroke. As a result, the motor changes conformation and develops an active contractile force.

3. The motor head unbinds post-power stroke when the ADP (remaining hydrolysis product) slowly releases and is displaced by a fresh ATP molecule.

4. Finally, the unbound motor head + ATP undergoes a hydrolysis reaction (ATP$\to$ ADP + Pi) that reprimes the motor to bind and perform the power-stroke once again.

Here, we shall combine the repriming step and the myosin binding step into one (with an overall rate $\omega_{\rm on}$), for simplicity. A sketch of the kinetic model is given in Fig. S1.

Let $\rho_b(\mathbf{x}, t)$ be the probability density of myosin motor heads being bound at position $\mathbf{x}$ and time $t$, $\rho_m(y; \mathbf{x}, t)$ be the probability density of bound active motors (i.e., after undergoing the power stroke) with an extension $y(t)$ and $\rho_u = 1 - \rho_b - \int {\rm d}y\ \rho_m$ is the density of unbound motors. The dynamics is prescribed by simple rate equations (shown pictorially in Fig. S1)

$$\partial_t \rho_b = -\omega_1 \rho_b + \omega_{\rm on} \rho_u\ , \tag{S11}$$

$$\partial_t \rho_m + \partial_y(\dot{y}\rho_m) = \omega_1 \rho_b \delta(y - y_0) - \omega_{\rm off} \rho_m\ . \tag{S12}$$

During the power stroke the motor head generates a prestrain $y = y_0 > 0$. To simplify the description even further, we note that the power stroke is the fastest step in the reaction cycle ($\omega_1 \sim 10^3\ {\rm s}^{-1}$) and on time scales longer than $\omega_1^{-1}$, we can neglect $\partial_t \rho_b \approx 0$ and write $\rho_b \simeq \rho_u \omega_{\rm on}/\omega_1$. This leads to the simpler model

$$\partial_t \rho_m + \partial_y(\dot{y}\rho_m) = \frac{\omega_{\rm on}}{1 + (\omega_{\rm on}/\omega_1)} (1 - n_m)\delta(y - y_0) - \omega_{\rm off} \rho_m\ , \tag{S13}$$

where $n_m = \int {\rm d}y\ \rho_m$ is the total density of bound myosin and the effective binding rate is now $\omega_{\rm on}/[1+(\omega_{\rm on}/\omega_1)] \simeq \omega_{\rm on}$ as $\omega_1 \gg \omega_{\rm on}$. In any case, we just have an effective binding rate that we shall continue to simply call $\omega_{\rm on}$. Kinetic models such as Eq. S13 have been used previously to describe the collective organization of molecular motors in a variety of settings [S30–S36].

When bound to actin, the motor extension $y$ evolves simply geometrically as $\partial_t y = \partial_t(h/\sin\theta)$, where we have assumed a simple binding geometry of the motor with an angle $\theta$ and the spacing between thick and thin filaments as $h$ (Fig. S1) [S37, S38]. For small deformations, we can write $h = h_0(1+\epsilon_\perp/2)$ (noting isotropy in the plane orthogonal to $z$) where $h_0 \sim 11-24$ nm is the equilibrium spacing between filaments [S26, S39]. Similarly, the longitudinal strain enters by noting the change in the horizontal projection $\ell = h/\tan\theta = \ell_0(1 + \epsilon_{zz})$ with $\ell_0 = h_0/\tan\theta_0$, which gives a linearized shear rotation of the crossbridge $\theta - \theta_0 = (\epsilon_\perp - 2\epsilon_{zz})\sin(2\theta_0)/4$ ($\theta_0$ being the equilibrium binding angle and $|\theta - \theta_0| \ll 1$). To linear order in strain, the crossbridge strain dynamics is then given by

$$\partial_t y = \frac{h_0}{\sin\theta_0}\left(\cos^2\theta_0\ \partial_t\epsilon_{zz} + \frac{1}{2}\sin^2\theta_0\ \partial_t\epsilon_\perp\right)\ . \tag{S14}$$

Note that here we do not include force dependent walking of the motor as muscle myosin II is a non-processive motor [S40].

The crossbridge is assumed to behave as a linear spring with a finite stiffness $k_m \sim 1-3$ pN/nm [S41–S43]. Once bound, the motors then generate an average force density

$$F_m = -k_m \int_0^\infty {\rm d}y\ \rho y\ , \tag{S15}$$

that is exerted on the filaments. Note that, at the initiation of the power stroke upon the phosphate release, $y = y_0 > 0$ and the force generated by a single motor head is $f_m = -k_m y_0 < 0$, which contracts the sarcomeric filaments as expected.

Finally, biophysical feedback is encoded via the kinetic rates that depend on the filament spacing $h$ and the crossbridge strain $y$. While many different microscopic mechansims of feedback are possible, here we consider the simplest setting. The binding rate $\omega_{\rm on}$ is chosen to depend on the spacing between the filaments $h = h_0(1 + \epsilon_\perp/2)$ simply for geometric reasons [S37, S38, S44–S46]. For a larger spacing, the probability of binding decreases such that $\omega_{\rm on} \propto e^{-k_m(h-d_m)^2/(k_BT)}$, so we have,

$$\omega_{\rm on}(\epsilon_\perp) = \omega_{\rm on}^0\ \exp\left[-\frac{k_m h_0^2}{k_BT}\left(1 - \frac{d_m}{h_0} + \frac{\epsilon_\perp}{4}\right)\epsilon_\perp\right]\ , \tag{S16}$$

where $\omega_{\rm on}^0$ is a base binding rate, $d_m(< h_0)$ is the equilibrium size of the motor head and $T$ is temperature ($k_B$ is Boltzmann's constant). While the mechanism of sensing filament spacing is geometric in our model, other mechanisms including elastic proteins such as titin [S47] can be subsumed within an effective strain dependent $\omega_{\rm on}(\epsilon_\perp)$.

On the other hand, the unbinding rate $\omega_{\rm off}$ depends on the local load as observed in single molecule experiments [S43, S48, S49]. We shall use simple thermodynamic arguments to obtain the form of mechanical feedback present. If $f_m$ is the force exerted by the motor on the filaments (so $-f_m$ is the load on the motor), the average internal energy of the crossbridge $\Delta U = \Delta U_0 - W$ includes the work done by the contractile force through the lever arm movement (the motor step) over a displacement $a \sim 1-3$ nm [S43, S48], given by $W = f_m a$. Using $f_m = -k_m y$ as the force generated by a single motor along with general thermodynamic considerations of Kramer's theory of barrier crossing, we can write a simple Arrhenius dependence for the unbinding rate as

$$\omega_{\rm off}(y) = \omega_{\rm off}^0 \exp\left[-\frac{a f_m}{k_B T}\right] = \omega_{\rm off}^0 \exp\left[\frac{k_m a}{k_B T} y\right] , \tag{S17}$$

where $\omega_{\rm off}^0$ is a base unbinding rate. For $a > 0$ as we have chosen here, this represents a conventional slip bond wherein motors unbind more frequently when they are more elongated and strained. Similar feedback models have been used previously to describe oscillatory instabilities in models of muscular contraction [S41, S50] and other motor filament assemblies [S31–S36]. Note that as $f_m$ is the force on the filaments, a tensile force $f_m > 0$ corresponds to filaments being stretched and sliding apart, which recruits more myosin (by supressing unbinding) to stabilize the sliding filament array. This behavior is consistent with observations of stretch response in muscle fibers [S51] and minimally recapitulates the well known physiological response of stretch activation in muscle [S52]. Nonetheless, the load dependence of actomyosin kinetics is known to be more complex, in particular, myosin also exhibits catch bond behavior at small strains/loads [S48, S49], i.e., when unbinding is suppressed by increasing load on the motor. This would correspond to $a < 0$ in Eq. S17, atleast within the window of catch bond behavior. One can show that, while catch bonding enhances the residence time of motors and can lead to clustering or collapse instabilities, it does not develop oscillatory instabilites within the simplified two-state model we use. But multistate models of actomyosin binding that account for intermediate conformation changes can support oscillatory instabilities with catch bond behavior [S32, S34, S50]. For simplicity, we shall assume the simple dependence in Eq. S17 and leave a more detailed analysis of multistage kinetic models for the future.

While Eq. S13 is the complete kinetic description, it is useful to work in the mean-field setting appropriate when the number of interacting motors is vary large. By integrating Eq. S13 over $y$, we define the average density of bound motors $n_m$ and the average crossbridge strain $\langle y\rangle$ as

$$n_m(\mathbf{x},t) = \int_0^\infty \mathrm{d}y\; \rho_m(\mathbf{x},t;y) , \quad \langle y\rangle(\mathbf{x},t) = \frac{1}{n_m(\mathbf{x},t)}\int_0^\infty \mathrm{d}y\; y\rho_m(\mathbf{x},t;y) , \tag{S18}$$

and obtain their coupled dynamics in the mean-field limit ($\langle\omega_{\rm off}(y)\rangle \approx \omega_{\rm off}(\langle y\rangle)$) to be

$$\partial_t n_m = \omega_{\rm on}(\epsilon_\perp)(1-n_m) - \omega_{\rm off}(\langle y\rangle) n_m , \tag{S19}$$

$$\partial_t\langle y\rangle = y_0\left(\lambda_\parallel\, \partial_t\epsilon_{zz} + \lambda_\perp\, \partial_t\epsilon_\perp\right) - \omega_{\rm on}(\epsilon_\perp)\left(\frac{1-n_m}{n_m}\right)(\langle y\rangle - y_0) . \tag{S20}$$

where we have defined the nondimensional geometric coefficients

$$\lambda_\parallel = \frac{h_0\cos^2\theta_0}{y_0\sin\theta_0} , \quad \lambda_\perp = \frac{h_0}{2y_0}\sin\theta_0 . \tag{S21}$$

where $\ell_s$ is the resting length of a single sarcomere and $h_0$ is the equilibrium spacing between the thick and thin filaments respectively. Similar equations have also been derived to describe collective dynamics of processive motors in flagella and cilia [S36].

With this molecular kinetic description in hand, we can construct the active stress. Each active crossbridge at position $\mathbf{x}_i$ behaves as a local force dipole oriented along the unit vector $\hat{\boldsymbol{\nu}}_i = (\sin\theta_i\cos\Phi_i, \sin\theta_i\sin\Phi_i, \cos\theta_i)$, where $\theta_i$ is the binding angle and $\Phi_i$ is the azimuthal angle of the $i^{th}$ motor head around a thick filament. The total active force density is given by

$$\begin{aligned}\mathbf{F}^a(\mathbf{x}) &= \sum_i f_m\left[\delta\left(\mathbf{x}-\mathbf{x}_i-\frac{d_m}{2}\hat{\boldsymbol{\nu}}_i\right) - \delta\left(\mathbf{x}-\mathbf{x}_i+\frac{d_m}{2}\hat{\boldsymbol{\nu}}_i\right)\right]\\ &= -\boldsymbol{\nabla}\cdot\left[\sum_i f_m d_m\hat{\boldsymbol{\nu}}_i\hat{\boldsymbol{\nu}}_i\delta(\mathbf{x}-\mathbf{x}_i)\right] + \mathcal{O}(d_m^2\nabla^2) ,\end{aligned} \tag{S22}$$

where we have performed a gradient expansion on scales much larger than the crossbridge size $d_m$. Upon assuming that $\Phi_i$ are distributed uniformly in $[0, 2\pi]$ (helical periodicity of myosin heads can also be included easily with minor quantitative changes) and each $\theta_i \simeq \theta_0$ (the average binding angle), we can write $\mathbf{F}^a = \boldsymbol{\nabla} \cdot \boldsymbol{\sigma}^a$ to obtain the well known active stress [S1, S53]

$$\boldsymbol{\sigma}^a = -N d_m F_m \left[\cos^2\theta_0 \hat{\mathbf{z}}\hat{\mathbf{z}} + \frac{1}{2}\sin^2\theta_0(\mathbf{I} - \hat{\mathbf{z}}\hat{\mathbf{z}})\right] , \tag{S23}$$

where $N$ is the linear density of myosin heads along the thick filament and $F_m = -k_m n_m \langle y \rangle$ (Eq. S15). As expected, $\boldsymbol{\sigma}^a$ is proportional to the motor density $n_m$, the motor strain $\langle y \rangle$ (which enter the force $F_m$) and is uniaxially anisotropic.

Eqs. S19 and S20 add two new dynamic variables describing the motor density $n_m$ and the active motor strain $\langle y \rangle$ (equivalently the active stress) which supplement the equations of poroelasticity (Eqs. S4, S5, S6) to complete our model description of muscle. We display the complete model here once again for convenience,

$$\boldsymbol{\nabla} \cdot \left[\phi(\boldsymbol{\sigma}^a + \boldsymbol{\sigma}^{el}) - (1-\phi)p\mathbf{1}\right] = \mathbf{0} , \tag{S24a}$$

$$(1-\phi)(\mathbf{v} - \partial_t \mathbf{u}) = -\frac{\ell_p^2}{\eta}\mathbf{K} \cdot \boldsymbol{\nabla} p , \tag{S24b}$$

$$\boldsymbol{\nabla} \cdot [\phi \partial_t \mathbf{u} + (1-\phi)\mathbf{v}] = 0 , \tag{S24c}$$

$$\partial_t n_m = \omega_{\text{on}}(\epsilon_\perp)(1-n_m) - \omega_{\text{off}}(\langle y \rangle) n_m , \tag{S24d}$$

$$\partial_t \langle y \rangle = y_0 \left(\lambda_\parallel \, \partial_t \epsilon_{zz} + \lambda_\perp \, \partial_t \epsilon_\perp\right) - \omega_{\text{on}}(\epsilon_\perp)\left(\frac{1-n_m}{n_m}\right)(\langle y \rangle - y_0) . \tag{S24e}$$

Constitutive relations determine the active stress $\boldsymbol{\sigma}^a = \boldsymbol{\sigma}^a(n_m, \langle y \rangle)$ (Eq. S23) and the elastic stress $\boldsymbol{\sigma}^{el} = \boldsymbol{\sigma}^{el}(\mathbf{u})$ (Eq. S2). Eq. S24 along with the boundary conditions quoted in Eqs. S9, S10 completes the full description of muscle as an active poroelastic material.

#### II. NONDIMENSIONALIZATION AND LINEARIZED DYNAMICS

For a long, slender muscular filament/fiber, we can assume axisymmetry and set $u_\varphi = 0$ and $\partial_\varphi = 0$. At steady state, we allow for some uniform background strain with $u_r = \epsilon_\perp^0 r/2$ and $u_z = \epsilon_\parallel^0 z$ which will depend on the boundary conditions imposed, e.g., if the boundaries are stress free. The average extension is $\langle y \rangle = y_0$ and the force exerted is the average stall force $F_m = -k_m y_0 n_m^0 \equiv -F_{\text{stall}}$ with the fraction of active bound motors determined by

$$n_m^0 = \frac{\omega_{\text{on}}(\epsilon_\perp^0)}{[\omega_{\text{on}}(\epsilon_\perp^0) + \omega_{\text{off}}(y_0)]} . \tag{S25}$$

In polar coordinates, force balance (Eq. S4) is given by

$$\partial_r \sigma_{rr} + \frac{1}{r}(\sigma_{rr} - \sigma_{\varphi\varphi}) + \partial_z \sigma_{rz} = 0 , \tag{S26}$$

$$\partial_r \sigma_{rz} + \frac{\sigma_{rz}}{r} + \partial_z \sigma_{zz} = 0 , \tag{S27}$$

where $\boldsymbol{\sigma} = \phi(\boldsymbol{\sigma}^{el} + \boldsymbol{\sigma}^a) - (1-\phi)p\mathbf{1}$ is the total stress. Upon neglecting variations in the solid fraction $\phi$ and using the constitutive relations (Eq. S2), we obtain

$$B\partial_r(\epsilon_\perp + \epsilon_{zz}) + G\left[\partial_r(\epsilon_{rr} - \epsilon_{\varphi\varphi}) + 2\frac{(\epsilon_{rr} - \epsilon_{\varphi\varphi})}{r} + 2\partial_z \epsilon_{rz}\right] + \partial_r \sigma_{rr}^a = \left(\frac{1-\phi}{\phi}\right)\partial_r p , \tag{S28}$$

$$(Y+B)\partial_z \epsilon_{zz} + B\partial_z \epsilon_\perp + 2G\left(\partial_r \epsilon_{rz} + \frac{\epsilon_{rz}}{r}\right) + \partial_z \sigma_{zz}^a = \left(\frac{1-\phi}{\phi}\right)\partial_z p . \tag{S29}$$

Here, we have written $B_\perp \equiv B$, $G_\perp = G_z \equiv G$ and $Y_z \equiv Y$. Of course, $B$, $G$ and $Y$ aren't independent (Eq. S3), but it is convenient to write them separately to ease the notation here. We can write this in terms of the displacements to get

$$(B+G)\left(\mathcal{L} - \frac{1}{r^2}\right)u_r + G\partial_z^2 u_r + (B+G)\partial_r \partial_z u_z + \partial_r \sigma_{rr}^a = \left(\frac{1-\phi}{\phi}\right)\partial_r p , \tag{S30}$$

$$G\mathcal{L}u_z + (Y+B)\partial_z^2 u_z + (B+G)\partial_z\left(\partial_r u_r + \frac{u_r}{r}\right) + \partial_z \sigma_{zz}^a = \left(\frac{1-\phi}{\phi}\right)\partial_z p , \tag{S31}$$

where the radial Laplacian is $\mathcal{L} = \partial_r^2 + (1/r)\partial_r$. We nondimensionalize space by the length ($L$) and radius ($R$) of the cylinder, time by the average residence time $\tau_k = [\omega_{\rm on}(\epsilon_\perp^0) + \omega_{\rm off}(y_0)]^{-1}$ of the bound myosin under stall conditions, the myosin extension by the prestrain ($y_0$), the force by the stall force $F_{\rm stall}$ and the pressure by the elastic moduli. This is given as

$$r = R\tilde{r}\ , \quad z = L\tilde{z}\ , \quad u_r = R\tilde{u}_r\ , \quad u_z = L\tilde{u}_z\ , \quad t = \tau_k\tilde{t}\ , \tag{S32}$$

$$F_m = F_{\rm stall}\tilde{F}\ , \quad p = \frac{\phi(B+G)\tilde{p}}{(1-\phi)}\ , \quad \langle y\rangle = \tilde{y}y_0\ , \quad n_m = n_m^0\tilde{n}\ . \tag{S33}$$

Dropping the tildes, we now write the nondimensionalized equations (Eqs. S24) as

$$\partial_t n = \frac{\tau_k\omega_{\rm on}(\epsilon_\perp)}{n_m^0} - \tau_k\left[\omega_{\rm on}(\epsilon_\perp) + \omega_{\rm off}(y)\right]n\ , \tag{S34}$$

$$\partial_t y = \lambda_\parallel\partial_t\epsilon_{zz} + \lambda_\perp\partial_t\epsilon_\perp - \frac{\tau_k\omega_{\rm on}(\epsilon_\perp)}{n_m^0}\frac{(1-n_m^0 n)}{n}(y-1)\ , \tag{S35}$$

$$\left[\mathcal{L} - \frac{1}{r^2} + a^2c_\perp\partial_z^2\right]u_r + \partial_r\partial_z u_z + \alpha_\perp\partial_r\sigma_a = \partial_r p\ , \tag{S36}$$

$$\frac{1}{a^2}\left[c_\perp\mathcal{L} + a^2c_\parallel\partial_z^2\right]u_z + \partial_z\left(\partial_r u_r + \frac{u_r}{r}\right) + \alpha_\parallel\partial_z\sigma_a = \partial_z p\ , \tag{S37}$$

$$\left[\partial_z^2 + \frac{K_r}{a^2}\mathcal{L}\right]p = {\rm Da}_\parallel\ \partial_t(\boldsymbol{\nabla}\cdot\mathbf{u})\ . \tag{S38}$$

Note the $y$ used henceforth is the nondimensionalized average crossbridge strain ($\langle y\rangle/y_0$) and should not be confused with the stochastic individual motor extension used previously. The aspect ratio of the muscle is $a = R/L < 1$ and the nondimensionalized active stress enters as $\sigma_a = ny$. The nondimensional activities $\alpha_{\perp,\parallel} > 0$, relative permeability $K_r$ and porosity dependent functions $c_{\parallel,\perp}(\phi) > 1$ are given as

$$\alpha_\parallel = \frac{Nd_mk_my_0n_m^0}{(B+G)}\cos^2\theta_0\ , \quad \alpha_\perp = \frac{Nd_mk_my_0n_m^0}{2(B+G)}\sin^2\theta_0\ , \tag{S39}$$

$$c_\parallel(\phi) = \frac{(2-\phi)(4-3\phi)}{3-2\phi}\ , \quad c_\perp(\phi) = \frac{1-\phi}{3-2\phi}\ , \quad K_r = \frac{K_\perp}{K_\parallel}\ . \tag{S40}$$

Note that for a dense solid with vanishing porosity ($\phi \to 1$), $c_\parallel \to 1$, and $c_\perp \simeq 1-\phi \to 0$. This is expected as $c_\parallel = (Y+B)/(B+G)$ and $c_\perp = G/(B+G)$ and $B \to \infty$ with $Y$ and $G$ finite when $\phi \to 1$. In practice, $\phi \lesssim \phi_* \approx 0.907$, the close packing limit at which the composite material becomes radially impermeable ($K_\perp \to 0$), and the bulk and shear moduli remain perfectly finite. In our model, the crossbridge geometry is assumed to be homogeneous and it controls both the reative strength of active forces and the strain dependent elongation of the motor in the same fashion, i.e., $\lambda_\perp/\lambda_\parallel = \alpha_\perp/\alpha_\parallel = \tan^2\theta_0/2$. If the binding angle were heterogeneous, as is the case in real muscle, then $\lambda_\perp/\lambda_\parallel \neq \alpha_\perp/\alpha_\parallel$. In particular $\lambda_\perp\alpha_\parallel/(\lambda_\parallel\alpha_\perp) = \langle\sin\theta\rangle\langle\cos^2\theta\rangle/\langle\cos\theta\cot\theta\rangle\langle\sin^2\theta\rangle \approx 1 - (2/\sin^2\theta_0)\langle(\theta-\theta_0)^2\rangle$, where $\langle\theta\rangle = \theta_0$. More general binding geometries also allow $\lambda_\perp/\lambda_\parallel \neq \alpha_\perp/\alpha_\parallel$.

The final parameter is the *poroelastic Damköhler number* that provides a measure of the relative rate of myosin kinetics ($\tau_k^{-1}$) to the rate of fluid permeation ($\tau_p^{-1}$). The anisotropic structure of muscle allows for two distinct Damköhler numbers, one characterizing axial transport (${\rm Da}_\parallel$) with $\tau_p^\parallel = [\eta/\phi(B+G)K_\parallel](L/\ell_p)^2$ and another for radial transport (${\rm Da}_\perp$) with $\tau_p^\perp = [\eta/\phi(B+G)K_\perp](R/\ell_p)^2$,

$${\rm Da}_\parallel = \frac{\tau_p^\parallel}{\tau_k} = \frac{\eta L^2}{\phi(B+G)K_\parallel\ell_p^2}[\omega_{\rm on}(\epsilon_\perp^0) + \omega_{\rm off}(y_0)]\ , \tag{S41}$$

$${\rm Da}_\perp = \frac{\tau_p^\perp}{\tau_k} = \frac{\eta R^2}{\phi(B+G)K_\perp\ell_p^2}[\omega_{\rm on}(\epsilon_\perp^0) + \omega_{\rm off}(y_0)]\ . \tag{S42}$$

Note, ${\rm Da}_\perp/{\rm Da}_\parallel = a^2/K_r \ll 1$ for $a = R/L \lesssim 0.1$ ($K_r = K_\perp/K_\parallel \sim 0.1$ for $\phi \sim 0.1-0.2$). When ${\rm Da}_{\parallel,\perp} \gg 1$, fluid permeation becomes rate limiting and the myosin binding kinetics occurs rapidly on the time scales of fluid transport. In the converse limit of ${\rm Da}_{\parallel,\perp} \ll 1$, kinetics dominates fluid transport and porous flow rapidly equilibriates hydraulic pressure throughout the muscle cell while contractility dynamics is slower.

We now linearize about the homogeneous steady state and write $n = 1 + \delta n$, $y = 1 + \delta y$, $\epsilon_\perp = \epsilon_\perp^0 + \delta\epsilon_\perp$, $\epsilon_{zz} = \epsilon_{zz}^0 + \delta\epsilon_{zz}$. We can expand the kinetic rates to linear order to get

$$\omega_{\rm on} = \omega_{\rm on}(\epsilon_\perp^0)[1 - \kappa\delta\epsilon_\perp] \;, \quad \omega_{\rm off} = \omega_{\rm off}(y_0)[1 + \beta\delta y] \;, \tag{S43}$$

where $\kappa = |\omega'_{\rm on}(\epsilon_\perp^0)|/\omega_{\rm on}(\epsilon_\perp^0) > 0$ and $\beta = y_0\omega'_{\rm off}(y_0)/\omega_{\rm off}(y_0) > 0$ are both positive couplings capturing mechanical feedback through both geometry and energetics. The linearized kinetic equations (Eqs. S34, S35) are

$$\partial_t\delta n = -\delta n - \beta f\delta y - \kappa f\delta\epsilon_\perp \;, \tag{S44}$$
$$\partial_t\delta y = -f\delta y + \lambda_\parallel\partial_t\delta\epsilon_{zz} + \lambda_\perp\partial_t\delta\epsilon_\perp \;. \tag{S45}$$

Here $f = 1 - n_m^0$ is the fraction of unbound myosin under stall conditions.

We retain the time dependence explicitly and write all the linearized equations in terms of the nondimensional active stress $\delta\sigma_a = \delta n + \delta y$ and the strains $\delta\epsilon_{zz}$ and $\delta\epsilon_\perp$. The force balance and pressure relaxation equations (Eqs. S36, S37, S38) can be rewritten as

$$\left[\mathcal{L} + c_\perp a^2\partial_z^2\right]\delta\epsilon_\perp + \mathcal{L}\delta\epsilon_{zz} + \alpha_\perp\mathcal{L}\delta\sigma_a = \mathcal{L}\delta p \;, \tag{S46}$$
$$\frac{1}{a^2}\left[c_\perp\mathcal{L} + c_\parallel a^2\partial_z^2\right]\delta\epsilon_{zz} + \partial_z^2\delta\epsilon_\perp + \alpha_\parallel\partial_z^2\delta\sigma_a = \partial_z^2\delta p \;, \tag{S47}$$
$$\left[\partial_z^2 + \frac{K_r}{a^2}\mathcal{L}\right]\delta p = {\rm Da}_\parallel\;\partial_t(\delta\epsilon_{zz} + \delta\epsilon_\perp) \;. \tag{S48}$$

#### III. OSCILLATORY INSTABILITY

Here we perform a stability analysis of the linearized equations above (Eqs. S44-S48) in different simplifying limits. We first take advantage of the slender geometry of muscle fibers and derive different reduced order models in 1D, and discuss the various regimes where molecular kinetics dominates and where active hydraulics dominates. At the end, we also provide a calculation of oscillatory dynamics that occurs when fully 3D deformations and spatial gradients are present and important.

##### A. 1D model with zero transverse strain ($|\epsilon_\perp| \ll |\epsilon_{zz}|$)

In the limit where axial deformations are larger than radial ones, we can neglect both transverse strains and radial gradients ($\epsilon_\perp = 0$, $\partial_r = 0$). Such a situation is appropriate for instance in *Drosophila* flight muscle that contracts with negligible changes in the filament lattice spacing [S54]. The linearized dynamics (Eq. S44-S48) then reduces to

$$\partial_t\delta n = -\delta n - \beta f\delta y \;, \tag{S49}$$
$$\partial_t\delta y = -f\delta y + \lambda_\parallel\partial_t\delta\epsilon_{zz} \;, \tag{S50}$$
$${\rm Da}_\parallel\;\partial_t\delta\epsilon_{zz} = \partial_z^2\left[c_\parallel\delta\epsilon_{zz} + \alpha_\parallel\delta\sigma_a\right] \;. \tag{S51}$$

This 1D description does not allow for strain cycles and neglects the free boundary condition ($\sigma_{rr}|_{r=1} = 0$) at the surface of the muscle fiber. The latter will necessitate radial gradients in a possibly small boundary layer near the surface that we neglect. Writing the above equations in terms of $\delta\sigma_a = \delta n + \delta y$, we obtain

$$\partial_t\delta n = (\beta f - 1)\delta n - \beta f\delta\sigma_a \;, \tag{S52}$$
$$\partial_t\delta\sigma_a = -(1+\beta)f\delta\sigma_a - [1 - (1+\beta)f]\delta n + \lambda_\parallel\partial_t\delta\epsilon_{zz} \;. \tag{S53}$$

When $\beta f > 1$ the growth rate of the motor density is positive ($\partial_t n/\delta n > 0$ for $\delta\sigma_a = 0$) and can be linearly unstable. For large Da, the axial strain relaxes slowly as well, while the active stress $\delta\sigma_a$ is the only variable that relaxes quickly on a finite time scale $\sim \tau_k/[f(1+\beta)]$. So at long times, we can set $\partial_t\delta\sigma_a = 0$ which yields

$$\delta\sigma_a \simeq \frac{[(1+\beta)f - 1]}{(1+\beta)f}\delta n + \frac{\lambda_\parallel}{(1+\beta)f}\partial_t\delta\epsilon_{zz} \;. \tag{S54}$$

This simplification is the basis of the calculation sketched in the main text. Here, we instead provide the complete solution retaining the dynamics of the active stress.

Assuming an exponential solution $\{\delta n, \delta y, \delta\epsilon_{zz}\} \sim e^{\mu t + i q_z z}$, and substituting into Eqs. S49-S51, we obtain a single cubic equation for the growth rate $\mu$

$$\mu^3 + A_2\mu^2 + A_1\mu + A_0 = 0 \,, \tag{S55}$$

$$A_0 = q_z^2 \frac{c_\parallel f}{\mathrm{Da}_\parallel} \,, \tag{S56}$$

$$A_1 = f + \frac{q_z^2}{\mathrm{Da}_\parallel}[c_\parallel(1+f) + \alpha_\parallel\lambda_\parallel(1-\beta f)] \,, \tag{S57}$$

$$A_2 = 1 + f + \frac{q_z^2}{\mathrm{Da}_\parallel}(c_\parallel + \alpha_\parallel\lambda_\parallel) \,. \tag{S58}$$

Note, the spatial mode number $q_z$ is dimensionless and for the longest wavelength mode we have $q_z = 2\pi$. As all the parameters in the model are defined to be positive, we have $A_0 > 0$ and $A_2 > 0$. $A_1$ is the only coefficient that can change sign if $\beta f > 1$ ($y_0\tau_k\omega'_{\mathrm{off}}(y_0) > 1$ in dimensionful terms) and if the activity $\alpha_\parallel$ is strong enough. Standard analysis based on the Routh-Hurwitz criterion [S55] dictates that $\mathrm{Re}[\mu] > 0$ (instability) iff $A_1 < 0$ or if $A_1A_2 < A_0$. In our case, the latter condition occurs first when increasing activity and we obtain a Hopf bifurcation when $A_1A_2 = A_0$. We can compute the threshold activity $\alpha_c$ for the longest wavelength mode to go unstable and the frequency of oscillations at onset by substituting $\mu = i\omega_c$ in Eq. S55 to get

$$\alpha_c = \frac{\mathrm{Da}_\parallel f + 4\pi^2 c_\parallel(1+f)}{4\pi^2\lambda_\parallel(\beta f - 1)} - \frac{c_\parallel f}{\lambda_\parallel[\beta f(1+f) - 1]} + \mathcal{O}\left(\frac{1}{\mathrm{Da}_\parallel}\right) \,, \tag{S59}$$

$$\omega_c = \sqrt{\frac{A_0}{A_2}} = 2\pi\sqrt{\frac{c_\parallel f}{4\pi^2(c_\parallel + \alpha_\parallel\lambda_\parallel) + \mathrm{Da}_\parallel(1+f)}} \propto \frac{1}{\sqrt{\mathrm{Da}_\parallel}} \,, \tag{S60}$$

where we have expanded out $\alpha_c$ for large $\mathrm{Da}_\parallel$ as physiologically relevant. Hence for $\alpha_\parallel > \alpha_c$ ($\beta f > 1$), we have spontaneous oscillations with frequency $\omega_c$. The scaling relation in Eq. S60 when restored to dimensionful form reads $\omega_c\tau_k \sim \mathrm{Da}_\parallel^{-1/2}$, which is a key result of our calculation. Using estimates of the relevant kinetic and poroelastic timescales for *Drosophila* (Tables I and II, with $\phi = 0.15$, $\lambda_\perp/\lambda_\parallel = \alpha_\perp/\alpha_\parallel = 0.5$), we obtain $\mathrm{Da}_\parallel \approx 2.06 \times 10^4$ and $\tau_k \approx 0.27$ ms [S56], which gives $\omega_c \approx 156$ Hz, a remarkably accurate prediction of the fruit fly wing-beat frequency!

##### B. 1D model with nonzero transverse strain ($|\epsilon_\perp| \sim |\epsilon_{zz}|$)

Here we retain both transverse and axial strains which are taken to be comparable in magnitude. The slender filamentous geometry of the muscle fiber still allows us to effectively derive a reduced order 1D model by considering the asymptotic limit $a = R/L \to 0$. We expand the variables in a regular power series as

$$u_z = u_z^{(0)} + a^2 u_z^{(1)} + \mathcal{O}(a^4) \,, \quad u_r = u_r^{(0)} + a^2 u_r^{(1)} + \mathcal{O}(a^4) \,, \quad p = p^{(0)} + a^2 p^{(1)} + \mathcal{O}(a^4) \,,$$
$$n = n^{(0)} + a^2 n^{(1)} + \mathcal{O}(a^4) \,, \quad \sigma_a = \sigma_a^{(0)} + a^2\sigma_a^{(1)} + \mathcal{O}(a^4) \,. \tag{S61}$$

Note that, as $a \to 0$, longitudinal and transverse strains remain $\mathcal{O}(1)$ and nonvanishing. The free stress ($\sigma_{rr}|_{r=1} = 0$, $\sigma_{rz}|_{r=1} = 0$) and no-flux ($\partial_r p|_{r=1} = 0$) boundary conditions (Eqs. S9, S10) can be similarly expanded to give

$$\partial_r u_z^{(0)}|_{r=1} = 0 \,, \quad \partial_r u_z^{(1)} + \partial_z u_r^{(0)}|_{r=1} = 0 \,, \tag{S62a}$$

$$\partial_r u_r^{(0)} + (1 - 2c_\perp)u_r^{(0)} + (1 - c_\perp)\epsilon_{zz}^{(0)} + \alpha_\perp\sigma_a^{(0)} - p^{(0)}|_{r=1} = 0 \,, \tag{S62b}$$

$$\partial_r p^{(0)}|_{r=1} = 0 \,, \quad \partial_r p^{(1)}|_{r=1} = 0 \,. \tag{S62c}$$

In such an expansion, two distinct cases are possible,

1. $\mathrm{Da}_\parallel$ is kept fixed (so $\mathrm{Da}_\perp \to 0$) as $a^2 \to 0$,

2. $\mathrm{Da}_\perp$ is kept fixed (so $\mathrm{Da}_\parallel \to \infty$) as $a^2 \to 0$.

Note, in the first case, as $\mathrm{Da}_\perp \to 0$ radial permeation offers negligible hydraulic resistance and fluid pressure equilibriates rapidly, essentially functioning as a radial shunt for easy fluid movement. In the second case, flow is hydraulically hindered in all directions, being blocked off entirely in the axial direction when $\mathrm{Da}_\parallel \to \infty$. The key point to note is that the fastest poroelastic time, i.e., the smallest Damköhler number controls the effective dynamics. In the presence of disorder and heterogeneity in the porous structure, we expect the same behaviour to hold, wherein the path of least hydraulic resistance (typically expected to be in the radial direction) controls the relevant (fastest) hydraulic timescale that can potentially compete with molecular kinetics.

**1. $\mathrm{Da}_{\parallel}$ fixed, $\mathrm{Da}_{\perp} \to 0$**

This limit is applicable when fluid flow occurs rapidly in the radial direction (so $\mathrm{Da}_{\perp} \ll 1$) and is consequently not rate limiting. As a result, here we expect that the frequency of spontaneous oscillations will be dominated by actomyosin kinetics. To see this in a systematic calculation, we consider the force balance and pressure dynamics equations (Eqs. S46-S48) along with boundary conditions to lowest order in $a^2$, which give

$$\mathcal{L}u_z^{(0)} = 0 \implies u_z^{(0)} = u_z^{(0)}(z,t) \ , \tag{S63}$$

$$\mathcal{L}p^{(0)} = 0 \implies p^{(0)} = p^{(0)}(z,t) \ , \tag{S64}$$

$$\left(\mathcal{L} - \frac{1}{r^2}\right) u_r^{(0)} + \partial_r \left[\alpha_\perp \sigma_a^{(0)} - p^{(0)}\right] = 0 \implies u_r^{(0)} = -\frac{r}{2}\epsilon_{zz}^{(0)} + \frac{r}{2(1-c_\perp)}[p^{(0)} - \alpha_\perp \sigma_a^{(0)}] \ , \tag{S65}$$

where in the last equation we have taken $\partial_r \sigma_a^{(0)} = 0$ for self-consistency purposes (as expected in a slender filament). The leading order transverse strain is then constant across the fiber cross-section and is obtained using Eq. S65 to be

$$\epsilon_\perp^{(0)} = -\epsilon_{zz}^{(0)} + \frac{p^{(0)} - \alpha_\perp \sigma_a^{(0)}}{(1-c_\perp)} \ . \tag{S66}$$

As anticipated, the deformation is affine across the cross-section of the fiber and transverse strains are instantaneously slaved to local changes in axial stretch, pressure and the active stress. Using this solution, the first order correction satisfies the following equations for the pressure and axial force balance (Eq. S46, S48),

$$\partial_z^2 p^{(0)} + K_r \mathcal{L} p^{(1)} = \mathrm{Da}_\parallel \, \partial_t \left(\epsilon_{zz}^{(0)} + \epsilon_\perp^{(0)}\right) \ , \tag{S67}$$

$$c_\perp \mathcal{L} u_z^{(1)} + \partial_z \left[c_\parallel \epsilon_{zz}^{(0)} + \epsilon_\perp^{(0)} + \alpha_\parallel \sigma_a^{(0)} - p^{(0)}\right] = 0 \ . \tag{S68}$$

The solvability condition is then simply obtained by integrating Eqs. S67, S68 over the radius ($\int_0^1 \mathrm{d}r \; r$), which gives

$$\partial_z^2 p^{(0)} = \mathrm{Da}_\parallel \, \partial_t \left(\epsilon_{zz}^{(0)} + \epsilon_\perp^{(0)}\right) \ , \tag{S69}$$

$$\partial_z \left[c_\parallel \epsilon_{zz}^{(0)} + \epsilon_\perp^{(0)} + \alpha_\parallel \sigma_a^{(0)} - p^{(0)}\right] = -2c_\perp \partial_r u_z^{(1)}|_{r=1} = 2c_\perp \partial_z u_r^{(0)}|_{r=1} \ . \tag{S70}$$

Upon eliminating $\epsilon_\perp^{(0)}$ using Eq. S66, dropping the (0) superscript and linearizing about the steady state ($n = 1 + \delta n$, $y = 1 + \delta y$, $p = \delta p$, $\boldsymbol{\epsilon} = \boldsymbol{\epsilon}^0 + \delta\boldsymbol{\epsilon}$, $\delta\sigma_a = \delta n + \delta y$), we finally obtain

$$\partial_z[(c_\parallel + c_\perp - 1)\delta\epsilon_{zz} + \Delta\alpha\delta\sigma_a] = 0 \ , \tag{S71}$$

$$\partial_t(\delta p - \alpha_\perp \delta\sigma_a) = \frac{(1-c_\perp)}{\mathrm{Da}_\parallel}\partial_z^2 \delta p \ , \tag{S72}$$

$$\partial_t \delta n = (\beta f - 1)\delta n - \beta f \delta\sigma_a + \kappa f \left(\delta\epsilon_{zz} - \frac{1}{\mathrm{Da}_\parallel}\partial_z^2 \delta p\right) \ , \tag{S73}$$

$$\partial_t \delta\sigma_a = -f(1+\beta)\delta\sigma_a + [(1+\beta)f - 1]\delta n + \Delta\lambda \, \partial_t \delta\epsilon_{zz} + \kappa f \delta\epsilon_{zz} + \frac{(\lambda_\perp - \kappa f)}{\mathrm{Da}_\parallel}\partial_z^2 \delta p \ . \tag{S74}$$

where $\Delta\alpha = \alpha_\parallel - \alpha_\perp > 0$, $\Delta\lambda = \lambda_\parallel - \lambda_\perp > 0$, and we have written the kinetic equations in terms of $\delta n$ and $\delta\sigma_a$. Note, if $\kappa = \lambda_\perp = 0$, pressure entirely decouples from the motor kinetics. Also, from Eq. S66 we can see that while oscillations in this parameter regime are not isochoric, they do not allow for substantial strain cycles as both $\epsilon_\perp$ and $\epsilon_{zz}$ vary in phase and are determined directly by the active stress (pressure plays only a minor role for large $\mathrm{Da}_\parallel$).

Assuming an exponential solution $\{\delta n, \delta y, \delta\epsilon_{zz}, \delta p\} \sim e^{\mu t + i q_z z}$, and substituting into Eqs. S71-S74, we obtain a single cubic equation for the growth rate $\mu$

$$\mu^3 + A_2\mu^2 + A_1\mu + A_0 = 0\ , \tag{S75}$$

$$A_0 = \frac{c_1(c_2 + \kappa f\Delta\alpha) f q_z^2}{\mathrm{Da}_\parallel (c_2 + \Delta\alpha\Delta\lambda)}\ , \tag{S76}$$

$$A_1 = \frac{f(c_2 + \kappa f\Delta\alpha)}{(c_2 + \Delta\alpha\Delta\lambda)} + \frac{q_z^2}{\mathrm{Da}_\parallel (c_2 + \Delta\alpha\Delta\lambda)}\left[c_1c_2(1+f) + c_1\Delta\alpha(\kappa f - \Delta\lambda(\beta f - 1)) - c_2\alpha_\perp(\kappa f^2 + \lambda_\perp(\beta f - 1))\right]\ , \tag{S77}$$

$$A_2 = \frac{[c_2(1+f) - \Delta\alpha(\Delta\lambda(\beta f - 1) - \kappa f)]}{(c_2 + \Delta\alpha\Delta\lambda)} + \frac{q_z^2}{\mathrm{Da}_\parallel}\left[c_1 + c_2\alpha_\perp\left(\frac{\lambda_\perp - \kappa f}{c_2 + \Delta\alpha\Delta\lambda}\right)\right]\ . \tag{S78}$$

Note $c_1 = 1 - c_\perp > 0$ and $c_2 = c_\parallel + c_\perp - 1 \geq 0$ for all solid fraction $\phi \in [0, 1]$, so $A_0 > 0$. When $\beta f > 1$, $A_1$ and $A_2$ can become negative for large enough activity. The relevant condition for the onset of an oscillatory instability is $A_1A_2 \leq A_0$. Upon writing $\alpha_\perp/\alpha_\parallel = \tan^2\theta_0/2$, we expand out the critical activity threshold for large $\mathrm{Da}_\parallel$

$$\Delta\alpha_c = \frac{c_2(1+f)}{[\Delta\lambda(\beta f - 1) - \kappa f]} - \frac{c_2^2 q_z^2 \tan^2\theta_0(1+f)(\kappa f - \lambda_\perp)}{(2 - \tan^2\theta_0)[\Delta\lambda(\beta f - 1) - \kappa f]^2\mathrm{Da}_\parallel} + \mathcal{O}\left(\frac{1}{\mathrm{Da}_\parallel^2}\right)\ , \tag{S79}$$

so that when $\Delta\alpha > \Delta\alpha_c$, we obtain spontaneous oscillations. In the limit in which $\kappa = 0$, $\lambda_\perp = 0$, we have $\Delta\alpha_c = c_2(1+f)/[\lambda_\parallel(\beta f - 1)]$ exactly. For $\theta_0 = 45^\circ$ and $q_z = 2\pi$, the activity threshold is given by

$$\alpha_c = \frac{2c_2(1+f)}{[\Delta\lambda(\beta f - 1) - \kappa f]} - \frac{8\pi^2 c_2^2(1+f)(\kappa f - \lambda_\perp)}{[\Delta\lambda(\beta f - 1) - \kappa f]^2\mathrm{Da}_\parallel} + \mathcal{O}\left(\frac{1}{\mathrm{Da}_\parallel^2}\right)\ . \tag{S80}$$

Note, this activity threshold only exists for sufficiently strong mechanical feedback, i.e., when $\Delta\lambda(\beta f - 1) > \kappa f$.

The oscillation frequency at threshold

$$\omega_c = \sqrt{\frac{A_0}{A_2}} = \sqrt{\frac{\Delta\lambda(\beta f - 1) + \kappa f^2}{\Delta\lambda(1+\beta) - \kappa}} + \mathcal{O}\left(\frac{1}{\mathrm{Da}_\parallel}\right)\ . \tag{S81}$$

is finite and independent of $\mathrm{Da}_\parallel$ to leading order, as expected of a kinetically dominated instability.

While the above calculation was performed for a fiber with an impermeable surface (representing an intact sarcolemma), a similar calculation is possible for a fully permeable membrane as present in glycerinated fibers. The fiber surface is then freely draining with the pressure boundary condition $p|_{r=1} = 0$. Using this, we instead obtain $p^{(0)} = 0$ and $2\partial_r p^{(1)}|_{r=1} = \mathrm{Da}_\parallel\partial_t(\epsilon_{zz}^{(0)} + \epsilon_\perp^{(0)})$, so pressure fluctuations are absent at leading order. Hence, the resulting equations only involve molecular kinetics and hydraulics plays no role in generating the instability with the threshold activity $\Delta\alpha_c = c_2(1+f)/[\Delta\lambda(\beta f - 1) - \kappa f]$ and oscillation frequency $\omega_c = \sqrt{[\Delta\lambda(\beta f - 1) + \kappa f^2]/[\Delta\lambda(1+\beta) - \kappa]}$.

**2. $\mathrm{Da}_\perp$ fixed, $\mathrm{Da}_\parallel \to \infty$**

Here we analyze a different limit, wherein the fiber has a *finite* hydraulic resistance in the radial direction, and a much larger (potentially infinite) resistance to fluid permeation in the axial direction. By keeping $\mathrm{Da}_\perp$ fixed and finite, fluid flow is not infinitely rapid, but instead takes a finite time, allowing the possibility of an interplay between hydraulics and kinetics.

Using the same expansion in Eq. S61, at lowest order in $a^2$, Eqs. S46-S48 take the form

$$\mathcal{L}u_z^{(0)} = 0 \implies u_z^{(0)} = u_z^{(0)}(z,t)\ , \tag{S82}$$

$$\mathcal{L}p^{(0)} = \mathrm{Da}_\perp\partial_t\left(\epsilon_{zz}^{(0)} + \epsilon_\perp^{(0)}\right)\ , \tag{S83}$$

$$\left(\mathcal{L} - \frac{1}{r^2}\right)u_r^{(0)} + \partial_r\left[\alpha_\perp\sigma_a^{(0)} - p^{(0)}\right] = 0\ . \tag{S84}$$

These can be solved with the boundary conditions in Eq. S62 to obtain

$$\epsilon_\perp^{(0)}(r,z,t) = -\epsilon_{zz}^{(0)}(z,t) + E_q(z,t)J_0(q_\perp r)\ , \tag{S85}$$

$$p^{(0)}(r,z,t) = P_0(z,t) + P_q(z,t)J_0(q_\perp r)\ , \tag{S86}$$

$$\sigma_a^{(0)}(r,z,t) = \Sigma_0(z,t) + \Sigma_q(z,t)J_0(q_\perp r)\ , \tag{S87}$$

where $E_q$, $P_q$, $\Sigma_q$ are undetermined functions of $(z,t)$ and $J_n(x)$ is a Bessel function and $q_\perp \approx 3.832$ is determined by the first nonzero root of $J_1(q_\perp) = 0$ (assuming the longest wavelength mode in the radial direction). As a reminder, we note some simple identities of Bessel functions - $\mathcal{L}J_0(qr) = -q^2 J_0(qr)$, $\partial_r J_0(qr) = -qJ_1(qr)$ and $[\partial_r + (1/r)]J_1(qr) = qJ_0(qr)$. The mode amplitudes satisfy

$$\mathrm{Da}_\perp \partial_t E_q + q_\perp^2 (E_q + \alpha_\perp \Sigma_q) = 0 \ , \quad P_0 = \alpha_\perp \Sigma_0 \ , \tag{S88}$$

while the radial dispacement is given by

$$u_r^{(0)} = \frac{r}{2}\left[-\epsilon_{zz}^{(0)} + \frac{E_q}{\Gamma(2)}\ {}_0F_1\left(2; -\frac{q_\perp^2 r^2}{4}\right)\right] \ , \tag{S89}$$

where ${}_0F_1(a;x)$ is a generalized hypergeometric function and $\Gamma(n)$ is the gamma function. Note, $P_0$ and $\Sigma_0$ are the radially averaged pressure and active stress respectively as $\int_0^1 \mathrm{d}r\ rJ_0(q_\perp r) = J_1(q_\perp)/q_\perp = 0$.

Axial force balance at next order gives

$$c_\perp \mathcal{L}u_z^{(1)} + \partial_z \left[c_\parallel \epsilon_{zz}^{(0)} + \epsilon_\perp^{(0)} + \alpha_\parallel \sigma_a^{(0)} - p^{(0)}\right] = 0 \ . \tag{S90}$$

Once again applying the solvability condition (here simply integrating as $\int_0^1 \mathrm{d}r\ r$) and noting that $J_1(q_\perp) = 0$ and ${}_0F_1(2; -q_\perp^2/4) = 0$, we obtain

$$\partial_z \left[(c_\parallel - 1)\epsilon_{zz}^{(0)} + \alpha_\parallel \Sigma_0 - P_0\right] = -2c_\perp \partial_r u_z^{(1)}|_{r=1} = 2c_\perp \partial_z u_r^{(0)}|_{r=1} \ . \tag{S91}$$

Upon writing $n^{(0)} = N_0 + N_q J(q_\perp r)$, dropping the (0) superscipt and linearizing about the steady state, we now have two separate sets of coupled dynamical equations, one for the dynamics in the axial direction and the other in the radial direction. In the axial direction, the average dynamics is dominated by molecular kinetics and following from Eqs. S91, S44, S45 we obtain

$$\partial_z \left[(c_\parallel + c_\perp - 1)\delta\epsilon_{zz} + \Delta\alpha\delta\Sigma_0\right] = 0 \ , \tag{S92}$$

$$\partial_t \delta N_0 = (\beta f - 1)\delta N_0 - \beta f \delta\Sigma_0 + \kappa f \delta\epsilon_{zz} \ , \tag{S93}$$

$$\partial_t \delta\Sigma_0 = -(1+\beta) f \delta\Sigma_0 + [(1+\beta)f - 1]\delta N_0 + \Delta\lambda \partial_t \delta\epsilon_{zz} + \kappa f \delta\epsilon_{zz} \ . \tag{S94}$$

As expected, fluid flow and pressure do not enter here at all. We can eliminate the active stress and strain fluctuations in favour of the motor density by differentiating Eq. S93 and substituting Eqs. S92, S94 to obtain an effective damped harmonic like oscillator in $\delta N_0$,

$$m\delta\ddot{N}_0 + \xi\delta\dot{N}_0 + k\delta N_0 = 0 \ , \tag{S95}$$

with an effective inertia $m$, damping $\xi$ and spring constant $k$ given by

$$m = c_2 + \Delta\alpha\Delta\lambda \ , \quad \xi = c_2(1+f) - \Delta\alpha\left[\Delta\lambda(\beta f - 1) - \kappa f\right] \ , \quad k = f(c_2 + \kappa f \Delta\alpha) \ , \tag{S96}$$

where $c_2 = c_\parallel + c_\perp - 1 \geq 0$ for all solid fraction $\phi \in [0,1]$. While $m > 0$ and $k > 0$ for all parameters, the effective damping $\xi$ can switch sign, becoming negative for large enough activity ($\Delta\alpha \geq \Delta\alpha_c$) when the feedback is strong enough ($\Delta\lambda(\beta f - 1) > \kappa f$). The threshold activity and oscillation frequency at onset are simply given as before

$$\Delta\alpha_c = \frac{c_2(1+f)}{[\Delta\lambda(\beta f - 1) - \kappa f]} \ , \quad \omega_c = \sqrt{\frac{k}{m}} = \sqrt{\frac{\Delta\lambda(\beta f - 1) + \kappa f^2}{\Delta\lambda(1+\beta) - \kappa}} \ . \tag{S97}$$

In the radial direction, the dynamics involves both molecular kinetics and hydraulics, so that Eqs. S88, S44, S45 become

$$\partial_t \delta N_q = (\beta f - 1)\delta N_q - \beta f \delta\Sigma_q - \kappa f \delta E_q \ , \tag{S98}$$

$$\partial_t \delta\Sigma_q = -(1+\beta) f \delta\Sigma_q + [(1+\beta)f - 1]\delta N_q + \lambda_\perp \partial_t \delta E_q - \kappa f \delta E_q \ , \tag{S99}$$

$$\partial_t \delta E_q = -\frac{q_\perp^2}{\mathrm{Da}_\perp}\left[\delta E_q + \alpha_\perp \delta\Sigma_q\right] \ , \tag{S100}$$

Note that the dynamics in the axial and radial directions are decoupled at leading order and cross-coupling terms appear only at higher order in $a^2$. This reduced set of dynamics is the radial analogue of the 1D model analyzed previously in Eqs. S49-S51. Assuming an exponential solution, $\{\delta E_q, \delta N_q, \delta\Sigma_q\} \sim e^{\mu t}$, and substituting in Eqs. S98-S100 we obtain a cubic equation for the growth rate

$$\mu^3 + A_2\mu^2 + A_1\mu + A_0 = 0\ , \tag{S101}$$

$$A_0 = \frac{q_\perp^2 f}{\mathrm{Da}_\perp}[1-\alpha_\perp\kappa f]\ , \tag{S102}$$

$$A_1 = f + \frac{q_\perp^2}{\mathrm{Da}_\perp}\left[1+f-\alpha_\perp\left(\lambda_\perp(\beta f-1)+\kappa f\right)\right]\ , \tag{S103}$$

$$A_2 = 1+f+\frac{q_\perp^2}{\mathrm{Da}_\perp}(1+\alpha_\perp\lambda_\perp)\ . \tag{S104}$$

The onset of an oscillatory instability is once again determined by the condition $A_1A_2 = A_0$, with the onset frequency given by $\omega_c = \sqrt{A_0/A_2}$. While we can obtain a general expression for the activity threshold and frequency, it is easiest to consider the case of $\kappa = 0$, where the expression simplifies considerably. The critical transverse activity above which we have spontaneous oscillations ($\alpha_\perp > \alpha_c^\perp$) and the corresponding frequency are easily obtained for large $\mathrm{Da}_\perp$ as

$$\alpha_c^\perp = \frac{\mathrm{Da}_\perp f + q_\perp^2(1+f)}{q_\perp^2\lambda_\perp(\beta f-1)} - \frac{f}{\lambda_\perp[\beta f(1+f)-1]} + \mathcal{O}\left(\frac{1}{\mathrm{Da}_\perp}\right)\ , \tag{S105}$$

$$\omega_c = q_\perp\sqrt{\frac{f}{q_\perp^2(1+\alpha_\perp\lambda_\perp)+\mathrm{Da}_\perp(1+f)}} \propto \frac{1}{\sqrt{\mathrm{Da}_\perp}}\ , \tag{S106}$$

which is the key scaling result of our work and demonstration of how active hydraulics crucially dictates the time-scale of repetitive muscular contractions. This result can be straightforwardly generalized for the case of $\kappa \neq 0$ as well, but the scaling with $\mathrm{Da}_\perp$ remains unchanged.

##### C. 3D oscillations with stable kinetics ($|\epsilon_\perp| \gg |\epsilon_{zz}|$)

Unlike the previous cases where the oscillations relied on a kinetic feedback mechanism being present, here we consider the opposite situation where oscillations arise purely from active mechanics in 3D without resorting to a kinetic instability. While we are currently unaware of any examples where muscle physiology may use such a mechanism, as we shall show below, spontaneous oscillations can develop from the emergent nonreciprocal active response present in our model, even with stable kinetics, similar to oscillations and waves seen in other odd elastic systems [S57, S58]. Crucially, these oscillations are unavailable in the reduced order 1D descriptions described earlier and they involve spatial gradients and strain deformations in 3D with transverse strains dominating axial ones in the slender fiber limit ($a \ll 1$).

We start with the linearized force balance and flow equations (Eq. S46-S48) and eliminate $\delta p$ in two different ways to obtain

$$\left[\mathcal{L} + \frac{a^2}{K_r}(1+c_\perp K_r)\partial_z^2\right]\delta\epsilon_\perp + \left[\left(1+\frac{c_\perp}{K_r}\right)\mathcal{L} + a^2\frac{c_\parallel}{K_r}\partial_z^2\right]\delta\epsilon_{zz} + \left[\alpha_\perp\mathcal{L} + \alpha_\parallel\frac{a^2}{K_r}\partial_z^2\right]\delta\sigma_a = \mathrm{Da}_\perp\ \partial_t(\delta\epsilon_{zz}+\delta\epsilon_\perp)\ , \tag{S107}$$

$$a^2c_\perp\partial_z^4\delta\epsilon_\perp + \frac{1}{a^2}\left[c_\perp\mathcal{L} - a^2(c_\parallel-1)\partial_z^2\right]\mathcal{L}\delta\epsilon_{zz} = \Delta\alpha\mathcal{L}\partial_z^2\delta\sigma_a\ . \tag{S108}$$

As before, $\Delta\alpha = \alpha_\parallel - \alpha_\perp > 0$ as the muscle fibres are more contractile in the longitudinal direction than in the transverse plane and $c_\parallel \geq 1$ for all solid fraction $\phi \in [0,1]$. We neglect the boundary conditions and simply assume the fields are well described using a single Fourier mode in space, i.e., $\{\delta\epsilon_{zz}, \delta\epsilon_\perp, \delta\sigma_a, \delta n\} \sim J_0(q_\perp r)e^{iq_z z}$ with (nondimensionalized) wavevectors $q_z$ and $q_\perp$ along the axial and radial directions respectively. This then gives the following compatibility like relation that follows from Eq. S108

$$\delta\epsilon_\perp = \Delta\alpha\ M_0\delta\sigma_a - \nu\delta\epsilon_{zz}\ , \tag{S109}$$

$$M_0 = \frac{1}{a^2c_\perp}\left(\frac{q_\perp}{q_z}\right)^2\ ,\quad \nu = \frac{1}{a^4}\left(\frac{q_\perp}{q_z}\right)^4\left[1 - a^2\frac{(c_\parallel-1)}{c_\perp}\left(\frac{q_z}{q_\perp}\right)^2\right]\ , \tag{S110}$$

where $M_0 > 0$ and the effective Poisson's ratio $\nu > 0$ for typical parameter values ($q_\perp/q_z \sim \mathcal{O}(1)$, $\phi \sim 0.1 - 0.2$, $a \lesssim 0.1$). Note $\nu$ is not a conventional Poisson's ratio as it depends on nonvanishing spatial gradients in strain, evidenced by the presence of $q_z$ and $q_\perp$. Furthermore, as $\nu \sim 1/a^4$, we immediately see that $|\delta\epsilon_\perp| \gg |\delta\epsilon_{zz}|$ as $a \to 0$, so transverse strains dominate axial ones here.

Using Eq. S109, the dynamics of volumetric deformations governed by Eq. S107 becomes

$$\mathrm{Da}_\perp\left[M_0\Delta\alpha\,\delta\dot{\sigma}_a + (1-\nu)\delta\dot{\epsilon}_{zz}\right] + (M_\parallel - \nu M_\perp)\delta\epsilon_{zz} + (M_\alpha + M_0 M_\perp\Delta\alpha)\delta\sigma_a = 0\ , \tag{S111}$$

where $M_{\parallel,\perp,\alpha} > 0$ all depend on $K_r$, $c_{\parallel,\perp}$, $a^2$ and the spatial wavevectors $q_{z,\perp}$. $M_\alpha$ also depends on activity through $\alpha_{\parallel,\perp}$. The expressions for $M_{\parallel,\perp,\alpha}$ are

$$M_\perp = q_\perp^2 + \frac{a^2}{K_r}(1 + c_\perp K_r)q_z^2\ , \tag{S112}$$

$$M_\parallel = \left(1 + \frac{c_\perp}{K_r}\right)q_\perp^2 + a^2\frac{c_\parallel}{K_r}q_z^2\ , \tag{S113}$$

$$M_\alpha = \alpha_\perp q_\perp^2 + \alpha_\parallel\frac{a^2}{K_r}q_z^2\ . \tag{S114}$$

The linearized motor binding kinetics (Eqs. S44, S45) along with Eq. S109 then gives for $\delta n$ and $\delta\sigma_a = \delta n + \delta y$ the following equations,

$$\delta\dot{n} = -(1-\beta f)\delta n - f(\beta + \kappa M_0\Delta\alpha)\delta\sigma_a + \nu\kappa f\delta\epsilon_{zz}\ , \tag{S115}$$

$$(1-\lambda_\perp M_0\Delta\alpha)\delta\dot{\sigma}_a = -\left[1 - f(1+\beta)\right]\delta n - f(1 + \beta + \kappa M_0\Delta\alpha)\delta\sigma_a + \nu\kappa f\delta\epsilon_{zz} + \left(\lambda_\parallel - \lambda_\perp\nu\right)\delta\dot{\epsilon}_{zz}\ . \tag{S116}$$

Here, we shall consider $\beta f < 1$, i.e., weak mechanical feedback on the binding kinetics. In the extreme limit of $\beta = 0$, the motor density relaxes on a fast time scale (here set to unity) and doesn't go unstable by itself. A necessary condition for a kinetic instability is $\beta f > 1$, so by prohibiting that, we focus instead on instabilities that arise from a combination of 3D contractility and poroelasticity. To emphasize the irrelevance of mechanical feedback, we shall work in the $\beta = 0$ limit (but retain geometric feedback with $\kappa > 0$) for simplicity. Then setting $\partial_t\delta n = 0$ and slaving $\delta n$ to the other variables, we get from Eq. S115

$$\delta n \simeq -f\kappa M_0\Delta\alpha\delta\sigma_a + \nu\kappa f\delta\epsilon_{zz}\ . \tag{S117}$$

This leads to the coupled dynamics of the active stress and the axial strain (from Eqs. S116, S111),

$$(1-\lambda_\perp M_0\Delta\alpha)\delta\dot{\sigma}_a + \left(\nu\lambda_\perp - \lambda_\parallel\right)\delta\dot{\epsilon}_{zz} = -f(1 + \kappa f M_0\Delta\alpha)\delta\sigma_a + \kappa\nu f^2\delta\epsilon_{zz}\ , \tag{S118}$$

$$\mathrm{Da}_\perp\ M_0\Delta\alpha\ \delta\dot{\sigma}_a - \mathrm{Da}_\perp\ (\nu-1)\delta\dot{\epsilon}_{zz} = (\nu M_\perp - M_\parallel)\delta\epsilon_{zz} - (M_\alpha + M_0 M_\perp\Delta\alpha)\delta\sigma_a\ . \tag{S119}$$

We can combine these two equations into a single second order in time equation for the active stress $\delta\sigma_a$, to get an effective description as a damped harmonic oscillator,

$$m\delta\ddot{\sigma}_a + \xi\delta\dot{\sigma}_a + k\delta\sigma_a = 0\ , \tag{S120}$$

with an effective mass $m$, friction $\xi$ and spring constant $k$ given by

$$m = \mathrm{Da}_\perp\left(\nu - \Delta\alpha\Delta\lambda M_0 - 1\right)\ , \tag{S121}$$

$$\xi = \nu M_\perp - M_\parallel + M_\alpha(\nu\lambda_\perp - \lambda_\parallel) - \Delta\alpha M_0(\lambda_\parallel M_\perp - \lambda_\perp M_\parallel) + \mathrm{Da}_\perp f\left(\nu - \kappa f\Delta\alpha M_0 - 1\right)\ , \tag{S122}$$

$$k = f\left[\nu M_\perp - M_\parallel - \kappa f(\Delta\alpha M_0 M_\parallel + \nu M_\alpha)\right]\ . \tag{S123}$$

As is evident, poroelasticity provides effective inertia to the active dynamics, although the flow itself is highly overdamped in the Darcy regime.

While there are many parameters here, a simple illustrative limit is to consider the case when $\kappa = 0$ and $\lambda_\parallel M_\perp > \lambda_\perp M_\parallel$ (only accessible when $\lambda_\perp \ll \lambda_\parallel$). In this case, $\alpha_\perp \ll \alpha_\parallel$ as well. Noting that $\nu \propto 1/a^4 \gg 1$, we have both $m > 0$ and $k > 0$, while $\xi$ can switch sign from positive to negative if activity is sufficiently strong ($\alpha_\parallel > \alpha_c$). For $a \ll 1$ and $\mathrm{Da}_\perp \gg 1$, the threshold activity for the instability scales as

$$\alpha_c \sim \frac{\nu}{M_0}\mathrm{Da}_\perp \propto \frac{\mathrm{Da}_\perp}{a^2}\ , \tag{S124}$$

while the oscillation frequency at onset scales as

$$\omega_c = \sqrt{\frac{k}{m}} \propto \frac{1}{\sqrt{\mathrm{Da}_\perp}}\ , \tag{S125}$$

recovering the same scaling of active hydraulic oscillations as before, though now in the absense of an underlying kinetic instability.

#### IV. LINEAR RESPONSE AND NONRECIPROCAL ELASTICITY

Here we analyze the emergent linear response behavior of muscle to an applied oscillatory strain $\boldsymbol{\epsilon}(\omega)$. Starting with the linearized kinetic equations (Eq. S44, S45) and Fourier transforming in time $[\psi(\omega) = \int \mathrm{d}t\, e^{-i\omega t}\psi(t)]$, we can write the linearized active stress density $\delta\sigma_a = \delta y + \delta n$ in terms of the strains as follows

$$\delta\sigma_a(\omega) = \chi_{\|}(\omega)\delta\epsilon_{zz}(\omega) + \chi_{\perp}(\omega)\delta\epsilon_{\perp}(\omega)\ , \tag{S126}$$

with the frequency dependent longitudinal and transverse susceptibilities given by

$$\chi_{\|}(\omega) = \frac{i\omega\lambda_{\|}}{(i\omega + f)}\left[1 - \frac{\beta f}{(i\omega+1)}\right]\ , \tag{S127}$$

$$\chi_{\perp}(\omega) = -\frac{\kappa f}{(i\omega+1)} + \frac{i\omega\lambda_{\perp}}{(i\omega+f)}\left[1 - \frac{\beta f}{(i\omega+1)}\right]\ . \tag{S128}$$

As is evident, the active stress response is naturally viscoelastic and it incorporates both a strain and strain rate dependence through biophysical feedback, as empirically suggested previously [S59]. At low frequencies and when $\beta f > 1$, $\chi_{\|} \propto -i\omega(\beta f - 1)$ providing a negative (hence active) viscous response of the active stress to strain due to the transient crosslinking of motors. In contrast, $\chi_{\perp}$ has both elastic and viscous elements at low frequencies due to both the binding kinetics and geometry. Note, if $\beta f > 1$, then the low frequency viscous response is negative, hence *active* and work producing. At high frequencies ($i\omega \gg 1, f$), both active susceptibilities asymptote to constants determined by the geometry of the sarcomeric structure ($\chi_{\|,\perp} \to \lambda_{\|,\perp}$), signalling an enhanced active stiffness at high frequencies arising from heavily crosslinked motors not having sufficient time to unbind (on time scales of $\omega^{-1}$).

By analyzing Eqs. S127, S128, we see that there are two poles, one at $-i\omega = 1$ ($\tau_k^{-1}$ in dimensionful units) and the other at $-i\omega = f$ ($\omega_{\mathrm{off}}(y_0)$ in dimensionful units). Under conditions of ATP depletion leading to *rigor*, $\omega_{\mathrm{off}} \to 0$ and all the myosin motors remain bound and stalled ($n_m^0 \to 1$ and $f \to 0$), leading to $\chi_{\|,\perp} = \lambda_{\|,\perp} > 0$, which provides an effective positive elastic element signalling an active *stiffening* of muscle by the crosslinked motors. Altogether we have three bare time scales present in the problem, two microscopic and one mesoscopic, with the general heirarchy $\tau_k < \omega_{\mathrm{off}}^{-1} \ll \tau_p$, or in nondimensional form $\mathrm{Da}^{-1} \ll f < 1$.

The total stress $\boldsymbol{\sigma} = \phi(\boldsymbol{\sigma}^{el} + \boldsymbol{\sigma}^{a}) - (1-\phi)p\mathbf{1}$ can be decomposed into isotropic and shear components $\sigma_{o,s} = \sigma_{zz} \pm \sigma_{\perp}/2$ along with the corresponding isotropic and shear strains $\epsilon_{o,s} = (\epsilon_{zz} \pm \epsilon_{\perp})/2$. Other shear components involving stresses in the azimuthal (hoop) direction exist, but they only play a minor role and can be included in a trivial extension. The linearized response is then encoded in the effective constitutive relation

$$\begin{pmatrix}\sigma_o\\ \sigma_s\end{pmatrix} = \phi\begin{bmatrix} Y_z + B_{\perp}(1+2\nu_z)^2 + 4(B_{\perp}+G)\alpha\chi & Y_z - B_{\perp}(1-4\nu_z^2) + 2(B_{\perp}+G)\alpha\Delta\chi \\ Y_z - B_{\perp}(1-4\nu_z^2) + 2(B_{\perp}+G)\Delta\alpha\chi & Y_z + B_{\perp}(1-2\nu_z)^2 + (B_{\perp}+G)\Delta\alpha\Delta\chi\end{bmatrix}\begin{pmatrix}\epsilon_o\\ \epsilon_s\end{pmatrix} - (1-\phi)p\begin{pmatrix}2\\ 0\end{pmatrix}\ , \tag{S129}$$

where $\alpha = (\alpha_{\|} + \alpha_{\perp})/2$ and $\Delta\alpha = \alpha_{\|} - \alpha_{\perp}$. Similarly, $\chi = (\chi_{\|} + \chi_{\perp})/2$ and $\Delta\chi = \chi_{\|} - \chi_{\perp}$. Here, we have retained the general form of the uniaxial elastic moduli (Eq. S2) and written the stress and pressue in dimensionful form. For $\nu_z = 1/2$ and considering a single spatial Fourier mode, this equation simplifies to (Fig. 3C, main text)

$$\begin{pmatrix}\sigma_o\\ \sigma_s\end{pmatrix} = \begin{bmatrix} B_{\mathrm{eff}} & C - \zeta\\ C + \zeta & Y_{\mathrm{eff}}\end{bmatrix}\begin{pmatrix}\epsilon_o\\ \epsilon_s\end{pmatrix}\ . \tag{S130}$$

Here we have used Darcy's law and incompressibility to replace the pressure in favour of the volumetric strain rate, i.e., $-[(a^2/K_r)q_z^2 + q_{\perp}^2]p = i\omega\ (B+G)[\phi\mathrm{Da}_{\perp}/(1-\phi)]\epsilon_o$. This leads to effective frequency dependent complex moduli given by

$$B_{\mathrm{eff}}(\omega) = \phi\left\{Y + 4B + (B+G)\left[\alpha\chi(\omega) + \frac{2i\omega\ \mathrm{Da}_{\perp}}{(a^2/K_r)q_z^2 + q_{\perp}^2}\right]\right\}\ , \tag{S131}$$

$$Y_{\mathrm{eff}}(\omega) = \phi\left[Y + (B+G)\Delta\alpha\Delta\chi(\omega)\right]\ , \tag{S132}$$

$$C(\omega) = \phi\left\{Y + (B+G)\left[\alpha\Delta\chi(\omega) + \Delta\alpha\chi(\omega)\right]\right\}\ , \tag{S133}$$

$$\zeta(\omega) = \phi(B+G)\left[\Delta\alpha\chi(\omega) - \alpha\Delta\chi(\omega)\right]\ . \tag{S134}$$

As noted in the main text, $B_{\mathrm{eff}}$ and $Y_{\mathrm{eff}}$ are effective bulk and Young's moduli, while $C$ is an effective anisotropic modulus present even in passive unixial solids and $\zeta$ is an active nonreciprocal modulus.

In a situation where deformations are primarily one-dimensional, the force response is governed by the effective (complex) Young's modulus, whose real ('elastic/storage' modulus) and imaginary ('viscous/loss' modulus) components are given by (in dimensionful form)

$$\mathrm{Re}[Y_{\mathrm{eff}}(\omega)] = Y_0 \left\{1 + A\,\mathrm{Re}[\Delta\chi(\omega)]\right\} \;, \quad \mathrm{Im}[Y_{\mathrm{eff}}(\omega)] = Y_0 A\,\mathrm{Im}[\Delta\chi(\omega)] \;, \tag{S135a}$$

$$\mathrm{Re}[\Delta\chi(\omega)] = \frac{\kappa f}{1+\omega^2\tau_k^2} + \frac{\Delta\lambda\omega^2\tau_k^2}{\omega^2\tau_k^2+f^2}\left[1 - \frac{\beta f(1+f)}{1+\omega^2\tau_k^2}\right] \;, \tag{S135b}$$

$$\mathrm{Im}[\Delta\chi(\omega)] = \frac{\omega f\tau_k}{(1+\omega^2\tau_k^2)(\omega^2\tau_k^2+f^2)}\left\{[\Delta\lambda(1+\beta)-\kappa]\,\omega^2\tau_k^2 - \left[\kappa f^2 + \Delta\lambda(\beta f - 1)\right]\right\} \;, \tag{S135c}$$

where $Y_0 = \phi Y$ and $A = \Delta\alpha(B+G)/Y$. This functional form is used to fit the rheology experiments in Fig. 4 (main text), see also Sec. V C. Some simple limits are instructive. At low frequencies ($\omega\tau_k \ll 1$),

$$\mathrm{Re}[Y_{\mathrm{eff}}(\omega)] = Y_0\left[1 + A\kappa f\right] + \mathcal{O}(\omega^2\tau_k^2) \;, \tag{S136}$$

$$\mathrm{Im}[Y_{\mathrm{eff}}(\omega)] = -Y_0 A\left[\kappa f + \frac{\Delta\lambda}{f}(\beta f - 1)\right]\omega\tau_k + \mathcal{O}(\omega^3\tau_k^3) \;. \tag{S137}$$

Notably, the low frequency stiffness acquires a positive active contribution ($\sim A\kappa f$), while the viscous response is *negative* (i.e., active) for strong enough feedback ($\beta f > 1$). At high frequencies ($\omega\tau_k \gg 1$)

$$\mathrm{Re}[Y_{\mathrm{eff}}(\omega)] = Y_0\left[1 + A\Delta\lambda\right] + \mathcal{O}\left(\frac{1}{\omega^2\tau_k^2}\right) \;, \tag{S138}$$

$$\mathrm{Im}[Y_{\mathrm{eff}}(\omega)] = +\frac{Y_0 A f}{\omega\tau_k}\left[\Delta\lambda(1+\beta) - \kappa\right] + \mathcal{O}\left(\frac{1}{\omega^3\tau_k^3}\right) \;. \tag{S139}$$

The high frequency stiffness is also actively enhanced (more than the low frequency contribution if $\Delta\lambda > \kappa f$), while the viscous response is positive (i.e., dissipative) and decaying for sufficiently strong feedback ($\Delta\lambda(1+\beta) > \kappa$). From Eq. S135 we see that the viscous modulus vanishes at an intermediary frequency $\omega_*$, i.e., $\mathrm{Im}[Y_{\mathrm{eff}}(\omega_*)] = 0$, which is given by

$$\omega_* = \sqrt{\frac{\kappa f^2 + \Delta\lambda(\beta f - 1)}{\Delta\lambda(1+\beta) - \kappa}} \;, \tag{S140}$$

which remarkably is the same as the onset oscillation frequency for a purely kinetic instability (see Eqs. S81, S97)! Hence, in a regime where spontaneous oscillations are driven by a kinetic instability (without hydraulics), little to no power is generated by the 1D viscous response as $\mathrm{Im}[Y_{\mathrm{eff}}(\omega_*)] = 0$. As a result, oscillatory work production requires either hydraulics (to shift the operating frequency away from $\omega = \omega_*$) or 3D deformations (allowing effects including odd elasticity) to be necessarily present.

The odd elastic ($\mathrm{Re}[\zeta(\omega)]$) and odd viscous ($\mathrm{Im}[\zeta(\omega)]$) moduli are given by (in dimensionful form)

$$\mathrm{Re}[\zeta(\omega)] = \phi(B+G)\left\{-\frac{\alpha_\parallel\kappa f}{1+\omega^2\tau_k^2} + \frac{(\alpha_\parallel\lambda_\perp - \alpha_\perp\lambda_\parallel)\omega^2\tau_k^2}{(\omega^2\tau_k^2+f^2)}\left[1 - \frac{\beta f(1+f)}{(1+\omega^2\tau_k^2)}\right]\right\} \;, \tag{S141a}$$

$$\mathrm{Im}[\zeta(\omega)] = \phi(B+G)\omega\tau_k f\left\{\frac{\alpha_\parallel\kappa}{1+\omega^2\tau_k^2} + \left(\frac{\alpha_\parallel\lambda_\perp - \alpha_\perp\lambda_\parallel}{\omega^2\tau_k^2+f^2}\right)\left[1 + \frac{\beta(\omega^2\tau_k^2 - f^2)}{1+\omega^2\tau_k^2}\right]\right\} \;. \tag{S141b}$$

Once again, simple limits are instructive. At low frequencies ($\omega\tau_k \ll 1$)

$$\mathrm{Re}[\zeta(\omega)] = -\phi(B+G)\alpha_\parallel\kappa f + \mathcal{O}(\omega^2\tau_k^2) \;, \tag{S142}$$

$$\mathrm{Im}[\zeta(\omega)] = \phi(B+G)\omega\tau_k f\left[\alpha_\parallel\kappa - (\alpha_\parallel\lambda_\perp - \alpha_\perp\lambda_\parallel)\frac{(\beta f - 1)}{f^2}\right] + \mathcal{O}(\omega^3\tau_k^3) \;. \tag{S143}$$

The odd elastic modulus is nonzero and negative at low frequency, while the odd viscous modulus can be of either sign depending on the feedback parameters. At high frequencies ($\omega\tau_k \gg 1$), we have

$$\mathrm{Re}[\zeta(\omega)] = \phi(B+G)(\alpha_\parallel\lambda_\perp - \alpha_\perp\lambda_\parallel) + \mathcal{O}\left(\frac{1}{\omega^2\tau_k^2}\right) \;, \tag{S144}$$

$$\mathrm{Im}[\zeta(\omega)] = \phi(B+G)\frac{f}{\omega\tau_k}\left[\alpha_\parallel\kappa + (\alpha_\parallel\lambda_\perp - \alpha_\perp\lambda_\parallel)(1+\beta)\right] + \mathcal{O}\left(\frac{1}{\omega^3\tau_k^3}\right) \;. \tag{S145}$$

The high-frequency limit of the odd elastic modulus vanishes if $\alpha_\perp/\alpha_\parallel = \lambda_\perp/\lambda_\parallel$ as we have in our model when we assume a simple homogeneous crossbridge binding geometry. In this case the odd (complex) modulus simplifies to be (i.e., with $\alpha_\parallel\lambda_\perp = \alpha_\perp\lambda_\parallel$)

$$\zeta(\omega) = -\phi(B+G)\frac{\alpha_\parallel \kappa f}{1+i\omega\tau_k} \ . \tag{S146}$$

In general though, $\zeta(\omega)$ could display the more complex frequency dependence due to $\alpha_\parallel\lambda_\perp \neq \alpha_\perp\lambda_\parallel$, though this only affects the high frequency response, while the low frequency odd modulus remains insensitive to these details, being primarily controlled by the axial activity $\alpha_\parallel$ and the geometric feedback parameter $\kappa$.

In order to understand why $\zeta$ is an active modulus and related to nonreciprocity, it is useful to consider a general linear stress-strain relation

$$\sigma_{ij} = \sigma^0_{ij} + \mathcal{A}_{ijk\ell}\delta\epsilon_{k\ell} \ , \tag{S147}$$

where $\sigma^0_{ij}$ is a steady state pre-stress and $\delta\epsilon_{k\ell}$ is the deviation in the strain from its steady state value. The modulus tensor $\mathcal{A}_{ijk\ell}$ enodes the spatial and temporal symmetries of the problem. In particular, if the material obeys energy conservation (in which case Eq. S147 derives from a potential energy function), then we have the following symmetry [S60, S61]

$$\mathcal{A}_{ijk\ell} - \mathcal{A}_{k\ell ij} = \sigma^0_{ij}\delta_{k\ell} - \sigma^0_{k\ell}\delta_{ij} \ , \tag{S148}$$

which enforces Maxwell-Betti reciprocity in passive media [S62]. Any violation of Eq. S148 necessitates a nonequilibrium drive that maintains an energy flux. As we consider the simple case of analyzing the response of muscle with vanishing prestress, $\sigma^0_{ij} = 0$ (but nonzero prestrain), mechanical nonreciprocity is then equivalent to a nonsymmetric tensor of elastic moduli (which in our model is quantified by $\zeta$). Other states of muscle with prestress are possible and can be addressed by a straightforward extension of our analysis.

The work produced in a periodic cycle is given by $W = -\oint \sigma_{ij}\mathrm{d}\epsilon_{ij}$ ($W > 0$ when work is produced and $W < 0$ when work is dissipated). Assuming $\sigma_{rr} = \sigma_{\varphi\varphi}$, then we can simply write the work as

$$W = -\oint \left[\sigma_s \mathrm{d}\epsilon_s + \sigma_o \mathrm{d}\epsilon_o\right] \ . \tag{S149}$$

The total work produced in periodic deformations with a cycling frequency $\omega$ can then be explicitly written as

$$W = W_{\text{even}} + W_{\text{odd}} \ , \tag{S150}$$

$$W_{\text{even}} = -\frac{\text{Im}[B_{\text{eff}}]}{\omega}\oint \mathrm{d}t\ (\dot\epsilon_o)^2 - \frac{\text{Im}[Y_{\text{eff}}]}{\omega}\oint \mathrm{d}t\ (\dot\epsilon_s)^2 - 2\frac{\text{Im}[C]}{\omega}\oint \mathrm{d}t\ \dot\epsilon_o\dot\epsilon_s \ , \tag{S151}$$

$$W_{\text{odd}} = -\text{Re}[\zeta]\oint \left[\epsilon_n \mathrm{d}\epsilon_s - \epsilon_s \mathrm{d}\epsilon_n\right] \ . \tag{S152}$$

The reciprocal elastic couplings (Re$[B_{\text{eff}}]$, Re$[Y_{\text{eff}}]$ and Re$[C]$) do not produce or dissipate any work. When the effective viscous response is negative, i.e., Im$[B_{\text{eff}}] < 0$, Im$[Y_{\text{eff}}] < 0$ or Im$[C] < 0$, then work is actively produced ($W_{\text{even}} > 0$). This mode of power generation relies on strain *rates* (rater than strains) and is present even in 1D deformations. When allowing for full 3D spatial deformations, a different mode of energy production is available through odd-cycles. The work done in the odd-elastic sector is given by

$$W_{\text{odd}} = -\text{Re}[\zeta]\oint \left[\epsilon_n \mathrm{d}\epsilon_s - \epsilon_s \mathrm{d}\epsilon_n\right] = \frac{\text{Re}[\zeta]}{2}\oint \left[\epsilon_{zz}\mathrm{d}\epsilon_\perp - \epsilon_\perp \mathrm{d}\epsilon_{zz}\right] \ , \tag{S153}$$

i.e., $W_{\text{odd}} = \text{Re}[\zeta]\times$(Area enclosed by $\epsilon_\perp - \epsilon_{zz}$ strain loop). Note that odd viscosity Im$[\zeta]$ does not generate or dissipate any work [S63].

#### V. DATA COMPILATION AND ANALYSIS

##### A. 3D strains in spontaneous sarcomeric oscillations

The plots in Fig. 2 use time series data of 3D sarcomeric deformations measured in spontaneously oscillating muscle fibers, that we compiled from the following references. The data were extracted using WebPlotDigitizer [S122].

| Organism | Toadfish (*Opsanus*) | Rattlesnake (*Crotalus*) | Fruit fly (*Drosophila*) | Cicada (*Cicadoidea*) | Mosquito (*Culicidae*) |
|---|---|---|---|---|---|
| Muscle[a] | Swimbladder sonic muscle (S) | Rattle tail shaker muscle (S) | Flight muscle (AS) | Tymbal muscle (AS/S) | Flight muscle (AS) |
| $\omega$ (Hz) | $100-220$ [S64–S67] | $20-90$ [S64, S68] | $156-200$ [S54, S69] | $40-550$ [S70–S72] | $300-800$ [S73, S74] |
| $\tau_k^{-1}$ (1/s) | 121 [S75] | $100-125$ [S64, S76] | $(3.5-3.9)\times 10^3$ [S56] | $66-125$ [S70, S77] | $(3.5-3.9)\times 10^3$ |
| $E$ (MPa) | $7.6-9.4$ [S75] | 8 | $0.16-0.26$ [S78] | 4 [S79] | 0.16-0.26 |
| $\eta$ (Pa s)[b] | 0.001 | 0.001 | 0.001 | 0.001 | 0.001 |
| $\ell_p$ (nm)[c] | 26.2 [S26, S80] | 20-30 [S81] | 55 [S78] | 33-50 [S70, S82] | 45 [S83] |
| $R$ ($\mu$m) | 5.8-14.3 [S84] | 23.5 [S85] | 50-100 [S86] | 26-32 [S77, S87] | 36-42.5 [S88] |
| $L$ (mm) | 10-13 [S89] | 20 [S85] | 1 [S69, S86] | 7.5 [S90] | $\lesssim 2$ [S83] |
| $\ell_s$ ($\mu$m) | 2.2-3.2 [S65] | 2.4 [S81] | 3.4 [S69] | 2.2-2.8 [S70, S77] | 1.5-2.9 [S91, S92] |
| Comments | Includes both juveniles and adults, with males producing the high-frquency "boatwhistle" | $\tau_k$ estimated from twitch time; $E$ and $\ell_p$ estimated similar to sonic and skeletal muscles | | Cicada flight muscle is synchronous, but the tymbal muscle can be either across genera, e.g., *Platypleura* has asynchronous sonic muscle [S70, S82]; $E$ is similar to *Lethocerus* flight muscle [S79] and $\tau_k$ estimated from twitch time | $E, \tau_k$ estimated similar to *Drosophila* (*Diptera*) [S93]. |

| Organism | Giant waterbug (*Lethocerus*) | Bumble bee (*Bombus*) | Rabbit (*Oryctolagus*) | Hummingbird (*Trochilidae*) | Mouse (*Mus*) |
|---|---|---|---|---|---|
| Muscle[a] | Flight muscle (AS) | Flight muscle (AS) | Psoas skeletal muscle (S) | Pectoralis flight muscle (S) | Cardiac muscle ($\sim$AS) |
| $\omega$ (Hz) | 38-44 [S94] | 120-150 [S95] | 0.6-1 [S96, S97] | 15-80 [S98] | 8.3-13 [S99, S100] |
| $\tau_k^{-1}$ (1/s) | 153-177 [S101] | 664-719 [S102, S103] | 352-526 [S104] | 125 [S105] | 80-100 [S106] |
| $E$ (MPa) | 4 [S107, S108] | 0.73 [S109] | 8-10 [S110] | 0.1-1 [S111] | 0.5 [S106] |
| $\eta$ (Pa s)[b] | 0.001 | 0.001 | 0.001 | 0.001 | 0.001 |
| $\ell_p$ (nm)[c] | 45.9 [S26] | 45 [S95] | 39.4-43.3 [S26] | 45-50 | 35-38 [S112] |
| $R$ ($\mu$m) | 27 [S94] | 33 [S103] | 37.5-40 [S113, S114] | 8-15 [S111, S115] | 5-10 [S100, S116] |
| $L$ (mm) | 3-5 [S101, S117] | 3-6 [S109, S118] | 10-12 [S119] | $\sim 5$ [S115] | $\sim 0.1$ [S100, S116] |
| $\ell_s$ ($\mu$m) | 2.7-2.8 [S94] | 2.2 [S118] | 2.2-3.7 [S26] | 1.7-2.3 [S111, S115] | 1.75-2.2 [S100] |
| Comments | | | $L$ estimated from other hindlimb muscles | $E$ and $\ell_p$ estimated similar to asynchronous flight muscle, $E$ consistent with reported low force generation [S111], assuming strain $\sim 1\%$ | Cardiac muscle shares stretch activation and mechanical characteristics with insect flight muscle [S120, S121]. |

[a] AS: asynchronous muscle, S: synchronous muscle; [b] viscosity is assumed to be that of water; [c] pore size estimated as inter-thick filament spacing $d_{10}$.

TABLE I. Estimates of kinetic, structural, material and geometric parameters of fast muscles in different organisms. The notation is $\omega$: natural oscillation frequency, $\tau_k$: kinetic rate, $E$: effective elastic modulus, $\eta$: viscosity, $\ell_p$: hydraulic pore size, $R$: resting fiber radius, $L$: resting fiber length, $\ell_s$: resting sarcomere length.

Figs. 2A, B reproduce data from Fig. 2A in Ref. [S97] and Fig. 3C in Ref. [S96] respectively. Both measurements involve high-resolution phase contrast optical imaging of glycerinated myofibrils (*in-vitro*) from rabbit psoas muscle at 25°C, with different resting sarcomere lengths ($SL_0$) and concentration of ATP (A: $SL_0 = 2.6$ $\mu$m, 0.2 mM MgATP; B: $SL_0 = 2.3$ $\mu$m, 2.2 mM ATP). The fibers oscillate spontaneously and the sarcomere length (SL) and width of the A-band (AW) are reported as a function of time. The longitudinal ($\epsilon_{zz}$) and transverse ($\epsilon_\perp$) strains were computed simply as,

$$\epsilon_{zz}(t) = \frac{\mathrm{SL}(t) - \overline{\mathrm{SL}}}{\overline{\mathrm{SL}}} , \quad \epsilon_{\perp}(t) = 2\left(\frac{\mathrm{AW}(t) - \overline{\mathrm{AW}}}{\overline{\mathrm{AW}}}\right) , \tag{S154}$$

where $\overline{f} = (1/T)\sum_{t=0}^{T} f(t)$ is the time-average of $f(t)$. We normalize the strains using the time-averged SL and AW to center the strain curves about zero (the time averaged length matches the resting length within error, when reported). The factor of 2 in the definition of the transverse strain is because $\epsilon_{\perp} = \epsilon_{rr} + \epsilon_{\varphi\varphi}$ includes both radial ($\epsilon_{rr}$) and hoop ($\epsilon_{\varphi\varphi}$) strains that become equal to each other ($\epsilon_{\varphi\varphi} = \epsilon_{rr}$) upon assuming the deformation is both axisymmetric and radially affine (i.e., $\partial_r \epsilon_{rr} = 0$). We bin the data with $\Delta t = 0.05$ s and slice the time-series into estimated cycles of equal length that we average over. The optimal size of the cycle is chosen by minimizing the unbiased sample variance obtained from averaging over multiple cycles in the time-series. This provides the best estimate of the oscillation time-period (A: $T = 1.69 \pm 0.05$ s, B: $T = 0.99 \pm 0.05$ s) consistent with the reported values in Refs. [S96, S97]. The corresponding mean strains are plotted as a function of time and as a phase portrait with the standard deviation shown as colored regions or error bars. To compute the signed area (Area $= (1/2)\oint[\epsilon_{zz}\mathrm{d}\epsilon_{\perp} - \epsilon_{\perp}\mathrm{d}\epsilon_{zz}]$) enclosed by the closed curve in strain space, we generate a synthetic ensemble of $\epsilon_{zz}$ and $\epsilon_{\perp}$ strains (sample size $N = 500$) at each time point that are each Gaussian distributed with the previously computed mean and standard deviation. We then evaluate the area integral Area $= (1/2)\oint[\epsilon_{zz}\mathrm{d}\epsilon_{\perp} - \epsilon_{\perp}\mathrm{d}\epsilon_{zz}]$ using the trapezoidal rule for each sample curve of strains and average the computed Area over the $N = 500$ samples. The resulting average and sample standard deviation for the enclosed Area are quoted in the plots.

Fig. 2C involves *in-vivo* measurements of strain in asynchronous flight muscle (DLM: dorsal longitudinal) of a living fruit fly in tethered flight [S54, S69, S123]. The data in this figure are compiled from multiple sources. The longitudinal strains ($\epsilon_{zz}$) plotted are obtained directly from high-speed optical measurements of muscle length in *Drosophila virilis* from Fig. 2A in Ref. [S69]. The time series is binned as before, now with $\Delta t = 0.5$ ms and the optimal time period is obtained by minimizing the overlap error across successive cycles to give $T = 6.4 \pm 0.5$ ms, consistent with the reported wingbeat frequency $\omega \approx 154$ Hz. These axial deformations estimated from muscle length changes are also consistent with independent measurements of filament strains estimated from small angle X-ray scattering in *Drosophila melanogaster* and *Drosophila metleri* [S123]. The transverse strain is negligible at all time, i.e., $\epsilon_{\perp} \sim 0$ (hence, no strain cycle), and was obtained from time-resolved X-ray diffraction measurements of the thick filament lattice spacing ($d_{10}$) in flying *Drosophila melanogaster*, with subnanometer precision from Ref. [S54]. Given the reported constancy of the lattice spacing, we estimate $\epsilon_{\perp} = (8 \pm 0.4) \times 10^{-3}$ using the maximal difference of $d_{10}$ between the up and down stroke [S54], and consider it to be constant in time. Although the strain measurements are obtained from different insects, we expect all *Drosophilids* to have similar in-vivo dynamics of muscle during flight. The Area enclosed by the strain curve (both its mean and standard deviation) is once again estimated as before using an ensemble of $N = 500$ Gaussian distributed sample points.

Fig. 2D involves *ex-vivo* measurements of strains in intact synchronous flight muscle (DLM) from *Manduca sexta* (hawkmoth) from Fig. 4A in Ref. [S124]. An oscillatory stretch (4% axial strain) is imposed on the isolated, whole muscle at a physiologically relevant frequency ($\omega = 25$ Hz), along with an electrical stimulation with an activation phase of 0.5 (matching *in-vivo* flight conditions). The transverse strain $\epsilon_{\perp}$ was estimated as in Eq. S154 from the $d_{10}$ lattice spacing displacements measured using time-resolved X-ray diffraction, upon assuming $\overline{d}_{10} = 49$ nm along with the 95% confidence interval for the displacement as reported in Ref. [S124]. The axial strain $\epsilon_{zz} = -0.04\cos(2\pi t/T)$ with $T = 40$ ms is exactly known as it is directly imposed on the muscle. The mean and standard deviation of the area enclosed by the strain cycle are computed again as before using a synthetic ensemble of $N = 500$ gaussian distributed strain samples. Note that the strains measured *ex-vivo* are consistent with the more variable *in-vivo* X-ray measurements of filament lattice deformations in flying hawkmoths [S125]. Hence, these results are expected to be physiologically relevant as well.

##### B. Estimating kinetic, poroelastic and contraction time scales in muscle

We compile structural, kinetic, material and geometric estimates of muscle fibers from different types of fast muscles and organisms. The data and associated references are listed in Table I. We estimate the relevant poroelastic time scales ($\tau_p^{\parallel,\perp}$) and corresponding Damköhler numbers ($\mathrm{Da}_{\parallel,\perp}$) in both the axial and radial directions (see Eqs. S41, S42). We neglect anisotropic effects in the elastic stiffness $E$ and assume the hydraulic permeabilities are constant with average values $K_{\parallel} \approx 0.29$ and $K_{\perp} \approx 0.037$ estimated using an average $\ell_p \sim 30$ nm, $r_0 \sim 8$ nm and $\phi \sim 0.15$ [$\phi \sim 0.22 - 0.097$ based on a hexagonal packing of thick and thin filaments with radii $r_{\mathrm{thick}} = 8$ nm, $r_{\mathrm{thin}} = 4$ nm and inter-thick filament spacing $d_{10} \sim 25 - 55$ nm [S26, S39, S78]]. As a result, $K_r = K_{\perp}/K_{\parallel} \approx 0.1$.

In the main text, we plot the experimental data as a function of $\mathrm{Da}_{\perp}$ as radial flow is expected to be the relevant (fastest) hydraulic process in a muscle fiber. In Fig. S2, we additionally plot the same data against $\mathrm{Da}_{\parallel}$ and $a^2\mathrm{Da}_{\parallel}$

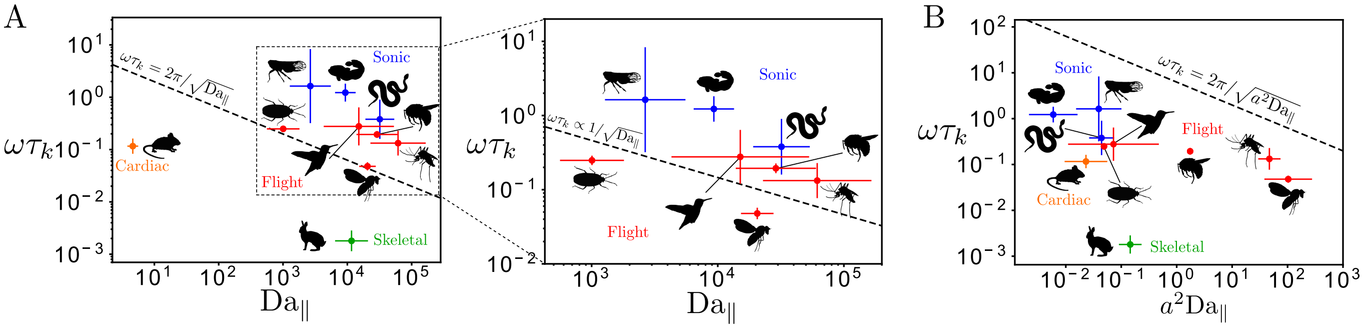


FIG. S2. Nondimensionalized characteristic oscillation frequencies ($\omega\tau_k$) in fast sonic (blue), flight (red), cardiac (orange) and skeletal (green) muscles across the animal kingdom plotted against the axial Damköhler number ($\mathrm{Da}_{\|}$, A with zoomed in view of the marked square shown enclosing all flight and sonic muscle examples) and the same multiplied by the squared aspect ratio of the fiber ($a^2\mathrm{Da}_{\|}$, B). The dashed lines show the expected scaling from active hydraulic oscillations (with numerical prefactor $2\pi$ displayed when fixed, else only scaling relation shown) assuming the corresponding poroelastic timescale is rate-limiting. The data plotted against $\mathrm{Da}_{\|}$ (A) displays a trend (for sonic and flight muscles, see zoomed inset) consistent with the hydraulic scaling $\omega\tau_k \propto 1/\sqrt{\mathrm{Da}_{\|}}$, while the plot against $a^2\mathrm{Da}_{\perp}$ (B) is simply a rescaled version of Fig. 3B (main text). The data are represented using crosses that denote the range (minimum to maximum) of the respective parameters and the central point is the geometric mean.

| | Parameter | *Drosophila* flight muscle | Mouse cardiac muscle | Rabbit skeletal muscle |
|---|---|---|---|---|
| Estimated | $Y_0 = \phi Y$ $^\dagger$ | $0.14 - 0.22$ | $0.034$ | $0.1 - 0.14$ |
| | $\Delta\lambda$ | $0.5 - 1$ | $0.5 - 1$ | $0.5 - 1$ |
| Fit | $\Omega_0 = \tau_k^{-1}$ $^*$ | $514 \pm 75$ | $95 \pm 1.5$ | $88 \pm 1.4$ |
| | $A = \Delta\alpha(B+G)/Y$ | $0.9 \pm 0.5$ | $100 \pm 25$ | $153 \pm 43$ |
| | $f$ | $0.5 \pm 0.1$ | $0.12 \pm 0.01$ | $0.26 \pm 0.01$ |
| | $\beta$ | $7 \pm 2$ | $9.2 \pm 0.7$ | $4.6 \pm 0.2$ |
| | $\kappa$ | $1.4 \pm 0.5$ | $1.1 \pm 0.3$ | $2 \pm 0.5$ |

$^*$ Hz; $^\dagger$ MN/m$^2$

TABLE II. Model parameters estimated ($Y_0, \Delta\lambda$) from literature (see text) and obtained ($\Omega_0, A, f, \kappa, \beta$) from a nonlinear fit of 1D oscillatory rheology measurements in different muscle fibers. The optimal fit parameter values are quoted as mean $\pm$ standard deviation, the latter including the variance from both the range of $Y_0, \Delta\lambda$ values and the estimated covariance from each nonlinear least squares fit.

($a = R/L$, the fiber aspect ratio). Although axial flow is not expected to be the dominant (rate determining) hydraulic process (so the calculated oscillation frequency is not a reliable upper bound), in Fig. S2 (left) we nonetheless find the contraction frequency for some of the fastest muscles (sonic and flight) across the animal kingdom to scale as $\omega\tau_k \sim 1/\sqrt{\mathrm{Da}_{\|}}$. Such a correlated scaling trend is perhaps expected on general allometric grounds ($R \propto L$). The plot against $a^2\mathrm{Da}_{\|}$ (Fig. S2, right) on the other hand is similar to the one in the main text (to be expected as $\mathrm{Da}_{\perp} = a^2\mathrm{Da}_{\|}/K_r$). In all the plots, the crosses denote the range (minimum to maximum) of the respective Damköhler numbers and nondimensionalized frequencies, with the central point denoting the geometric mean.

##### C. Oscillatory muscle rheology experiments

Data on sinusoidal uniaxial rheology of permeabilized muscle fibers were compiled from Fig.2A-D in Ref. [S78] (Drosophila flight muscle), Fig. 6A-B in Ref. [S106] (Mouse cardiac muscle) and Fig.3A-B (activation) in Ref. [S110] (Rabbit skeletal muscle). We extract $\mathrm{Re}[Y_{\mathrm{eff}}(\omega)]$ and $\mathrm{Im}[Y_{\mathrm{eff}}]$ using the WebPlotDigitizer [S122]. When error bars are not provided for the measured data, we use the plotted marker size as an estimate of the error in data extraction. For rabbit skeletal muscle, the data from Ref. [S110] only reports the dynamic modulus (amplitude $|Y_{\mathrm{eff}}(\omega)|$) and the phase ($\arg[Y_{\mathrm{eff}}(\omega)]$) of the response, which we convert into the real and imaginary parts of the response by constructing a synthetic ensemble ($N = 500$) of the frequency dependent phase and amplitude of the modulus (gaussian distributed about the mean), linearly interpolating over $\omega$ and then computing $\mathrm{Re}[Y_{\mathrm{eff}}(\omega)]$ and $\mathrm{Im}[Y_{\mathrm{eff}}]$ (both the sample mean and standard deviation) with frequency steps of $\Delta\omega = 0.5$ Hz.

We then fit the experimental data in all cases to our linearized model assuming affine deformations. As the experiments are performed with glycerinated muscle fibers, fluid can be freely exchanged with the bath and hydraulics is no longer rate limiting. As a result, pressure gradients and fluid permeation are negligible and deformations are assumed isochoric ($\delta\epsilon_{zz} + \delta\epsilon_\perp = 0$). The uniaxial response to a small amplitude periodic stretch (assuming $\sigma_\perp = 0$ in a slender sarcomeric bundle, due to the free boundary) is then simply given by $Y_{\rm eff}(\omega) \equiv \delta\sigma_{zz}(\omega)/\delta\epsilon_{zz}(\omega) = \phi[Y + \Delta\alpha(B+G)\Delta\chi(\omega)]$ (Eq. S132). The same result can also be obtained systematically using a slender body description, as shown in Sec. III B. The real and imaginary parts of the response then correspond to the elastic (Re[$Y_{\rm eff}$], green) and the viscous (Im[$Y_{\rm eff}$], magenta) moduli, see Eq. S135. The low frequency response in the experiments sometimes includes a weak power law or logarithmic dependence on frequency, possibly arising from internal passive relaxation processes that occur on multiple time scales. We do not intend to capture these detailed features, but rather just the overall characteristic shape of the response curves across a wide variety of striated muscles.

The effective Young's modulus involves 7 independent parameters (5 dimensionless, 2 dimensionful): $\kappa$, $\beta$, $\Delta\lambda$ $f$, $A = \Delta\alpha(B+G)/Y$, $\tau_k$ and $Y_0 = \phi Y$. Note that the typical shape of the measured response curve requires atleast 7 parameters, all of which are traditionally fit for in a phenomenological fashion [S110]. Performing a nonlinear 7 parameter fit leads to large (orders of magnitude) variability in the parameter estimates as the model is sloppy [S126, S127]. In particular, a sensitivity analysis of a nonlinear least square fit reveals a typical exponential heirarchy of covariance matrix eigenvalues [S126], with a linear combination of variations in the parameters $Y_0$, $\kappa$, $A$ and $\Delta\lambda$ associated with the two sloppiest eigendirections. The most robust parameter is $f$, the fraction of unbound myosin, which is related to the motor duty ratio. As the four sloppiest parameters directly control the low frequency [Re[$Y_{\rm eff}(0)$] $= Y_0(1 + A\kappa f)$] and high frequency stiffness [Re[$Y_{\rm eff}(\infty)$] $= Y_0(1 + A\Delta\lambda)$], we can lift the degeneracy in parameter space by separately evaluating $Y_0$ and $\Delta\lambda$. Using typical estimates of the lattice and crossbridge geometry, we assume a thick-thin interfilament spacing $h_0 \sim 15 - 23$ nm, the power stroke size $y_0 \sim 8 - 10$ nm and the binding angle $\theta_0 \sim 45^\circ$ [S26, S37–S39, S128] to obtain $\Delta\lambda \approx 0.5 - 1$. Furthermore, we use independent measurements of the passive/relaxed stiffness ($Y_0$) of muscle fibers from previous experiments

- *Drosophila* flight muscle: $Y_0 = 0.14 - 0.22$ MN/m$^2$ [S129, S130]
- Mouse cardiac muscle: $Y_0 = 0.034$ MN/m$^2$ [S131]
- Rabbit psoas skeletal muscle: $Y_0 = 0.1 - 0.14$ MN/m$^2$ [S110, S132]

With these parameters fixed the model is no longer sloppy and we can robustly fit for the remaining 5 parameters $(\kappa, \beta, A, f, \tau_k)$.

The joint fit for both Re[$Y_{\rm eff}(\omega)$] and Im[$Y_{\rm eff}$] (Eq. S135) is performed using a custom Python [S133] code. A constrained nonlinear weighted least square minimization is performed using SciPy's [S134] in-built curve_fit function using the Trust Region Reflective algorithm that is a robust optimization algorithm for large sparse problems with bounds. The standard deviations extracted from the data are used as weights (absolute_sigma=True) and bounds are specified to ensure positivity of all parameters and $0 < f < 1$. For the data on mouse cardiac muscle, we perform the fit across the whole frequency range available, while for *Drosophila* flight muscle and rabbit skeletal muscle, we exclude points with $\omega < 4.5$ Hz as the curves include a low frequency (weak) power law response that is not captured by our model. Nonetheless, even if we include some or all of the low frequency data points, we obtain the same fits and parameter estimates (within error bars) as reported in Table II, but finding these optimal values becomes harder, with many runs not converging as more low frequency points are included. Hence to keep things tractable and simple, we exclude points $\omega < 4.5$ Hz for the flight and skeletal muscle data, while our checks ensure the robustness of our results to this cutoff.

To handle the range of *a priori* parameter estimates, we construct equispaced arrays of length 5 for both $Y_0$ and $\Delta\lambda$ spanning their prescribed ranges and for each pairwise combination of $(Y_0, \Delta\lambda)$ values (total 25 pairs), we perform a nonlinear fit and evaluate the model curves (Re[$Y_{\rm eff}$], Im[$Y_{\rm eff}$]; Eq. S135) given the optimal fit parameters. The fit model curves are then averaged over all values of $(Y_0, \Delta\lambda)$ within the specified range and plotted in Fig. 4 (main text) as solid lines (Re[$Y_{\rm eff}$]: green, Im[$Y_{\rm eff}$]: magenta). Standard deviations computed for these curves is smaller than the linewidth used in the plots. We have also checked that a different optimization routine using Mathematica's [S135] in-built NonlinearModelFit function with sequential minimization performed using Differential Evolution for global search and a QuasiNewton method for local minimization recovers the same results.

To evaluate the fit parameters and the odd elastic moduli, we generate, for each value of $(Y_0, \Delta\lambda)$, a synthetic ensemble ($N = 100$) of parameter values $(\Omega_0, A, f\beta, \kappa)$ normally distributed about their optimal values (obtained from curve fitting) with a standard deviation computed from the least-square covariance matrix. We ensure that all $N = 100$ randomly generated parameter values are positive with $0 < f < 1$ and discard any values that violate these bounds. With these synthetically generated parameter values, we compute the odd elastic modulus (Re[$\zeta$] $= \phi(B+G)$Re[$\alpha_\parallel\chi_\perp(\omega) - \alpha_\perp\chi_\parallel(\omega)$], Eq. S141) and evaluate the average odd modulus curve and its standard deviation

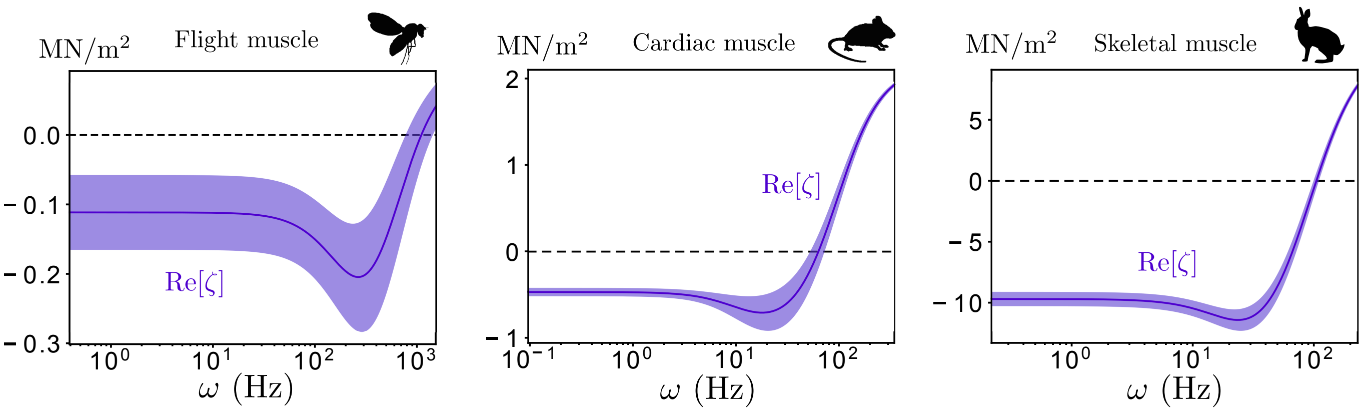


FIG. S3. Odd elastic moduli estimated for different muscle types - *Drosophila* flight muscle (left), mouse cardiac muscle (center), rabbit skeletal muscle (right) - using the parameters quoted in Table II along with the ratios $\lambda_\perp/\lambda_\parallel = 0.5$ and $\alpha_\perp/\alpha_\parallel = 0.1$. The average value is plotted as a blue line with the shaded region denoting one standard deviation. The dashed black line marks zero. Note, at moderate to high frequencies, the odd modulus develops a negative dip, eventually crossing through zero and becoming positive (when $\alpha_\parallel\lambda_\perp - \alpha_\perp\lambda_\parallel > 0$) for $\omega \to \infty$. The low frequency behaviour remains unchanged from Fig. 4 (main text) though.

(mean plotted as a blue curve with standard deviation as light blue shaded region in Fig. 4, main text) across all the random ensembles for all the values of $(Y_0, \Delta\lambda)$ ($N_{\rm tot} = 25N = 2500$). Similarly, the parameter estimates (both mean and standard deviation) reported in Table II are evaluated by averaging (and computing the variance of) the corresponding parameter values across all the synthetic random ensembles for all the values of $(Y_0, \Delta\lambda)$ ($N_{\rm tot} = 2500$).

In order to evaluate the odd elastic modulus, we additionally require the ratio of transverse to axial active forces ($\alpha_\perp/\alpha_\parallel$). Within our simple geometric model of crossbridge binding, $\alpha_\perp/\alpha_\parallel = \lambda_\perp/\lambda_\parallel = 0.5$ (assuming the crossbridge binding angle $\theta_0 \sim 45°$). As long as $\alpha_\perp/\alpha_\parallel = \lambda_\perp/\lambda_\parallel$, the qualitative functional form of the odd elastic modulus is independent of the precise value of the ratio (see Eq. S146) and is instead entirely controlled by the value of $\kappa$ (that we fit for). As seen in Fig. 4 (main text), the odd elastic modulus is always negative for all frequencies and vanishes as $\omega \to \infty$ (Eq. S146). More detailed models of crossbridge binding geometry do not need to obey the equality $\alpha_\perp/\alpha_\parallel = \lambda_\perp/\lambda_\parallel$. For instance, keeping $\lambda_\perp/\lambda_\parallel = 0.5$, we can set $\alpha_\perp/\alpha_\parallel = 0.1$, as suggested by more sophisticated geometric arguments of crossbridge binding [S37, S38]. Now the odd modulus acquires an additional contribution $\propto \alpha_\perp\lambda_\parallel - \alpha_\parallel\lambda_\perp$ (see Eq. S141) that only affects the high frequency behaviour (low frequency behaviour is unchanged). The resulting odd moduli for the three different muscle types (all other parameters the same as in Fig. 4, main text) are plotted in Fig. S3.

How do our parameter fits compare with our expectations from structural and molecular measurements? Molecular estimates for the feedback parameters $(\kappa, \beta)$ can be obtained using our microscopic description. From Eqs. S16, S17, we have $\beta = k_m y_0 a/k_B T$, $\kappa = k_m h_0(h_0 - d_m)/k_B T$ and using a typical crossbridge stiffness $k_m \sim 1 - 2$ pN/nm, power-stroke size $y_0 \sim 8 - 10$ nm, motor head size $d_m \sim 12 - 14$ nm, lever arm shift $a \sim 1$ nm, thick-thin interfilament spacing $h_0 \sim 15 - 23$ nm, and $k_B T = 4.14$ pN nm (300 K) [S33, S39, S41, S128] we get $\kappa \sim \mathcal{O}(1 - 100)$ and $\beta \sim \mathcal{O}(1 - 10)$. The parameter values we obtain by fitting for $\kappa$ and $\beta$ (Table II) are reasonably consistent with these highly simplified estimates. On the other hand, the kinetic time scale $\tau_k$ for *Drosophila* flight muscle and rabbit skeletal muscle (not mouse cardiac muscle) is substantially different from previous measurements both in-vitro and in reconstituted preparations (see Table I and references therein). It is currently unclear what the source of this discrepancy is, it may be due to our neglect of the low frequency power law response (previous *ad hoc* fits that include additional dissipative and power law terms in the response do recover a more consistent estimate of $\tau_k$ [S78, S110]), or that our two state kinetic description is too oversimplified, or fluid flow may also be relevant for high enough frequencies, despite the permeable boundary, or perhaps (most likely) a combination of all these effects. Note, we face no such issue with the fit for murine cardiac muscle, where the kinetic time scale obtained by fitting is consistent with previous independent measurements.

##### D. Silhouette image credits

The animal silhouettes used in this paper were adapted from the online open database PhyloPic. Individual image credits are listed below:

- Hawkmoth: Gareth Monger, license
- Fruitfly: Ramiro Morales-Hojashttp
- Rabbit: Sarah Werning, license
- Mouse: Jiro Wada
- Oyster toadfish: No copyright, uncredited
- Rattlesnake: Blair Perry
- Cicada: Christoph Schomburg
- Giant waterbug: Karina Garcia, license
- Bumble bee: Ferran Sayol
- Hummingbird: Margot Michaud
- Mosquito: Richard J. Harris

[S1] M Cristina Marchetti, Jean-François Joanny, Sriram Ramaswamy, Tanniemola B Liverpool, Jacques Prost, Madan Rao, and R Aditi Simha. Hydrodynamics of soft active matter. *Reviews of Modern Physics*, 85(3):1143, 2013.

[S2] Jacques Prost, Frank Jülicher, and Jean-François Joanny. Active gel physics. *Nature physics*, 11(2):111–117, 2015.

[S3] Andrew C Callan-Jones and Frank Jülicher. Hydrodynamics of active permeating gels. *New Journal of Physics*, 13(9): 093027, 2011.

[S4] Shiladitya Banerjee, Tanniemola B Liverpool, and M Cristina Marchetti. Generic phases of cross-linked active gels: relaxation, oscillation and contractility. *EPL (Europhysics Letters)*, 96(5):58004, 2011.

[S5] Shiladitya Banerjee and M Cristina Marchetti. Instabilities and oscillations in isotropic active gels. *Soft Matter*, 7(2): 463–473, 2011.

[S6] Stefan Günther and Karsten Kruse. Spontaneous waves in muscle fibres. *new Journal of physics*, 9(11):417, 2007.

[S7] Ananyo Maitra and Sriram Ramaswamy. Oriented active solids. *Physical review letters*, 123(23):238001, 2019.

[S8] Tanniemola B Liverpool and M Cristina Marchetti. Bridging the microscopic and the hydrodynamic in active filament solutions. *EPL (Europhysics Letters)*, 69(5):846, 2005.

[S9] Aphrodite Ahmadi, M Cristina Marchetti, and Tanniemola B Liverpool. Hydrodynamics of isotropic and liquid crystalline active polymer solutions. *Physical Review E*, 74(6):061913, 2006.

[S10] David Saintillan and Michael J Shelley. Active suspensions and their nonlinear models. *Comptes Rendus Physique*, 14 (6):497–517, 2013.

[S11] Sebastian Fürthauer, Bezia Lemma, Peter J Foster, Stephanie C Ems-McClung, Che-Hang Yu, Claire E Walczak, Zvonimir Dogic, Daniel J Needleman, and Michael J Shelley. Self-straining of actively crosslinked microtubule networks. *Nature physics*, 15(12):1295–1300, 2019.

[S12] Sebastian Fürthauer, Daniel J Needleman, and Michael J Shelley. A design framework for actively crosslinked filament networks. *New Journal of Physics*, 23(1):013012, 2021.

[S13] Sebastian Fürthauer and Michael J Shelley. How cross-link numbers shape the large-scale physics of cytoskeletal materials. *Annual Review of Condensed Matter Physics*, 13:365–384, 2022.

[S14] Mark Warner and Eugene Michael Terentjev. *Liquid crystal elastomers*, volume 120. Oxford university press, 2007.

[S15] Paul Dalhaimer, Dennis E Discher, and Tom C Lubensky. Crosslinked actin networks show liquid crystal elastomer behaviour, including soft-mode elasticity. *Nature Physics*, 3(5):354–360, 2007.

[S16] Lev Davidovich Landau, Evgenij M Lifšic, Evegnii Mikhailovich Lifshitz, Arnold Markovich Kosevich, and Lev Petrovich Pitaevskii. *Theory of elasticity: volume 7*, volume 7. Elsevier, 1986.

[S17] RC Rossi. Prediction of the elastic moduli of composites. *Journal of the American Ceramic Society*, 51(8):433–440, 1968.

[S18] YH Zhao, GP Tandon, and GJ Weng. Elastic moduli for a class of porous materials. *Acta Mechanica*, 76(1):105–131, 1989.

[S19] Gp P Tandon and Gj J Weng. The effect of aspect ratio of inclusions on the elastic properties of unidirectionally aligned composites. *Polymer composites*, 5(4):327–333, 1984.

[S20] Maurice A Biot. General theory of three-dimensional consolidation. *Journal of applied physics*, 12(2):155–164, 1941.

[S21] Joseph B Keller. Viscous flow through a grating or lattice of cylinders. *Journal of fluid mechanics*, 18(1):94–96, 1964.

[S22] B Rikard Gebart. Permeability of unidirectional reinforcements for rtm. *Journal of composite materials*, 26(8):1100–1133, 1992.

[S23] John Happel. Viscous flow relative to arrays of cylinders. *AIChE Journal*, 5(2):174–177, 1959.

[S24] Graham W Jackson and David F James. The permeability of fibrous porous media. *The Canadian Journal of Chemical Engineering*, 64(3):364–374, 1986.

[S25] Ashok S Sangani and C Yao. Transport processes in random arrays of cylinders. ii. viscous flow. *The Physics of fluids*, 31(9):2435–2444, 1988.

[S26] Barry M Millman. The filament lattice of striated muscle. *Physiological reviews*, 78(2):359–391, 1998.

[S27] RW Lymn and Edwin W Taylor. Mechanism of adenosine triphosphate hydrolysis by actomyosin. *Biochemistry*, 10(25): 4617–4624, 1971.

[S28] Albert M Gordon, Earl Homsher, and Mike Regnier. Regulation of contraction in striated muscle. *Physiological reviews*, 80(2):853–924, 2000.

[S29] Robert K Josephson, Jean G Malamud, and Darrell R Stokes. Asynchronous muscle: a primer. *Journal of Experimental Biology*, 203(18):2713–2722, 2000.

[S30] Frank Jülicher and Jacques Prost. Spontaneous oscillations of collective molecular motors. *Physical review letters*, 78 (23):4510, 1997.

[S31] Stephan W Grill, Karsten Kruse, and Frank Jülicher. Theory of mitotic spindle oscillations. *Physical review letters*, 94 (10):108104, 2005.

[S32] Thomas Guérin, Jacques Prost, Pascal Martin, and Jean-François Joanny. Coordination and collective properties of molecular motors: theory. *Current opinion in cell biology*, 22(1):14–20, 2010.

[S33] Thomas Guérin, J Prost, and J-F Joanny. Dynamical behavior of molecular motor assemblies in the rigid and crossbridge models. *The European Physical Journal E*, 34(6):1–21, 2011.

[S34] Andrej Vilfan, Erwin Frey, and Franz Schwabl. Elastically coupled molecular motors. *The European Physical Journal B-Condensed Matter and Complex Systems*, 3(4):535–546, 1998.

[S35] Andrej Vilfan and Erwin Frey. Oscillations in molecular motor assemblies. *Journal of physics: Condensed matter*, 17 (47):S3901, 2005.

[S36] Arvind Gopinath, Raghunath Chelakkot, and L Mahadevan. Effective elasticity and persistence of strain in active filament-motor assemblies. *bioRxiv*, 2021.

[S37] M Schoenberg. Geometrical factors influencing muscle force development. i. the effect of filament spacing upon axial forces. *Biophysical journal*, 30(1):51–67, 1980.

[S38] M Schoenberg. Geometrical factors influencing muscle force development. ii. radial forces. *Biophysical journal*, 30(1): 69–77, 1980.

[S39] Joseph D Powers, Sage A Malingen, Michael Regnier, and Thomas L Daniel. The sliding filament theory since andrew huxley: Multiscale and multidisciplinary muscle research. *Annual Review of Biophysics*, 50:373–400, 2021.

[S40] J Howard. Sinauer associates; sunderland, ma: 2001. *Mechanics of motor proteins and the cytoskeleton.*, 2001.

[S41] Andrej Vilfan and Thomas Duke. Instabilities in the transient response of muscle. *Biophysical Journal*, 85(2):818–827, 2003.

[S42] Marco Linari, Marco Caremani, Claudia Piperio, Philip Brandt, and Vincenzo Lombardi. Stiffness and fraction of myosin motors responsible for active force in permeabilized muscle fibers from rabbit psoas. *Biophysical journal*, 92(7): 2476–2490, 2007.

[S43] Alexandre Lewalle, Walter Steffen, Olivia Stevenson, Zhenqian Ouyang, and John Sleep. Single-molecule measurement of the stiffness of the rigor myosin head. *Biophysical journal*, 94(6):2160–2169, 2008.

[S44] Thomas C Irving, John Konhilas, Darold Perry, Robert Fischetti, and Pieter P De Tombe. Myofilament lattice spacing as a function of sarcomere length in isolated rat myocardium. *American Journal of Physiology-Heart and Circulatory Physiology*, 279(5):H2568–H2573, 2000.

[S45] C David Williams, Michael Regnier, and Thomas L Daniel. Axial and radial forces of cross-bridges depend on lattice spacing. *PLoS computational biology*, 6(12):e1001018, 2010.

[S46] C David Williams, Mary K Salcedo, Thomas C Irving, Michael Regnier, and Thomas L Daniel. The length–tension curve in muscle depends on lattice spacing. *Proceedings of the Royal Society B: Biological Sciences*, 280(1766):20130697, 2013.

[S47] Younss Ait-Mou, Karen Hsu, Gerrie P Farman, Mohit Kumar, Marion L Greaser, Thomas C Irving, and Pieter P de Tombe. Titin strain contributes to the frank–starling law of the heart by structural rearrangements of both thin-and thick-filament proteins. *Proceedings of the National Academy of Sciences*, 113(8):2306–2311, 2016.

[S48] Claudia Veigel, Justin E Molloy, Stephan Schmitz, and John Kendrick-Jones. Load-dependent kinetics of force production by smooth muscle myosin measured with optical tweezers. *Nature cell biology*, 5(11):980–986, 2003.

[S49] Bin Guo and William H Guilford. Mechanics of actomyosin bonds in different nucleotide states are tuned to muscle contraction. *Proceedings of the National Academy of Sciences*, 103(26):9844–9849, 2006.

[S50] TAJ Duke. Molecular model of muscle contraction. *Proceedings of the National Academy of Sciences*, 96(6):2770–2775, 1999.

[S51] Gabriella Piazzesi, Massimo Reconditi, Marco Linari, Leonardo Lucii, Pasquale Bianco, Elisabetta Brunello, Valérie Decostre, Alex Stewart, David B Gore, Thomas C Irving, et al. Skeletal muscle performance determined by modulation of number of myosin motors rather than motor force or stroke size. *Cell*, 131(4):784–795, 2007.

[S52] John William Sutton Pringle. The croonian lecture, 1977-stretch activation of muscle: function and mechanism. *Proceedings of the Royal Society of London. Series B. Biological Sciences*, 201(1143):107–130, 1978.

[S53] R Aditi Simha and Sriram Ramaswamy. Hydrodynamic fluctuations and instabilities in ordered suspensions of self-propelled particles. *Physical review letters*, 89(5):058101, 2002.

[S54] TC Irving and DW Maughan. In vivo x-ray diffraction of indirect flight muscle from drosophila melanogaster. *Biophysical journal*, 78(5):2511–2515, 2000.

[S55] Qazi Ibadur Rahman, Gerhard Schmeisser, et al. *Analytic theory of polynomials.* Number 26 in London mathematical society monographs. Oxford University Press, 2002.

[S56] Douglas M Swank, Vivek K Vishnudas, and David W Maughan. An exceptionally fast actomyosin reaction powers insect flight muscle. *Proceedings of the National Academy of Sciences*, 103(46):17543–17547, 2006.

[S57] Colin Scheibner, Anton Souslov, Debarghya Banerjee, Piotr Surówka, William TM Irvine, and Vincenzo Vitelli. Odd elasticity. *Nature Physics*, 16(4):475–480, 2020.

[S58] Tzer Han Tan, Alexander Mietke, Junang Li, Yuchao Chen, Hugh Higinbotham, Peter J Foster, Shreyas Gokhale, Jörn Dunkel, and Nikta Fakhri. Odd dynamics of living chiral crystals. *Nature*, 607(7918):287–293, 2022.

[S59] Bradley M Palmer. A strain-dependency of myosin off-rate must be sensitive to frequency to predict the b-process of sinusoidal analysis. In *Muscle Biophysics*, pages 57–75. Springer, 2010.

[S60] Maurice A Biot. Xliii. non-linear theory of elasticity and the linearized case for a body under initial stress. *The London, Edinburgh, and Dublin Philosophical Magazine and Journal of Science*, 27(183):468–489, 1939.

[S61] Maurice A Biot. *Mechanics of incremental deformations.* John Wiley & Sons, 1965.

[S62] C Truesdell. The meaning of betti's reciprocal theorem. *J. Research of NBS B*, 67:85–86, 1963.

[S63] JE Avron. Odd viscosity. *Journal of statistical physics*, 92(3):543–557, 1998.

[S64] Lawrence C Rome, Douglas A Syme, Stephen Hollingworth, Stan L Lindstedt, and Stephen M Baylor. The whistle and the rattle: the design of sound producing muscles. *Proceedings of the National Academy of Sciences*, 93(15):8095–8100, 1996.

[S65] Lawrence C Rome. Design and function of superfast muscles: new insights into the physiology of skeletal muscle. *Annu. Rev. Physiol.*, 68:193–221, 2006.

[S66] Michael L Fine. Seasonal and geographical variation of the mating call of the oyster toadfish opsanus tau l. *Oecologia*, 36(1):45–57, 1978.

[S67] Michael L Fine, Noelle M Burns, and Thomas M Harris. Ontogeny and sexual dimorphism of sonic muscle in the oyster toadfish. *Canadian Journal of Zoology*, 68(7):1374–1381, 1990.

[S68] Lawrence C Rome and Stan L Lindstedt. The quest for speed: muscles built for high-frequency contractions. *Physiology*, 13(6):261–268, 1998.

[S69] Wai Pang Chan and Michael H Dickinson. In vivo length oscillations of indirect flight muscles in the fruit fly drosophila virilis. *The Journal of experimental biology*, 199(12):2767–2774, 1996.

[S70] Robert K Josephson and David Young. Synchronous and asynchronous muscles in cicadas. *Journal of Experimental Biology*, 91(1):219–237, 1981.

[S71] David Young and Robert K Josephson. 100 hz is not the upper limit of synchronous muscle contraction. *Nature*, 309 (5965):286–287, 1984.

[S72] Robert K Josephson and David Young. A synchronous insect muscle with an operating frequency greater than 500 hz. *Journal of Experimental Biology*, 118(1):185–208, 1985.

[S73] Ken-ichi OGAWA and Tozo KANDA. Wingbeat frequencies of some anopheline mosquitoes of east asia (diptera: Culicidae). *Applied entomology and zoology*, 21(3):430–435, 1986.

[S74] Robert Dudley. *The biomechanics of insect flight: form, function, evolution.* Princeton University Press, 2002.

[S75] Lawrence C Rome, Chris Cook, Douglas A Syme, Martin A Connaughton, Miriam Ashley-Ross, Andrei Klimov, Boris Tikunov, and Yale E Goldman. Trading force for speed: why superfast crossbridge kinetics leads to superlow forces. *Proceedings of the National Academy of Sciences*, 96(10):5826–5831, 1999.

[S76] James H Martin and Roland M Bagby. Properties of rattlesnake shaker muscle. *Journal of Experimental Zoology*, 185 (3):293–300, 1973.

[S77] Robert K Josephson and David Young. Fiber ultrastructure and contraction kinetics in insect fast muscles. *American zoologist*, 27(4):991–1000, 1987.

[S78] Bertrand CW Tanner, Gerrie P Farman, Thomas C Irving, David W Maughan, Bradley M Palmer, and Mark S Miller. Thick-to-thin filament surface distance modulates cross-bridge kinetics in drosophila flight muscle. *Biophysical journal*, 103(6):1275–1284, 2012.

[S79] DJ Aidley and DCS White. Mechanical properties of glycerinated fibres from the tymbal muscles of a brazilian cicada. *The Journal of Physiology*, 205(1):179, 1969.

[S80] Thomas Burgoyne, John M Heumann, Edward P Morris, Carlo Knupp, Jun Liu, Michael K Reedy, Kenneth A Taylor, Kuan Wang, and Pradeep K Luther. Three-dimensional structure of the basketweave z-band in midshipman fish sonic muscle. *Proceedings of the National Academy of Sciences*, 116(31):15534–15539, 2019. Assuming similar filament spacing across toadfish sonic muscles.

[S81] AW Clark and E Schultz. Rattlesnake shaker muscle: Ii. fine structure. *Tissue and Cell*, 12(2):335–351, 1980.

[S82] Hiroyuki Iwamoto. The tymbal muscle of cicada has flight muscle-type sarcomeric architecture and protein expression. *Zoological Letters*, 3(1):1–10, 2017.

[S83] Hiroyuki Iwamoto. X-ray diffraction pattern from the flight muscle of toxorhynchites towadensis reveals the specific phylogenic position of mosquito among diptera. *Zoological letters*, 1(1):1–7, 2015.

[S84] Michael L Fine, Barbara Bernard, and Thomas M Harris. Functional morphology of toadfish sonic muscle fibers: relationship to possible fiber division. *Canadian Journal of Zoology*, 71(11):2262–2274, 1993.

[S85] E Schultz, AW Clark, A Suzuki, and RG Cassens. Rattlesnake shaker muscle: I. a light microscopic and histochemical study. *Tissue and Cell*, 12(2):323–334, 1980.

[S86] Douglas M Swank. Mechanical analysis of drosophila indirect flight and jump muscles. *Methods*, 56(1):69–77, 2012.
[S87] Patrick C Nahirney, Jeffrey G Forbes, H Douglas Morris, Susanne C Chock, and Kuan Wang. What the buzz was all about: superfast song muscles rattle the tymbals of male periodical cicadas. *The FASEB journal*, 20(12):2017–2026, 2006.
[S88] Alvin R Hylton. Histopathological changes with age in the flight muscles of the african mosquito, eretmapodites chrysogaster. *Journal of Invertebrate Pathology*, 8(1):75–78, 1966.
[S89] Jonathan E Hirsch, John W Bigbee, and Michael L Fine. Continuous adult development of multiple innervation in toadfish sonic muscle. *Journal of neurobiology*, 36(3):348–356, 1998.
[S90] HC Bennet-Clark and AG Daws. Transduction of mechanical energy into sound energy in the cicada cyclochila australasiae. *Journal of Experimental Biology*, 202(13):1803–1817, 1999.
[S91] Bruce G Johnson Jr and Wayne A Rowley. Age-related ultrastructural changes in the flight muscle of the mosquito, culex tarsalis. *Journal of insect physiology*, 18(12):2375–2389, 1972.
[S92] Antonio Celestino-Montes, Salvador Hernández-Martínez, Mario Henry Rodríguez, Febe Elena Cázares-Raga, Carlos Vázquez-Calzada, Anel Lagunes-Guillén, Bibiana Chávez-Munguía, José Ángel Rubio-Miranda, Felipe de Jesús Hernández-Cázares, Leticia Cortés-Martínez, et al. Development of the indirect flight muscles of aedes aegypti, a main arbovirus vector. *BMC developmental biology*, 21(1):1–15, 2021.
[S93] Hiroyuki Iwamoto. Structure, function and evolution of insect flight muscle. *Biophysics*, 7:21–28, 2011.
[S94] Justin Edward Molloy. *Active insect fibrillar flight muscle, its mechanical performance and cross-bridge kinetics*. PhD thesis, University of York, 1988.
[S95] H Iwamoto and N Yagi. The molecular trigger for high-speed wing beats in a bee. *Science*, 341(6151):1243–1246, 2013.
[S96] Takumi Washio, Seine A Shintani, Hideo Higuchi, Seiryo Sugiura, and Toshiaki Hisada. Effect of myofibril passive elastic properties on the mechanical communication between motor proteins on adjacent sarcomeres. *Scientific reports*, 9(1): 1–17, 2019.
[S97] Fumiaki Kono, Seitaro Kawai, Yuta Shimamoto, and Shin'ichi Ishiwata. Nanoscopic changes in the lattice structure of striated muscle sarcomeres involved in the mechanism of spontaneous oscillatory contraction (spoc). *Scientific reports*, 10(1):1–14, 2020.
[S98] Crawford H Greenewalt. Dimensional relationships for flying animals. *Smithsonian miscellaneous collections*, 1962.
[S99] Heidi L Lujan and Stephen E DiCarlo. Cardiac output, at rest and during exercise, before and during myocardial ischemia, reperfusion, and infarction in conscious mice. *American Journal of Physiology-Regulatory, Integrative and Comparative Physiology*, 304(4):R286–R295, 2013.
[S100] Paul ML Janssen, Brandon J Biesiadecki, Mark T Ziolo, and Jonathan P Davis. The need for speed: mice, men, and myocardial kinetic reserve. *Circulation research*, 119(3):418–421, 2016.
[S101] Bernadette M Glasheen, Catherine C Eldred, Leah C Sullivan, Cuiping Zhao, Michael K Reedy, Robert J Edwards, and Douglas M Swank. Stretch activation properties of drosophila and lethocerus indirect flight muscle suggest similar calcium-dependent mechanisms. *American Journal of Physiology-Cell Physiology*, 313(6):C621–C631, 2017.
[S102] Hiroyuki Iwamoto. The earliest molecular response to stretch of insect flight muscle as revealed by fast x-ray diffraction recording. *Scientific reports*, 7(1):1–10, 2017.
[S103] KM Gilmour and CP Ellington. Power output of glycerinated bumblebee flight muscle. *Journal of experimental biology*, 183(1):77–100, 1993.
[S104] Stefan Galler, Brant Gang Wang, and Masataka Kawai. Elementary steps of the cross-bridge cycle in fast-twitch fiber types from rabbit skeletal muscles. *Biophysical Journal*, 89(5):3248–3260, 2005.
[S105] Susumu Hagiwara, Shiko Chichibu, and Norman Simpson. Neuromuscular mechanisms of wing beat in hummingbirds. *Zeitschrift für vergleichende Physiologie*, 60(2):209–218, 1968.
[S106] Bradley M Palmer, Takeki Suzuki, Yuan Wang, William D Barnes, Mark S Miller, and David W Maughan. Two-state model of acto-myosin attachment-detachment predicts c-process of sinusoidal analysis. *Biophysical journal*, 93(3): 760–769, 2007.
[S107] MGA Wilson and DCS White. The role of magnesium adenosine triphosphate in the contractile kinetics of insect fibrillar flight muscle. *Journal of Muscle Research & Cell Motility*, 4(3):283–306, 1983.
[S108] HL Granzier and Kuan Wang. Interplay between passive tension and strong and weak binding cross-bridges in insect indirect flight muscle. a functional dissection by gelsolin-mediated thin filament removal. *The Journal of general physiology*, 101(2):235–270, 1993.
[S109] R Josephson and C Ellington. Power output from a flight muscle of the bumblebee bombus terrestris. i. some features of the dorso-ventral flight muscle. *The Journal of experimental biology*, 200(8):1215–1226, 1997.
[S110] Masataka Kawai and Philip W Brandt. Sinusoidal analysis: a high resolution method for correlating biochemical reactions with physiological processes in activated skeletal muscles of rabbit, frog and crayfish. *Journal of Muscle Research & Cell Motility*, 1(3):279–303, 1980.
[S111] Peter J Reiser, Kenneth C Welch Jr, Raul K Suarez, and Douglas L Altshuler. Very low force-generating ability and unusually high temperature dependency in hummingbird flight muscle fibers. *Journal of Experimental Biology*, 216(12): 2247–2256, 2013.
[S112] I Matsubara, DW Maughan, Y Saeki, and N Yagi. Cross-bridge movement in rat cardiac muscle as a function of calcium concentration. *The Journal of Physiology*, 417(1):555–565, 1989.
[S113] Masataka Kawai, John S Wray, and Yan Zhao. The effect of lattice spacing change on cross-bridge kinetics in chemically skinned rabbit psoas muscle fibers. i. proportionality between the lattice spacing and the fiber width. *Biophysical journal*,

64(1):187–196, 1993.

[S114] P Bryant Chase, Todd M Denkinger, and Martin J Kushmerick. Effect of viscosity on mechanics of single, skinned fibers from rabbit psoas muscle. *Biophysical journal*, 74(3):1428–1438, 1998.

[S115] O Mathieu-Costello, RK Suarez, and PW Hochachka. Capillary-to-fiber geometry and mitochondrial density in hummingbird flight muscle. *Respiration physiology*, 89(1):113–132, 1992.

[S116] Thomas F Robinson, Barbara S Hayward, John W Krueger, Edmund H Sonnenblick, and Beatrice A Wittenberg. Isolated heart myocytes: ultrastructural case study technique. *Journal of Microscopy*, 124(2):135–142, 1981.

[S117] Marco Linari, Michael K Reedy, Mary C Reedy, Vincenzo Lombardi, and Gabriella Piazzesi. Ca-activation and stretch-activation in insect flight muscle. *Biophysical Journal*, 87(2):1101–1111, 2004.

[S118] KM Gilmour and CP Ellington. In vivo muscle length changes in bumblebees and the in vitro effects on work and power. *Journal of experimental biology*, 183(1):101–113, 1993.

[S119] Richard L Lieber and Field T Blevins. Skeletal muscle architecture of the rabbit hindlimb: functional implications of muscle design. *Journal of morphology*, 199(1):93–101, 1989.

[S120] Ramesh Vemuri, Edward B Lankford, Karl Poetter, Shahin Hassanzadeh, Kazuyo Takeda, Zu-Xi Yu, Victor J Ferrans, and Neal D Epstein. The stretch-activation response may be critical to the proper functioning of the mammalian heart. *Proceedings of the National Academy of Sciences*, 96(3):1048–1053, 1999.

[S121] Michael S Tu and Thomas L Daniel. Cardiac-like behavior of an insect flight muscle. *Journal of Experimental Biology*, 207(14):2455–2464, 2004.

[S122] Ankit Rohatgi. Webplotdigitizer: Version 4.5, 2021. URL https://automeris.io/WebPlotDigitizer.

[S123] Michael Dickinson, Gerrie Farman, Mark Frye, Tanya Bekyarova, David Gore, David Maughan, and Thomas Irving. Molecular dynamics of cyclically contracting insect flight muscle in vivo. *Nature*, 433(7023):330–334, 2005.

[S124] Julie A Cass, C Dave Williams, Tom C Irving, Eric Lauga, Sage Malingen, Thomas L Daniel, and Simon N Sponberg. A mechanism for sarcomere breathing: volume change and advective flow within the myofilament lattice. *Biophysical Journal*, 2021.

[S125] Sage A Malingen, Anthony M Asencio, Julie A Cass, Weikang Ma, Thomas C Irving, and Thomas L Daniel. In vivo x-ray diffraction and simultaneous emg reveal the time course of myofilament lattice dilation and filament stretch. *Journal of Experimental Biology*, 223(17):jeb224188, 2020.

[S126] Joshua J Waterfall, Fergal P Casey, Ryan N Gutenkunst, Kevin S Brown, Christopher R Myers, Piet W Brouwer, Veit Elser, and James P Sethna. Sloppy-model universality class and the vandermonde matrix. *Physical review letters*, 97 (15):150601, 2006.

[S127] Ryan N Gutenkunst, Joshua J Waterfall, Fergal P Casey, Kevin S Brown, Christopher R Myers, and James P Sethna. Universally sloppy parameter sensitivities in systems biology models. *PLoS computational biology*, 3(10):e189, 2007.

[S128] Andrew F Huxley and Ro M Simmons. Proposed mechanism of force generation in striated muscle. *Nature*, 233(5321): 533–538, 1971.

[S129] Hongjun Liu, Mark S Miller, Douglas M Swank, William A Kronert, David W Maughan, and Sanford I Bernstein. Paramyosin phosphorylation site disruption affects indirect flight muscle stiffness and power generation in drosophila melanogaster. *Proceedings of the National Academy of Sciences*, 102(30):10522–10527, 2005.

[S130] Michael H Dickinson, Christopher J Hyatt, Fritz-Olaf Lehmann, Jeffrey R Moore, Mary C Reedy, Amanda Simcox, Rutawain Tohtong, Jim O Vigoreaux, Hiroshi Yamashita, and David W Maughan. Phosphorylation-dependent power output of transgenic flies: an integrated study. *Biophysical Journal*, 73(6):3122–3134, 1997.

[S131] Nicholas MP King, Methajit Methawasin, Joshua Nedrud, Nicholas Harrell, Charles S Chung, Michiel Helmes, and Henk Granzier. Mouse intact cardiac myocyte mechanics: cross-bridge and titin-based stress in unactivated cells. *Journal of General Physiology*, 137(1):81–91, 2011.

[S132] HL Granzier and Kuan Wang. Passive tension and stiffness of vertebrate skeletal and insect flight muscles: the contribution of weak cross-bridges and elastic filaments. *Biophysical journal*, 65(5):2141–2159, 1993.

[S133] Guido Van Rossum and Fred L. Drake. *Python 3 Reference Manual*. CreateSpace, Scotts Valley, CA, 2009. ISBN 1441412697.

[S134] Pauli Virtanen, Ralf Gommers, Travis E. Oliphant, Matt Haberland, Tyler Reddy, David Cournapeau, Evgeni Burovski, Pearu Peterson, Warren Weckesser, Jonathan Bright, Stéfan J. van der Walt, Matthew Brett, Joshua Wilson, K. Jarrod Millman, Nikolay Mayorov, Andrew R. J. Nelson, Eric Jones, Robert Kern, Eric Larson, C J Carey, İlhan Polat, Yu Feng, Eric W. Moore, Jake VanderPlas, Denis Laxalde, Josef Perktold, Robert Cimrman, Ian Henriksen, E. A. Quintero, Charles R. Harris, Anne M. Archibald, Antônio H. Ribeiro, Fabian Pedregosa, Paul van Mulbregt, and SciPy 1.0 Contributors. SciPy 1.0: Fundamental Algorithms for Scientific Computing in Python. *Nature Methods*, 17:261–272, 2020. doi:10.1038/s41592-019-0686-2.

[S135] Wolfram Research, Inc. Mathematica, Version 13.0.0. URL https://www.wolfram.com/mathematica. Champaign, IL, 2021.
